# Microtubule stiffening by the doublecortin-domain protein ZYG-8 contributes to mitotic spindle orientation during zygote division in *Caenorhabditis elegans*

**DOI:** 10.1101/2024.11.29.624795

**Authors:** Louis Cueff, Loïc Schmitt, Ewen Huet, Sylvain Pastezeur, Méline Coquil, Talia Savary, Anouk Sénard, Jacques Pécréaux, Hélène Bouvrais

**Affiliations:** CNRS, Univ Rennes, IGDR (Institut de Génétique et Développement de Rennes) – UMR 6290, F-35000 Rennes, France

## Abstract

In the *Caenorhabditis elegans* zygote, mutations in *zyg-8^DCLK1^*, the sole Doublecortin-family member, disrupt mitotic spindle positioning, as seen by immunofluorescence. Doublecortin proteins bind microtubules and are thought to stabilise or rigidify them. In the zygote, ZYG-8 only modestly affects microtubule growth and nucleation. We thus investigated whether these moderate dynamic perturbations alone could explain the spindle mispositioning observed in *zyg-8* mutants.

Using three complementary genetic perturbations—RNAi-mediated depletion of ZYG-8, its overexpression, and the thermosensitive *zyg-8(or484ts)* mutant (that disrupts microtubule binding)—we observed altered spindle pole oscillations and changes in microtubule cortical-contact behaviour, indicative of impaired cortical forces. Importantly, these phenotypes could not be fully explained by previously reported alterations in microtubule dynamics, suggesting an additional mechanism. Our findings indicate that ZYG-8 increases microtubule rigidity: ZYG-8 depletion or mutation led to more frequent microtubule bending and higher curvature and tortuosity. Simulations confirmed that reduced rigidity prolongs cortical contact lifetimes, an effect we experimentally observed in *zyg-8(RNAi)* embryos.

Using custom biophysical assays, we showed that microtubule softening in *zyg-8(RNAi)* embryos and *zyg-8* mutants reduced the efficiency of centring forces, leading to exaggerated spindle-pole oscillations. In mutants, the largest oscillations caused spindle poles to move closer to the cell periphery, preventing re-centring and resulting in spindle mispositioning and misorientation during late anaphase. Importantly, reducing cortical pulling forces rescued orientation defects, highlighting the importance of balanced pulling-pushing forces for proper spindle positioning.

We propose that sufficient microtubule rigidity is essential for generating effective cortical pushing forces, potentially in synergy with other microtubule properties, which contribute to centring mechanisms that ensure accurate spindle orientation in late mitosis. Given that DCLK1 is frequently deregulated in human cancers and that accurate spindle positioning is essential for maintaining cell proliferation-differentiation balance, these findings may have implications for understanding how disruptions in microtubule mechanics contribute to carcinogenesis.

**Author summary:** During cell division, the mitotic spindle—a structure made of microtubules that separates chromosomes in daughter cells—must position itself precisely to ensure healthy cell growth and differentiation. Microtubules are tiny, dynamic, semi-flexible fibers that notably generate forces at the cell periphery, cortical pulling and pushing. These two opposite forces contribute to position the spindle. In the worm *C. elegans*, the protein ZYG-8 (similar to human DCLK1) stabilizes the microtubules. Our study reveals that ZYG-8 also makes microtubules stiffer, a property critical for generating strong enough pushing forces. When *zyg-8* was impaired (mutation or RNAi), microtubules bent excessively, weakening the pushing forces that keep the spindle correctly centered and oriented. As a result, softer microtubules in *zyg-8* mutants caused the spindle to drift toward the cell’s periphery, resulting in mispositioning and misorientation when mitosis ends. By reducing the opposing cortical pulling forces in *zyg-8* mutants, we restored normal spindle orientation, highlighting that microtubule rigidity helps balance the forces that position the spindle correctly. Our findings suggest that microtubule stiffness is essential for accurate cell division—a process vital for both normal development and disease prevention, including cancers where homologous proteins are often disrupted.

## INTRODUCTION

Microtubules are semi-rigid polymers that form a crucial component of the cell cytoskeleton alongside actin and intermediate filaments. In coordination with molecular motors and microtubule-associated proteins (MAPs), microtubules perform essential functions, including cell shaping [1, 2], intracellular trafficking [3], and cell migration [4]. Microtubules also play major roles in cell division; for example, spindle microtubules assemble into the mitotic spindle and segregate chromosomes, while astral microtubules help position the spindle [5–11]. Along a structural line, each microtubule comprises protofilaments that are linear chains of tubulin dimers added or removed dynamically. Microtubule dynamic instability, characterised by stochastic transitions between growing and shrinking phases, is regulated *in vivo* by numerous MAPs [12–14]. The mechanical properties of microtubules, particularly their flexural rigidity—i.e., their resistance to bending—have received comparatively less attention. Notably, microtubules exhibit the highest flexural rigidity among cytoskeletal filaments [15]. While *in vitro* reconstituted microtubules are bent at the scale of a few millimetres [15–18], *in vivo* observations revealed that microtubules appear bent at the micrometre scale [19–21]. On one hand, high microtubule rigidity is necessary for efficient intracellular trafficking [22], for generating pushing forces against the cell periphery that contribute to spindle or nucleus positioning [23–30], or for exerting pushing forces on chromosome arms to produce the polar ejection force with the assistance of chromokinesins [31, 32]. On the other hand, microtubules must also be flexible enough to bend, for example when growing through the crowded cytoplasm [33]. This highlights the need to regulate microtubule flexural rigidity *in vivo* to support its various biological functions [21].

Research across various biological systems has identified three major protein families involved in regulating microtubule rigidity: the Doublecortin family (including DCX and DCLK1), the MAP2/MAP4/Tau superfamily, and the SPD-1/Ase1/PRC1/MAP65 family [16, 18, 34–36]. While the latter family plays crucial roles in non-neuronal contexts, the first two have been primarily studied in neuronal systems. For instance, the simultaneous knockdown of DCX and DCLK1 in rat neurons led to more curved microtubules at the growing axon tip [37]. Doublecortin-family proteins associate with microtubules through their doublecortin domains. Notably, these proteins bind between the microtubule protofilaments at the vertices of four tubulin dimers, contributing to the lateral coupling of adjacent protofilaments [38, 39]. This binding likely restricts protofilament sliding, thereby increasing microtubule rigidity [40, 41]. Along a similar line, Tau-deficient axons have been shown to exhibit curled microtubules [42]. Tau family proteins bind longitudinally along the outer ridges of microtubule protofilaments. It would strengthen the longitudinal contacts between tubulin heterodimers within a protofilament, straightening the protofilaments [34, 43, 44]. In contrast to Doublecortin and Tau proteins, MAP65/Ase1, by binding to individual microtubules and microtubule bundles, decreases their flexural rigidity [36]. Upon crossing-linking antiparallel microtubules, proteins of this family favour the formation and stabilisation of microtubule bundles [36, 45–47]. Importantly, unlike the SPD-1/Ase1/PRC1/MAP65 family, both the Doublecortin and Tau families can also regulate microtubule dynamics, making it challenging to disentangle their roles in mechanical versus dynamic regulation. Some *in vivo* studies indicate that these proteins stabilise microtubules and/or promote their growth [42, 48]. They may reduce their dynamics by decreasing catastrophe frequency, depolymerisation and nucleation rates [48–50]. While the impacts of controlling microtubule rigidity have primarily been studied in neurons, microtubule-rigidity contribution during cell division also warrants investigations.

Previous studies on cell division involving the targeting of proteins from the Doublecortin, Tau, and PRC1 families revealed several defects. These included mitotic spindle mispositioning, abnormalities in spindle formation and integrity, and impaired cell cycle progression [51–60]. During neurogenesis, altered expression of Doublecortin-family proteins led to incorrect cell-fate determination for neural progenitors [51, 52]. Additionally, the deregulation of Tau (whether through mutation or overexpression) resulted in mitotic defects, such as issues with spindle formation, aneuploidy and chromosome misalignment [61, 62]. Proteins from the PRC1 family were also suggested to regulate mitotic spindle mechanics [45, 47]. Thus, targeting proteins from these three families can lead to a wide range of meiotic and mitotic disturbances, although the underlying mechanisms remain unclear. Proposed explanations include perturbations in microtubule stability and in molecular motor activity or recruitment [51–56, 63]. Whether the underlying mechanism could involve microtubule rigidity is yet to be investigated.

In the *Caenorhabditis elegans* zygote, the pronuclei-centrosome complex (NCC) and the mitotic spindle exhibit stereotyped movements during cell division [64]. It makes this system a well-established model for studying asymmetric divisions. In these divisions, the correct positioning and orientation of the spindle are critical for cell fate determination and, more broadly, for the organism’s development [9]. In further detail, the NCC forms at the zygote posterior pole, migrates anteriorly, and the centrosomes align with the anteroposterior axis during prophase. The mitotic spindle assembles near the cell centre after the nuclear envelop breakdown (NEBD), remains centred during metaphase, and is then positioned posteriorly, mainly during anaphase [65]. During this posterior displacement, the spindle undergoes transverse oscillations driven by strong cortical pulling forces [66]. While not essential for asymmetric division, these oscillations have been instrumental in studying spindle positioning forces [67–72]. Astral microtubules, which emanate from the centrosomes, play a central role in spindle movements. These microtubules can transmit or generate three types of forces [5, 7, 73]: first, cortical pulling forces mediated by dynein motors anchored at the cell cortex interacting with shrinking microtubules [67, 74, 75]; second, cytoplasmic pulling forces that are reaction forces generated by organelle movement along microtubules, driven by dynein motors [76]; and third, cortical pushing forces generated by microtubules that grow against the cell cortex [25, 27, 77]. Various models have addressed the roles of these forces. Kimura *et al.* proposed that cytoplasmic pulling forces contribute to NCC centration [76], while others have suggested that these forces do not aid in maintaining spindle position at the cell centre during metaphase [25] or in driving anaphase spindle oscillations [68]. Cortical pulling forces, which are stronger at the posterior, are well-established drivers of posterior spindle displacement and anaphase oscillations [67–69, 75, 78, 79]. However, a restoring force must exist during spindle-pole oscillations, notably to stabilise the spindle position against these strong cortical pulling forces and permit oscillations. This centring mechanism could arise from the embryo’s geometry [68, 75] or microtubule dynamics [67]. Alternative mechanisms include cytoplasmic pulling forces or cortical pushing forces. The latter may result from astral microtubules pushing against the cortex or from microtubule buckling [27, 66, 67]. We and others suggested that microtubule buckling—bending under compressive load—contributes to centring the spindle, in particular for spindle maintenance at the cell centre during metaphase [25, 27, 77, 80]. It positions the flexural rigidity of microtubules as a candidate key factor in spindle choreography. We envisioned that sufficiently high microtubule rigidity is necessary to enable stronger cortical pushing forces. These forces may be required to counterbalance elevated cortical pulling forces during anaphase.

In *C. elegans* zygote, mutations in *zyg-8^dclk1^*, the sole Doublecortin family member, have been shown to cause improper spindle positioning during anaphase, as observed through immunofluorescence and DIC microscopy [54, 81]. ZYG-8 contains a kinase domain at the C-terminus and two doublecortin (DC) domains at the N-terminus, the latter enabling its binding to microtubules. Studies of the human homolog DCLK1 revealed that its kinase domain auto-phosphorylates the DC domains. Besides, hyperphosphorylation of these domains reduces DCLK1’s binding affinity to microtubules [82]. In *C. elegans* zygote, mutations in either domain of *zyg-8* led to spindle orientation defects [54]. It has been proposed that ZYG-8 promotes microtubule assembly, which is required for correct spindle positioning [54, 81]. Interestingly, Srayko *et al.* found that ZYG-8 contributes to microtubule growth, although to a lesser extent compared to the well-established polymerase ZYG-9^XMAP215^ [48]. Besides, ZYG-8 was shown to limit astral microtubule nucleation, although the perturbations were less pronounced than those observed with *spd-2^CEP192^(RNAi)* [48]. Could these moderate effects on microtubule dynamics cause by depleting ZYG-8 be sufficient to explain the spindle mis-positioning seen in *zyg-8* mutants during anaphase? To address this question and better understand ZYG-8’s role in spindle positioning, we used microscopy techniques that allow dynamic studies in live embryos, along with advanced image analysis. We explored the potential roles of ZYG-8 in regulating both microtubule dynamics and flexural rigidity. We applied three complementary genetic perturbations: RNAi-mediated depletion of ZYG-8, overexpression, and the thermosensitive *zyg-8(or484ts)* mutant that disrupts microtubule binding [81]. Upon *zyg-8* targeting, we analysed anaphase spindle oscillations and microtubule cortical contacts. We found that the observed changes could not be explained solely by altered microtubule dynamics. This prompted us to investigate whether ZYG-8 also regulates microtubule rigidity during zygote division. To assess changes in microtubule rigidity, we notably measured the distributions of microtubule local curvatures. We also examined spindle positioning forces using three biophysical assays. Our results demonstrated that ZYG-8 is needed to maintain the correct spindle orientation during anaphase, primarily by permitting high centring forces. Importantly, these centring forces depend on microtubule flexural rigidity, and their elevated levels specifically counterbalance the increased cortical pulling forces observed during anaphase. Overall, our study highlights the need for sufficient microtubule rigidity for proper mitotic spindle positioning and orientation during *C. elegans* zygote division.

## RESULTS

### 1 ZYG-8 affects spindle-pole oscillations beyond microtubule growth and nucleation

In *C. elegans* zygotes, mutations in *zyg-8^dclk1^* disrupted spindle positioning, particularly during anaphase, as shown by DIC microscopy recordings [54, 81]. These mutations were proposed to perturb astral microtubule stability, as microtubules stained with anti-tubulin antibodies appeared shorter. Immunofluorescence revealed that ZYG-8 localises at least to the spindle and its poles [54]. To investigate its association with astral microtubules, we generated three strains using CRISPR-Cas9 genome editing: one expressing fluorescently tagged ZYG-8 (mNeonGreen::ZYG-8) from the endogenous locus; a second with *zyg-8* tagged at the locus with three OLLAS epitope tags; and a third carrying an integrated transgene overexpressing *zyg-8* (*pPie-1*::mNG::3*OLLAS::ZYG-8) in addition to the endogenous copy (Table S1). During the first embryonic division, in both endogenously expressed and overexpressed *zyg-8* strains, ZYG-8 showed significant co-localisation with astral and spindle microtubules (Figure S1, Method M5). This localisation was consistent with its previously proposed role in promoting microtubule assembly [54].

**Figure 1:**
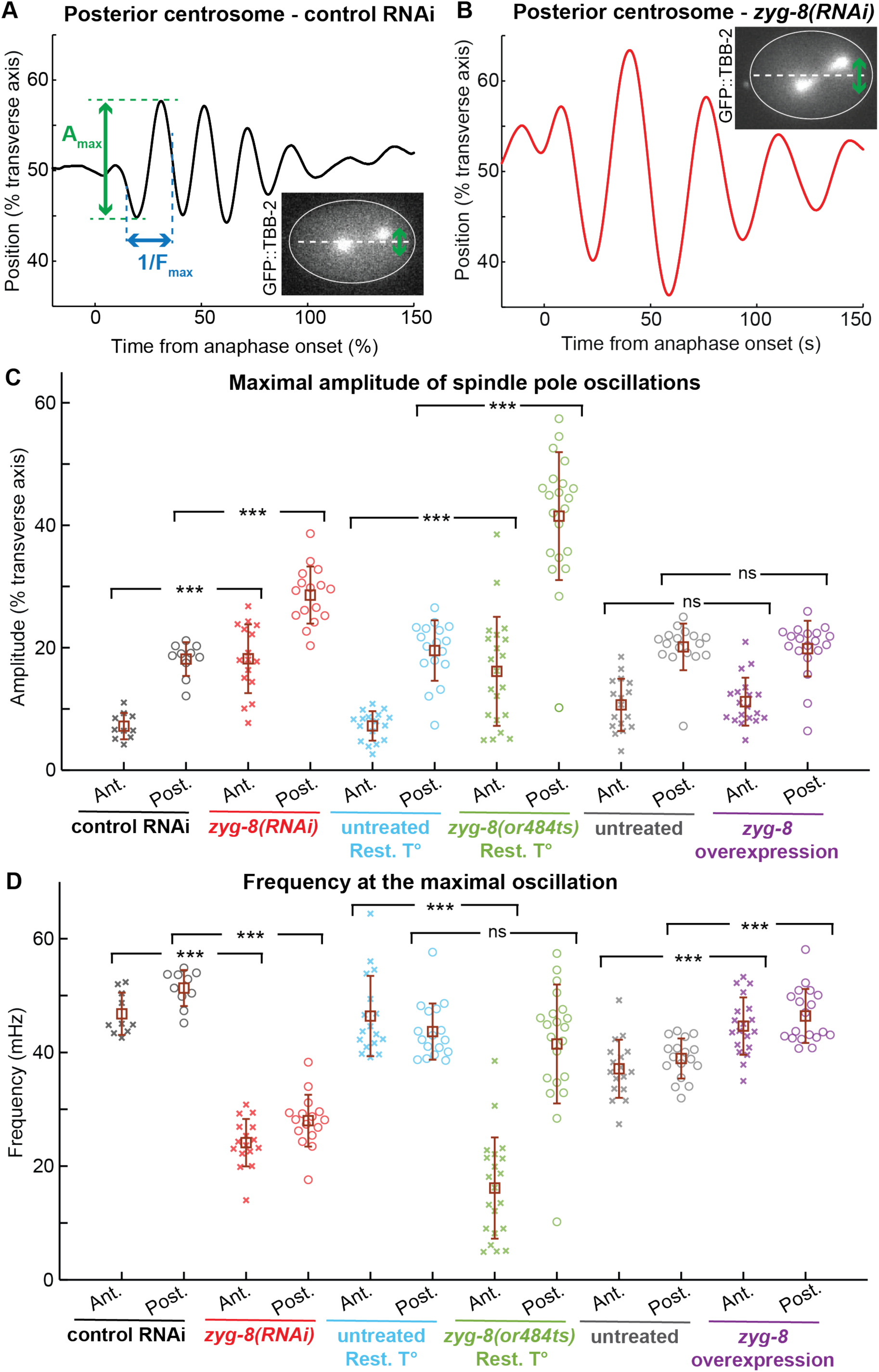
ZYG-8 limits spindle-pole oscillation amplitudes and increases their frequencies during anaphase. (**A-B**) Exemplar posterior centrosome positions along the transverse axis for (A) a control RNAi embryo and (B) a *zyg-8(RNAi)-*treated embryo. Maximal amplitude is highlighted in green, while its frequency is annotated in blue. We applied a moving-average filter over a window size of 5 s to smooth the centrosome positions. Exemplar images used for centrosome analysis with a GFP::TBB-2 fluorescent labelling. (**C**) Maximal oscillation amplitudes and (**D**) their frequencies for the (cross) anterior and (circle) posterior centrosomes during anaphase. We tracked the centrosomes and analysed their positions in: (red) *N* = 16 *zyg-8(RNAi)*-treated embryos and (black) *N* = 10 control RNAi embryos expressing GFP::TBB-2; (light green) *N* = 21 *zyg-8(or484ts)* mutants and (light blue) *N* = 17 untreated embryos, both at the restrictive temperature (Rest. T°) expressing EBP-2::mKate2; (purple) *N* = 19 *zyg-8* overexpressing embryos and (grey) *N* = 17 untreated embryos expressing mCherry::tubulin (Method M7). *N* represents the total number of embryos analysed across all replicates. The brown squares and error bars correspond to the means and SD. *** indicates significant differences (*p* ≤ 1x10^-4^) and ns denotes non-significant differences (*p* > 0.05) (Method M15). Exemplar posterior centrosome positions along the transverse axis, along with representative images for a *zyg-8(or484ts)* mutant and an untreated embryo, both at the restrictive temperature, are shown in Figure S3.

Previous studies in *C. elegans* zygotes focussing on ZYG-8 have either examined its role in spindle positioning using *zyg-8* mutants or investigated its effects on microtubule dynamics through *zyg-8(RNAi)* treatment [48, 54, 81]. We sought to complement these studies by introducing three distinct genetic perturbations targeting *zyg-8*. First, we used the thermosensitive mutant *zyg-8(or484ts)*, which carries a mutation in the first doublecortin domain [54, 83]. This mutant has been reported to prevent ZYG-8’s ability to bind microtubules when maintained at the restrictive temperature of 25°C for more than 12 hours [81]. Second, we used the strain described above that overexpresses ZYG-8. Western blot analysis of worm lysates revealed increased OLLAS levels, indicating elevated ZYG-8 expression, as the OLLAS tag was inserted into the overexpression transgene (Figure S2A, Method M6). Besides, mNG::ZYG-8 fluorescence on both spindle and astral microtubules in zygotes increased approximately threefold (Figure S2C, D & G, Method M6). Third, we reduced ZYG-8 expression using RNAi. Western blot analysis of worm lysates from the strain carrying OLLAS-tagged *zyg-8* showed decreased OLLAS levels upon *zyg-8(RNAi)*, indicating reduced ZYG-8 expression (Figure S2B, Method M6). Besides, we observed an approximately fivefold decrease in mNG::ZYG-8 fluorescence on zygotic microtubules upon *zyg-8(RNAi)* (Figure S2E, F & H, Method M6). These genetic perturbations—loss of ZYG-8 from microtubules, overexpression, and partial depletion—provided the tools to investigate how different ZYG-8 levels affect mitotic spindle choreography.

Since ZYG-8 has been proposed to play a key role in spindle positioning, particularly during anaphase, we focussed on spindle pole oscillations occurring at this stage [66]. These oscillations have been widely used in previous studies to assess perturbations in spindle-positioning forces [67–69, 71, 72]. We quantified both the amplitude and frequency of the maximal oscillation at each spindle pole in strains expressing GFP::TBB-2 (*zyg-8(RNAi)* or *zyg-8* mutants) or mCherry::tubulin (*zyg-8* overexpression) (Method M7). Following *zyg-8(RNAi)*, we observed a large increase in oscillation amplitudes (Figure 1A-C) and a major decrease in frequencies (Figure 1A, B & D). In *zyg-8(or484ts)* mutants at the restrictive temperature, oscillation amplitudes were further elevated (Figures 1C and S3). In mutants, oscillation frequencies at the anterior centrosome were further decreased compared to RNAi treatment, while those at the posterior centrosome were more variable than in other conditions – likely due to spindle misorientation – and therefore not significantly affected (Figure 1D). The stronger phenotype in mutant embryos compared to *zyg-8(RNAi)-*treated embryos was consistent with the partial depletion of ZYG-8 by RNAi (Figure S2B, H). These results suggested that ZYG-8 acts to limit spindle oscillation amplitudes during anaphase. In embryos overexpressing *zyg-8*, maximal amplitudes remained unchanged (Figure 1C), but oscillation frequencies increased significantly (Figure 1D), indicating that oscillation frequency depends on ZYG-8 levels.

**Figure 2:**
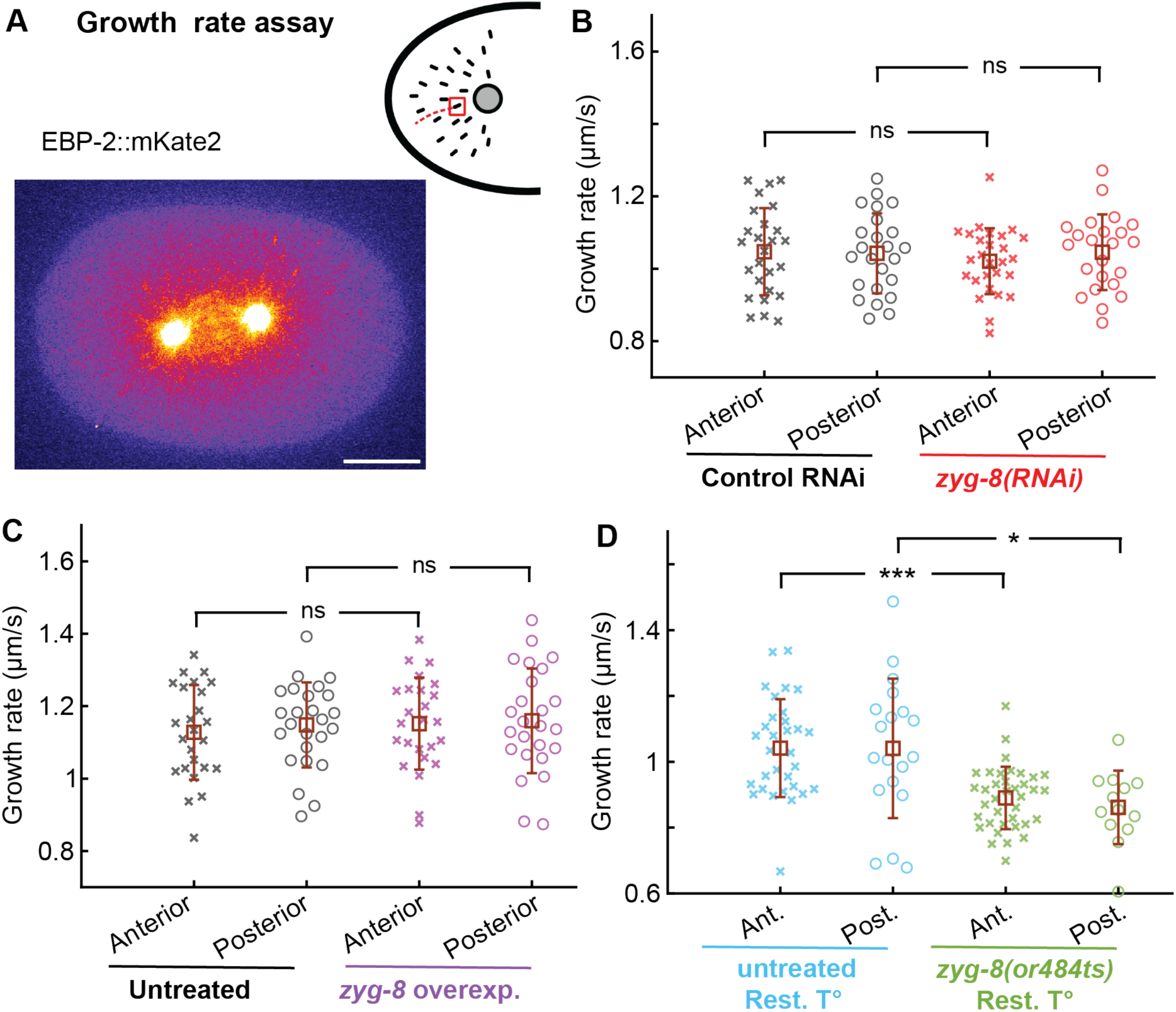
In *zyg-8* mutants, microtubule growth rate is reduced by 15%. (**A**) Microtubule growth rate assay by tracking comets in embryos whose microtubule plus-ends were fluorescently labelled with EBP-2::mKate2 (Method M8). Example of a control RNAi embryo processed with Kalman denoising, a preprocessing step performed prior to quantification, visualised with the Fire LUT. Scale bar represents 10 μm. (**B-D**) Growth rates of astral microtubules measured at (cross) anterior and (circle) posterior centrosomes: (B) (red) *Nc* = 27 comets from anterior centrosome and *Nc* = 23 comets from posterior centrosome in 10 *zyg-8(RNAi)*-treated embryos, and (black) *Nc* = 26 comets from anterior centrosome and *Nc* = 24 comets from posterior centrosome in 8 control RNAi embryos; (C) (purple) *Nc* = 25 comets from anterior centrosome and *Nc* = 25 comets from posterior centrosome in 10 *zyg-8* overexpressing embryos and (black) *Nc* = 25 comets from anterior centrosome and *Nc* = 25 comets from posterior centrosome in 10 untreated embryos; (D) (light green) *Nc* = 37 comets from microtubules emanating from the anterior centrosome and *Nc* = 13 comets from posterior centrosome in 10 *zyg-8(or484ts)* mutants, and (light blue) *Nc* = 31 comets from anterior-centrosome and *Nc* = 19 posterior-centrosome comets in 13 untreated embryos, both at the restrictive temperature (Rest. T°). *Nc* represents the total number of comets analysed across all replicates, separately for anterior and posterior centrosomes, with about 5 comets analysed per embryo. The brown squares and error bars correspond to the means and SD. * and *** indicate significant differences with 1x10^-3^ &< *p* ≤ 1x10^-2^ and *p* ≤ 1x10^-4^, respectively. ns denotes non-significant differences (*p* > 0.05) (Method M15).

We and others have shown that the characteristics of spindle-pole oscillations were sensitive to microtubule dynamics [25, 72]. Besides, a previous study has investigated microtubule growth and nucleation perturbations in *C. elegans* zygotes across a broad range of MAP depletions, including ZYG-8 [48]. This study found that ZYG-8 promotes microtubule growth and limits microtubule nucleation, although its effects were weaker than those of the polymerase ZYG-9^XMAP215^ and the nucleator SPD-2^CEP192^. To further explore the functions of ZYG-8, we quantified both parameters in our three genetic perturbations. We used an EB-labelling of microtubule plus-ends to assess whether the astral microtubule growth rate was altered (Method M8, Figure 2A). We detected no significant changes in growth rate in either *zyg-8(RNAi)*-treated embryos (Figure 2B), for which depletion was partial (Figure S2B), or in those overexpressing *zyg-8* (Figure 2C). This suggested that the oscillation amplitude variations in these two perturbative conditions could have an alternative origin. In *zyg-8(or484ts)* mutants at the restrictive temperature, we observed a significant 15% reduction in microtubule growth rate (Figure 2D), consistent with previous results using *zyg-8(RNAi)* by injection [48]. To test whether this 15% reduction in microtubule growth could account for the exaggerated spindle-pole oscillations observed in *zyg-8* mutants, we performed a partial RNAi depletion of the microtubule polymerase ZYG-9^XMAP215^ [48, 84, 85]. This treatment shortened the mitotic spindle (Figure S4A), confirming efficient depletion [84], and reduced the microtubule growth rate by 16%, comparable to the effect in *zyg-8* mutants (Figure S4B). However, spindle-pole oscillation amplitudes and frequencies were either unaffected or slightly reduced, and did not phenocopy the *zyg-8* mutant phenotype (Figure S4C, D). These results suggested that the limited contribution of ZYG-8 in promoting microtubule growth was not sufficient on its own to account for its critical function in regulating spindle dynamics during anaphase.

Alternatively, ZYG-8 has been proposed to limit microtubule nucleation in the *C. elegans* zygote [48]. To test this, we estimated the nucleation rate of astral microtubules under our three genetic perturbation conditions. Specifically, we counted comet-like structures representing EBP-2::mKate2 labelled plus-ends of growing microtubules emanating from centrosomes (Method M8, Figure 3A). We observed a significant increase in the nucleation rate in *zyg-8(or484ts)* mutants and a decrease upon *zyg-8* overexpression, whereas *zyg-8(RNAi)* had no significant effect (Figure 3B-D). Since centrosome diameter correlates with nucleation capacity [86–88], we also measured centrosome size using mCherry::TBG-1 (Method M9, Figure 3E, F). We found statistically significant differences in centrosome diameter, with a 12% decrease upon *zyg-8* overexpression (Figure 3G) and a 6% increase following *zyg-8(RNAi)* treatment (Figure 3H). These results collectively indicated that ZYG-8 functions to restrict microtubule nucleation. Next, we tested whether altered nucleation alone could explain the spindle-pole oscillation phenotype observed upon *zyg-8* perturbation. To this end, we partially depleted SPD-2^CEP192^ by RNAi to reduce microtubule nucleation and assessed the effects on spindle pole oscillations [89]. *spd-2(RNAi)* caused a 21% reduction in centrosome diameter (Figure S5A, B), an effect comparable to—but slightly more pronounced than—that of *zyg-8* overexpression (Figure 3E, G). While *zyg-8* overexpression increases oscillation frequency by 20% (Figure 1D), *spd-2(RNAi)* decreased frequency by 11% (Figure S5D). Neither condition significantly altered oscillation amplitude (Figures 1C and S5C). Together, these findings suggested that ZYG-8’s role in limiting microtubule nucleation alone was not sufficient to account for the anaphase spindle oscillation phenotypes observed upon *zyg-8* targeting.

**Figure 3:**
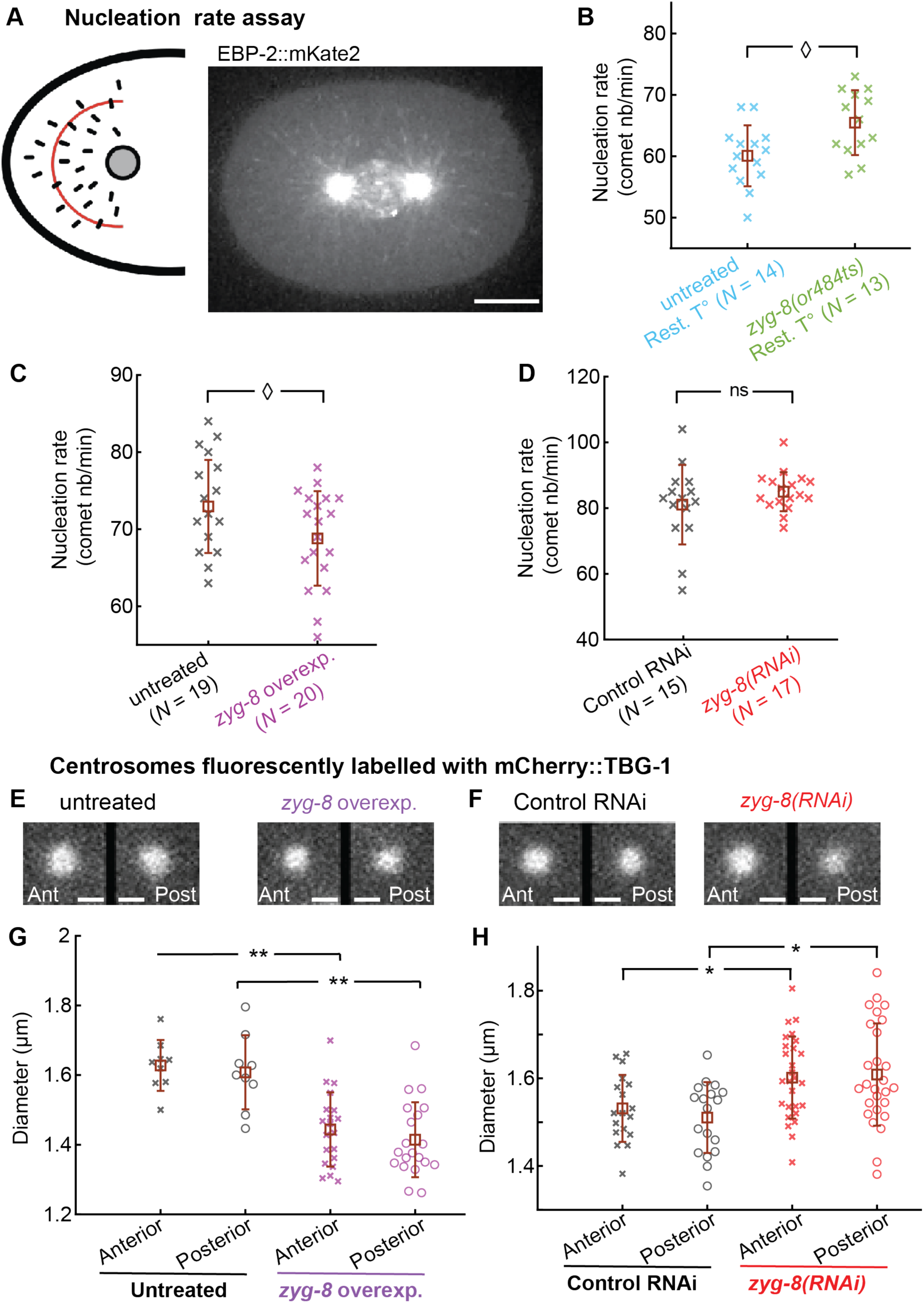
ZYG-8 limits microtubule nucleation. (**A**) Microtubule nucleation rate assay by counting comets crossing a half-circle region of interest positioned 9 µm from the centrosome in embryos whose microtubule plus-ends were fluorescently labelled using EBP-2::mKate2 (Method M8). Example of a control RNAi embryo processed with Candle denoising, a preprocessing step performed prior to comet counting. Scale bars represent 10 μm. **(B-D)** Nucleation rates of the astral microtubules with *N* indicating the number of analysed centrosomes per condition: (A) (light green) *Ne* = 8 *zyg-8(or484ts)* mutants and (light blue) *Ne* = 9 untreated embryos, both at the restricted temperature (Rest. T°); (B) (purple) *Ne* = 10 *zyg-8* overexpressing embryos and (black) *Ne* = 10 untreated embryos; (C) (red) *Ne* = 10 *zyg-8(RNAi)*-treated embryos and (black) *Ne* = 8 control RNAi embryos. (**E-F**) Regions centred on the centrosomes (40 x 40 pixels; 6.4 x 6.4 μm) from exemplar microscopy images showing centrosomes fluorescently labelled with mCherry::TBG-1: (E) representative images of untreated and *zyg-8* overexpressing embryos; (F) representative images of control RNAi and *zyg-8(RNAi)*-treated embryos. Scale bars represent 2 μm. **(G-H**) Diameters during metaphase of (cross) anterior and (circle) posterior centrosomes (Method M9): (G) (purple) *Ne* = 21 *zyg-8* overexpressing embryos and (black) *Ne* = 9 untreated embryos; (H) (red) *Ne* = 27 *zyg-8(RNAi)*-treated embryos and (black) *Ne* = 18 control RNAi embryos. *Ne* represents the total number of embryos analysed across all replicates. The brown squares and error bars correspond to the means and SD. ◊, *, and ** indicate significant differences with 1x10^-2^ &< *p* ≤ 5x10^-2^, 1x10^-3^ &< *p* ≤ 1x10^-2^, and 1x10^-4^ &< *p* ≤ 1x10^-3^, respectively. ns denotes non-significant differences (*p* > 0.05) (Method M15).

### 2 Cortical microtubule dynamics do not support a role for ZYG-8 in microtubule stabilisation

Previous works in human cells and *in vitro* have proposed that members of the Doublecortin family stabilise microtubules by preventing catastrophe and/or inhibiting depolymerisation [49, 90–92]. We investigated whether a similar regulatory mechanism operates in the *C. elegans* zygote. Inspired by a previous study in nematodes on the role of CLS-2^CLASP^ in microtubule stability [93], we imaged, at the cell cortex, embryos expressing GFP::TBB-2 (*zyg-8(RNAi)* treatment) or mCherry::tubulin (*zyg-8* overexpression) and examined both the number and duration of astral microtubule contacts there (Figure 4A, B; Movies S1-S4). Since microtubule catastrophes are rare in *C. elegans* zygote cytoplasm, most astral microtubules grow until they reach the cortex [72, 94]. If ZYG-8 stabilises microtubules by limiting depolymerisation or preventing catastrophe in the cytoplasm, *zyg-8(RNAi)* would be expected to reduce the number of cortical contacts. However, we observed the opposite: *zyg-8(RNAi)* increased the density of cortical contacts by an average of 50%, while *zyg-8* overexpression led to a 13% reduction (Figure 4C, D). These results argued against a cytoplasmic stabilisation role for ZYG-8. Next, to explore whether ZYG-8 might stabilise microtubules at the cell cortex by reducing catastrophe frequency, we analysed the duration of microtubule cortical contacts. If ZYG-8 acted to stabilise microtubules at the cortex, *zyg-8(RNAi)* would be expected to increase the number of short-duration contacts. In contrast, *zyg-8(RNAi)* resulted in a 10-15% reduction in the short-duration contacts (Figure 4E, F). Conversely, *zyg-8* overexpression led to a 7% increase in short-duration contacts (Figure 4G, H). Altogether, these cortical measurements did not support a role for ZYG-8 in stabilising microtubules within the cytoplasm or at the cortex.

**Figure 4:**
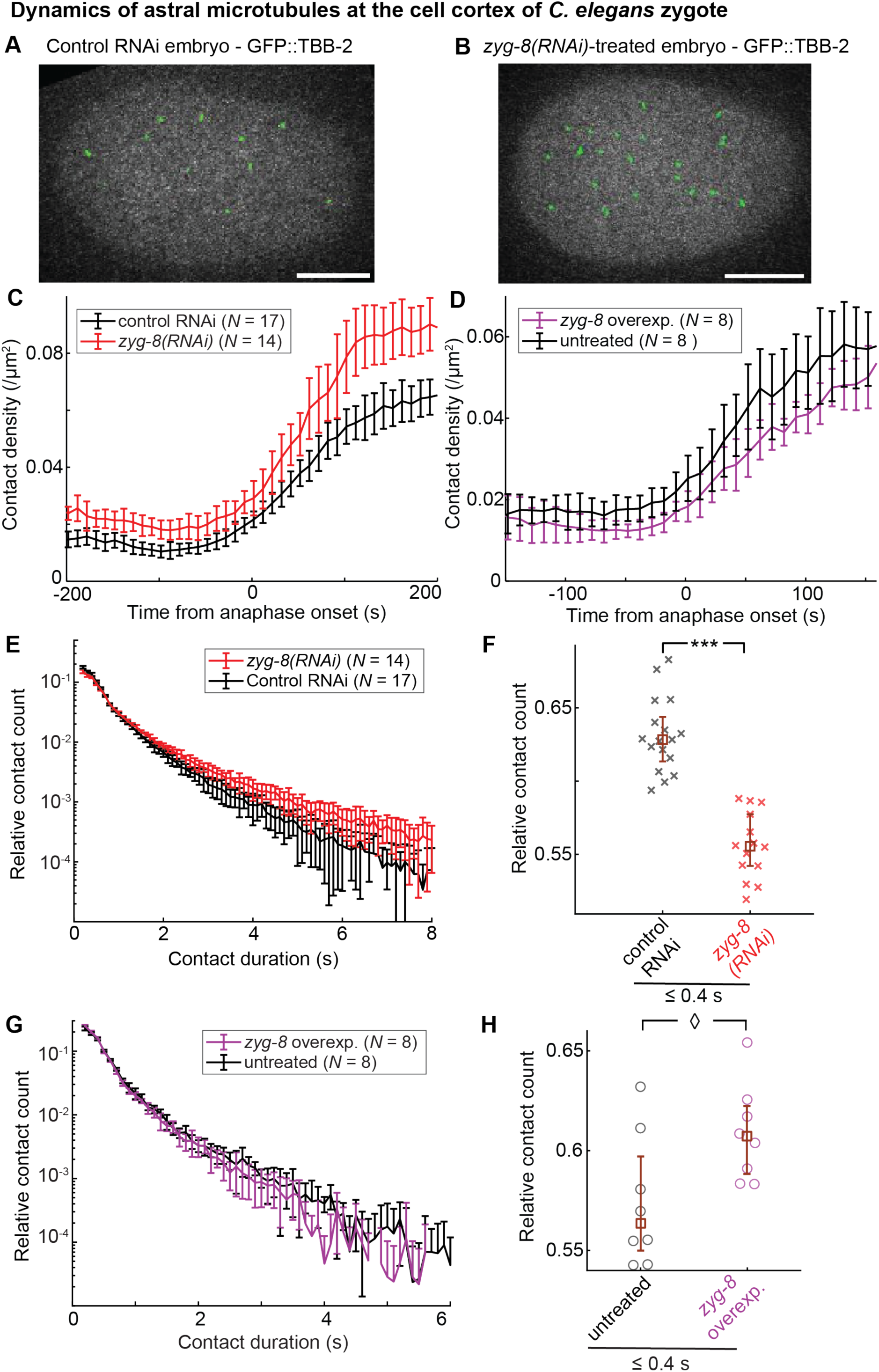
Perturbations in microtubule cortical dynamics upon targeting *zyg-8* do not support a role for ZYG-8 in stabilising microtubules. (**A-B**) Embryos whose microtubules are labelled by GFP::TBB-2 and imaged at the cortical plane (Method M4): examples of (A) a control RNAi embryo (Movie S1), and (B) a *zyg-8(RNAi)*-treated embryo (Movie S2) at anaphase onset. Green spots highlight the detected contacts (Method M10). Scale bars represent 10 μm. (**C-D**) Embryo-averaged density of astral microtubule contacts at the cortex plane along mitosis. We tracked and analysed the cortical contacts of microtubules labelled: (C) by GFP::TBB-2 in (red) *N* =14 *zyg-8(RNAi)*-treated embryos and (black) *N* = 17 control RNAi ones; (D) by mCherry::tubulin in (purple) *N* = 8 *zyg-8* overexpressing embryos and (black) *N* = 8 untreated ones. Embryo-averaged distributions of cortical lifetimes of astral microtubules during metaphase (Kolmogorov–Smirnov test: *p* = 6x10^-4^, Methods M10 and M15). Embryos are the same as in panel C. Relative counts of contacts with durations ≤ 0.4 s (from distributions presented in panel E), highlighting differences in brief contact durations. (**G**) Embryo-averaged distributions of cortical lifetimes of astral microtubules during metaphase (Kolmogorov–Smirnov test: *p* = 0.18, Methods M10 and M15). Embryos are the same as in panel D. (**H**) Relative counts of contacts with durations ≤ 0.4 s (from distributions in panel G), highlighting variations in short-lived durations. Representative movies of each of the four present conditions are provided as Movies S1-S4. Microtubule contacts were tracked using u-track (Method M10). *N* represents the total number of embryos analysed across all replicates. The brown squares and error bars correspond to the medians and the quartiles. ◊ and *** indicate significant differences with 1x10^-2^ &< *p* ≤ 5x10^-2^ and *p* ≤ 1x10^-4^, respectively (Method M15). This figure does not include data for *zyg-8(or484ts)* mutant, as severe spindle misorientation precluded reliable quantification for this analysis.

To further test this finding, we compared cortical microtubule dynamics and spindle pole oscillations in *zyg-8(RNAi)*-treated embryos with those following depletion of CLS-2^CLASP^—a known microtubule stabiliser that prevents catastrophe and promotes rescue [93, 95]. We validated the penetrance of the RNAi treatment by observing accelerated mitotic spindle elongation, which suggested midzone integrity perturbations [97, 98] (Figure S6A). At the cell cortex, in *cls-2(RNAi)*-treated embryos, we found a reduced density of microtubule contacts during anaphase (Figure S6F), and a decreased proportion of long-duration contacts (Figure S6G). These results aligned with CLS-2’s known role in catastrophe prevention, but differed from *zyg-8(RNAi)* phenotypes, further suggesting that ZYG-8 does not primarily prevent microtubule catastrophe in *C. elegans* zygote. Notably, *cls-2(RNAi)* caused an increase in oscillation amplitude, although weakly significant, with no significant variation in oscillation frequency (Figure S6B, C). These phenotypes were distinct from *zyg-8(RNAi)*. This suggested that altered microtubule catastrophe alone induces only minor perturbations in spindle pole oscillations and, therefore, could not explain the pronounced oscillation phenotypes observed in *zyg-8(RNAi)* embryos.

We then compared *zyg-8(RNAi)* phenotypes with those following hypomorphic depletion of KLP-7^MCAK^, which destabilises microtubules by promoting depolymerisation [96]. We confirmed *klp-7(RNAi)* treatment penetrance by observing slightly increased spindle elongation (suggesting only a mild alteration of midzone integrity) and longer metaphase spindle, as expected [99] (Figure S7A). At the cell cortex, KLP-7 depletion increased the microtubule contact density and the proportion of long-duration contacts (Figure S7F, G). These findings reflected impaired depolymerisation activity. If ZYG-8 stabilised microtubules by preventing depolymerisation, we would have observed cortical microtubule phenotypes opposite to those of KLP-7 depletion. This was not the case. Regarding spindle pole oscillations, KLP-7 depletion increased amplitudes at the posterior centrosome and decreased frequencies at both centrosomes (Figure S7B, C)—phenotypes not opposite to *zyg-8(RNAi)*. Thus, a putative role of ZYG-8 in preventing microtubule depolymerisation could not account for the observed phenotypes.

Overall, our data indicated that the *zyg-8*-associated perturbations in spindle-pole oscillations and cortical microtubule dynamics could not be fully attributed to alterations in microtubule dynamics alone. Interestingly, proteins of the Doublecortin family have been proposed to regulate microtubule rigidity in neurons [37, 100]. We asked whether the observed perturbations in spindle-pole oscillations and cortical microtubule behaviour could additionally result from altered microtubule rigidity upon *zyg-8* targeting.

### 3 ZYG-8 would control microtubule flexural rigidity

Direct measurements of microtubule rigidity in *C. elegans* embryos are not feasible, as standard techniques (e.g., thermal fluctuations, optical trap/tweezers, or calibrated flow) require isolated microtubules [15–17, 34, 101–103]. As a quantitative alternative, we employed a set of complementary metrics that collectively capture different aspects of microtubule bending behaviour (Method M11). Such an approach has been previously used as a proxy for rigidity under the assumption that the pulling forces acting on the microtubule network are similar between conditions [20, 104, 105]. Notably, microtubules in axonal growth cones were more often strongly curved upon depletion of the Doublecortin-family proteins DCX and DCLK1 [37]. First, we analysed local curvature distributions: a higher proportion of strongly curved segments reflects reduced rigidity (Figure S8A). To complement this population-level analysis with a more filament-specific readout, we extracted the 95^th^ percentile of curvature values along individual microtubules, as a robust measure of maximum bending per microtubule (Figure S8B). Finally, we incorporated filament tortuosity—a global shape descriptor independent of curvature computation and less sensitive to sampling resolution. Defined as the ratio between curvilinear and end-to-end distances, tortuosity has also been used previously [35] (Figure S8C).

We first asked whether astral microtubules displayed increased local curvature in *zyg-8(or484ts)* mutants or in *zyg-8(RNAi)*-treated embryos. We imaged live embryos expressing GFP::TBB-2 using a super-resolved confocal microscope (Nikon with Nsparc detector, Method M4). We observed regions of high local curvatures along astral microtubules, which appeared more frequently in the mutants (Figure 5A, Movies S5 and S6) than in untreated embryos (Figure 5B and Movie S7), both at the restrictive temperature. Besides, astral microtubules in the mutants were long and extended to the cell periphery. Consistently, astral microtubules exhibited increased bending in live *zyg-8(RNAi)*-treated embryos (Figure S9A, Movie S8) compared to control RNAi embryos (Figure S9B, Movie S9). In contrast, *zyg-8* overexpression in live embryos did not significantly alter microtubule curvature amplitudes or frequencies (Figure S10, Movies S10 & S11). The fast motion of astral microtubules required high image acquisition rate to avoid motion blur, which led, unfortunately, to images with low signal-to-noise ratio complicating microtubule curvature quantification. In *zyg-8(or484ts)* mutants, significant out-of-plane movements hindered microtubule visualisation along their full length, calling for acquisitions on multiple z-sections (Movies S5 & S6). As a result, we opted for investigating curvature phenotypes in fixed samples.

**Figure 5:**
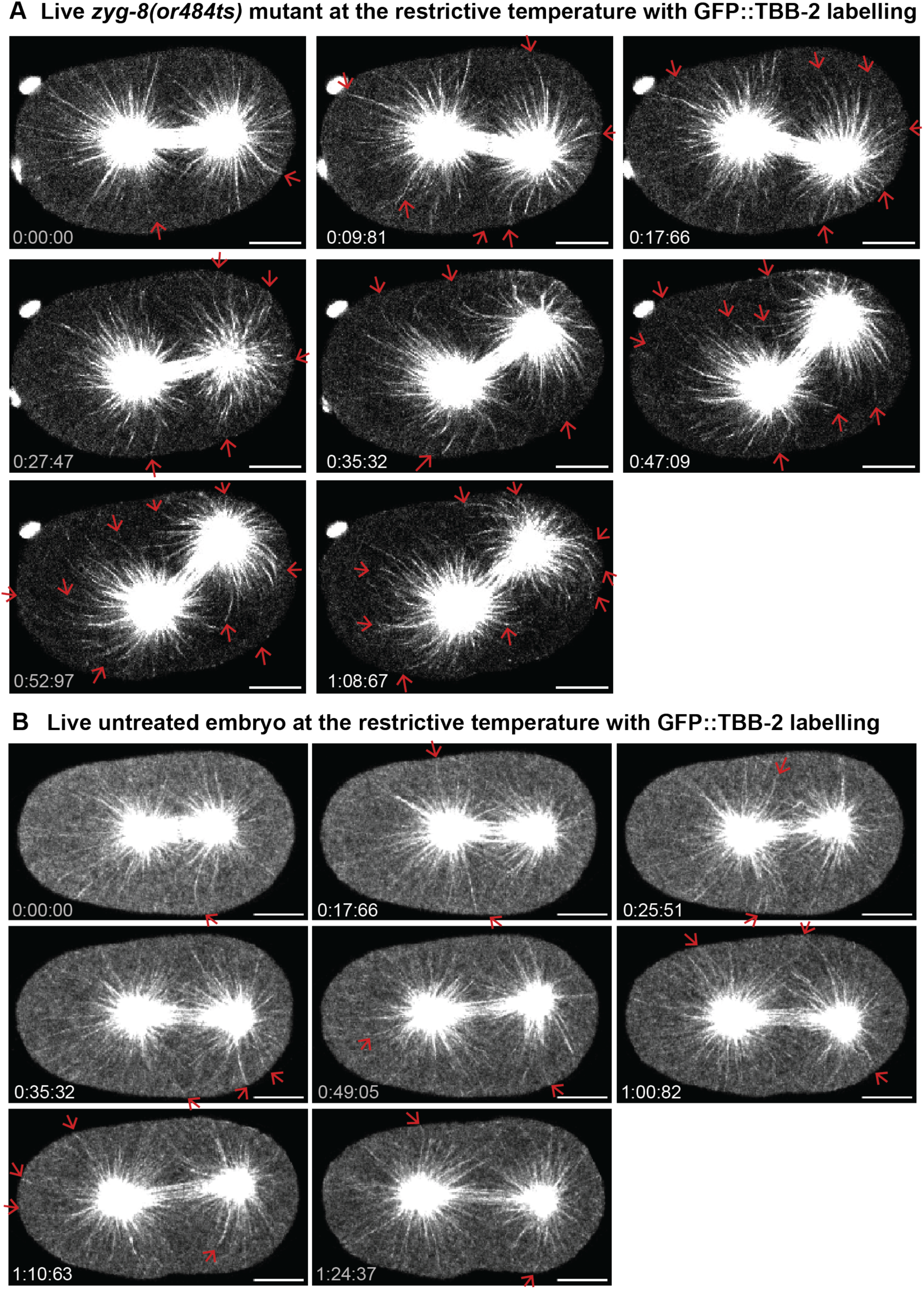
In *zyg-8(or484ts)* mutants, astral microtubules are highly bent during the exaggerated spindle pole oscillations. Time-lapse confocal super-resolved images along anaphase of (**A**) an exemplar *zyg-8(or484ts)* mutant and (**B**) an exemplar untreated embryo, both at the restrictive temperature. Microtubules were labelled by GFP::TBB-2. Time is indicated from first image in mm:ss:ms. Scale bars represent 10 μm. These time-lapse images are sourced from the movies S5 and S7. Red arrows highlight bent microtubules. Similar images for *zyg-8(RNAi)* and *zyg-8* overexpression are shown in Figures S9 and S10, with associated movies provided as Movies S8-S11.

We imaged microtubules stained for α-tubulin using a confocal super-resolution microscopy (Methods M3 and M4). In *zyg-8(RNAi)*-treated fixed embryos, we again observed more bent astral microtubules (Figure S11A, B). This phenotype was even more pronounced in *zyg-8(or484ts)* fixed mutants at the restrictive temperature, where microtubule curvatures were noticeably increased (Figure S11C, D). We also detected fragmented microtubules in the cytoplasm of some fixed mutant embryos (Figure S11D). Interestingly, such fragmentation may result from excessive curvatures, as previous studies have linked microtubule breakage to strong bending [104, 106, 107]. Fragmentation could also contribute to the appearance of shorter astral microtubules. This phenotype—very-short astral-microtubule—was observed only in a subset of *zyg-8(or484ts)* fixed mutants, resembling the initial images of stained microtubules in this mutant [54, 81].

To go beyond visual inspection, we quantified the distribution of microtubule curvatures. Accurate segmentation was essential for this analysis but proved challenging due to faint and uneven microtubule labelling. To overcome this, we assembled a five-step image processing pipeline enabling the segmentation of astral microtubules and computation of local curvatures on a pixel-by-pixel basis using a three-point method (Figure S8D-J, Method M11). By averaging curvature distributions across conditions, we observed a slight decrease in the proportion of low local curvatures following *zyg-8(RNAi)* treatment (Figure 6A, B). In *zyg-8(or484ts)* mutants at the restrictive temperature, the distribution shifted significantly further toward higher curvature values compared to controls, showing a stronger effect than RNAi (Figure 6C, D). Notably, some local curvature values reached up to 0.8 μm^-1^, consistent with curvature thresholds associated with microtubule breakage [106, 107]. Focussing on the highest curvatures, we observed increases of 13% in metaphase and 12% in anaphase following *zyg-8(RNAi)*. These increases were further amplified in *zyg-8(or484ts)* mutants, reaching 33% and 42% in metaphase and anaphase, respectively (Figure 6E, F). Lastly, microtubule tortuosity was significantly elevated following *zyg-8(RNAi)* treatment, and increased even further in *zyg-8(or484ts)* mutants at the restrictive temperature (Figure 6G, H). Together, these analyses demonstrated that ZYG-8 modulates microtubule curvature and tortuosity, supporting its role in regulating microtubule flexural rigidity.

We foresaw that reduced rigidity would lead to increased bending, resulting in weaker pushing forces against the cortex. This, in turn, would delay force-dependent catastrophes and increase the cortical lifetimes of astral microtubules [108, 109] (Figure 7A). We used the established agent-centred simulation *Cytosim* [110] to test this hypothesis (Table S2, Method M12). We conducted three sets of simulations in which only the rigidity of astral microtubules was varied, using values of 2, 10, and 25 pN.μm^2^ (Movies S12-S14, Figure S12A). These values were based on *in vitro* measurements of microtubule rigidity [15–17, 101, 103]. We found that these *in silico* changes in rigidity affected both microtubule curvature and tortuosity. Specifically, the highest curvatures and tortuosity increased as rigidity decreased (Figure S12B, C). Importantly, microtubules with lower rigidity exhibited longer cortical lifetimes, supporting our initial prediction (Figure S12D). Returning to *in vivo* experiments, we measured the cortical contact durations of astral microtubules in *zyg-8(RNAi)*-treated embryos expressing GFP::TBB-2 and found an increased proportion of microtubules with prolonged contacts compared to control RNAi embryos (Figure 4E). This observation was consistent with reduced microtubule rigidity in *zyg-8(RNAi)*-treated embryos.

**Figure 6:**
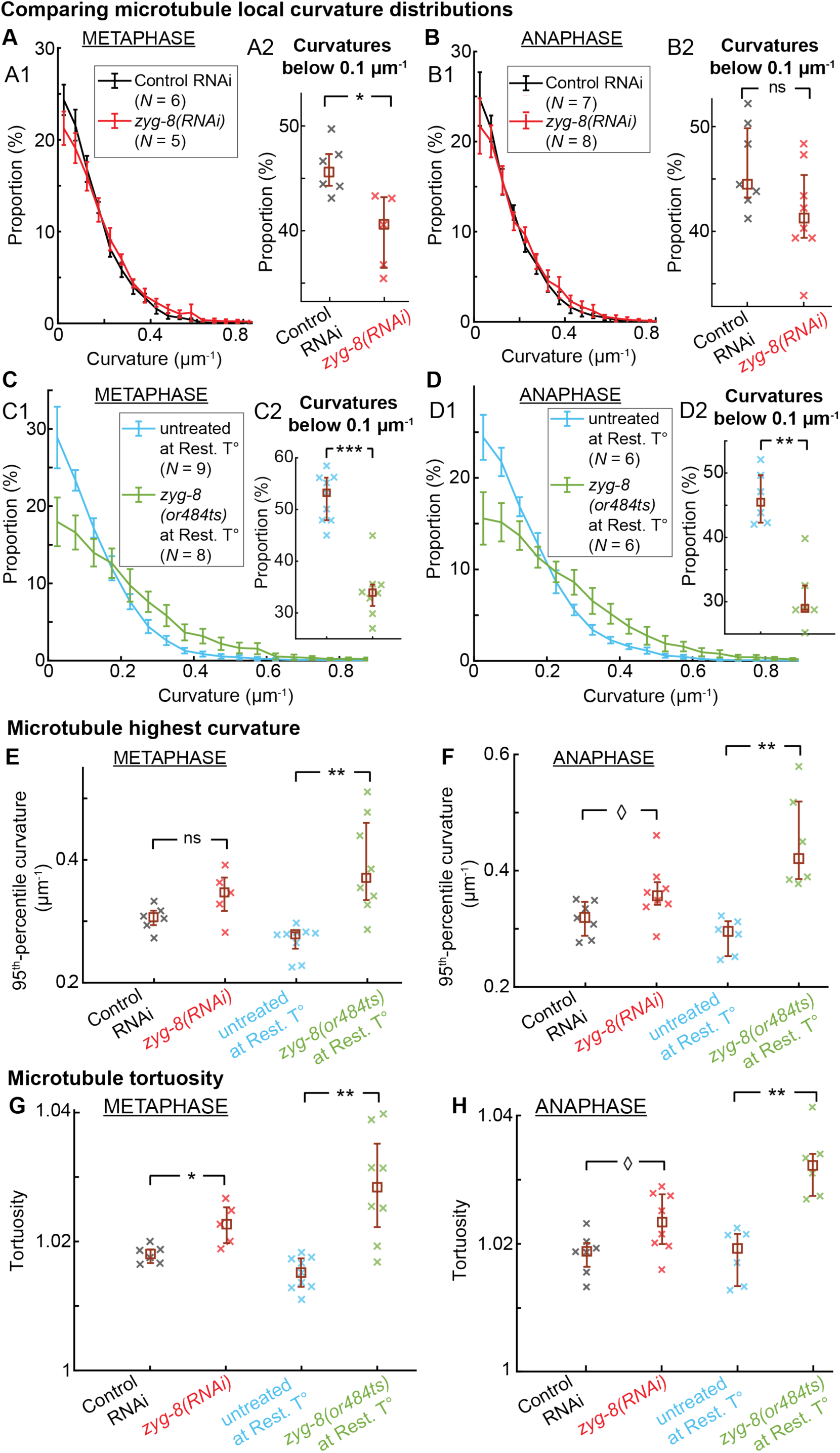
Astral microtubules appear more curved and tortuous in *zyg-8(RNAi)*-treated embryos, and even more so in *zyg-8(or484ts)* mutants. Characterisation of microtubule shapes in fixed embryos analysed using the image-processing pipeline depicted in Figure S8. (**A-B**) Embryo-averaged distributions of astral microtubule local curvatures of (red) *zyg-8(RNAi)*-treated embryos and (black) control RNAi embryos: (A1) during metaphase with (red) *N* = 5 z-projected stacks and (black) *N* = 6 z-projected stacks (Kolmogorov–Smirnov test: *p* = 0.36, Method M15); and (B1) during anaphase with (red) *N* = 8 z-projected stacks and (black) *N* = 7 z-projected stacks (Kolmogorov–Smirnov test: *p* = 0.36, Method M15). (A2, B2) Focus on the proportions of low local curvatures (≤ 0.01 μm^-1^) for the same embryos as in panels A1 & B1. **(C-D)** Embryo-averaged distributions of astral microtubule local curvatures of (light green) *zyg-8(or484ts)* mutants and (light blue) untreated embryos, both at the restrictive temperature: (C1) during metaphase with (light green) *N* = 8 z-projected stacks and (light blue) *N* = 9 z-projected stacks (Kolmogorov–Smirnov test: *p* = 0.03, Method M15); and (D1) during anaphase with (light green) *N* = 6 z-projected stacks and (light blue) *N* = 6 z-projected stacks (Kolmogorov–Smirnov test: *p* = 0.014, Method M15). (C2, D2) Focus on the proportions of low local curvatures (≤ 0.01 μm^-1^) for the same embryos as in panels C1 & D1. (**E-F**) Medians per z-projected stack of the highest curvatures (measured as 95^th^ percentile curvature): (E) during metaphase (data are the same as in panels A & C); and (F) during anaphase (same data as in panels B & D). (**G-H**) Medians per z-projected stack of microtubule tortuosity: (G) during metaphase, using the same embryos as in panels A, C & E; and (H) during anaphase, using the same embryos as in panels B, D & F. Embryos were fixed and immuno-stained for α-tubulin; microtubules were segmented; and local curvature and tortuosity were computed (Methods M3, M4, M11). *N* represents the total number of z-projected stacks analysed across all replicates. The brown squares and error bars correspond to the medians and the quartiles. ◊, *, **, and *** indicate significant differences with 1x10^-2^ &< *p* ≤ 5x10^-2^, 1x10^-3^ &< *p* ≤ 1x10^-2^, 1x10^-4^ &< *p* ≤ 1x10^-3^, and *p* ≤ 1x10^-4^, respectively. ns denotes non-significant differences (*p* > 0.05) (Method M15). This figure does not include data for *zyg-8* overexpressing embryos, as live imaging (Figure S10) demonstrated no significant differences in the astral microtubule curvatures.

Finally, we investigated spindle pole oscillations in light of ZYG-8’s role in microtubule rigidity. We previously proposed an antagonistic-motors model to explain these oscillations, in which the oscillation frequency is in particular proportional to the square root of the centring rigidity [66]. This centring rigidity has been associated with the pushing force generated by microtubules growing against the cortex [25, 77]. Consequently, lower microtubule rigidity—leading to reduced pushing force—was expected to lower centring rigidity, and thus decrease oscillation frequency. Indeed, we observed a reduction in oscillation frequency in *zyg-8(RNAi)*-treated embryos (Figure 1D). Conversely, this frequency was increased in embryos overexpressing *zyg-8* (Figure 1D).

Taken together, these three experiments supported a model in which ZYG-8 contributes to microtubule stiffening.

### 4 ZYG-8 ensures spindle centring mostly by controlling microtubule rigidity

Although the centring force generated by microtubules pushing against the cell cortex—critically dependent on microtubule rigidity—was mostly invoked in metaphase [25, 27, 80], it is also thought to act as a restoring force for spindle rocking in anaphase [66, 69]. Indeed, we previously proposed that, when a spindle pole moves toward one side of the cortex, the centring spring opposes this displacement [66]. Consequently, a reduction in microtubule rigidity would decrease the stiffness of this centring spring, allowing greater spindle pole displacement and thereby increasing oscillation amplitudes (Figure 8G). In *zyg-8(RNAi)* condition or in *zyg-8(or484ts)* mutants, we observed a large increase in the amplitudes of spindle pole oscillations (Figure 1C). We therefore considered two possible and non-mutually exclusive explanations for this increase: enhanced cortical pulling or reduced cortical pushing.

To distinguish between these possibilities, we first measured the most posterior position of the posterior centrosome, projected along the anteroposterior (AP) axis, during anaphase following *zyg-8(RNAi)* treatment. We found no significant difference compared to controls (80.7 ± 1.4 % of AP axis for *zyg-8(RNAi)* (*N* = 16), versus 79.8 ± 1.0 % for control RNAi embryos (*N* = 10), Student’s *t*-test: *p* = 0.07)). This suggested that posterior displacement, driven by cortical pulling forces, remained largely unaltered. Next, we applied the DiLiPop analysis, a biophysical method that enables to assess cortical pulling and pushing events during anaphase. Our previous work has shown that two distinct populations of astral microtubules reside at the cell cortex, disentangled by statistical analysis of cortical contact durations [80] (Method M10). By genetically perturbing either pulling or pushing cortical forces, we previously associated the short-lived population with pulling events and the long-lived population with pushing [80]. Upon *zyg-8(RNAi)*, we observed a 10% rise in the lifetime of the pulling events (Figure 7B) and a 3% reduction in their density (Figure 7C). Since these measurements reflect the number of events (i.e., the number of microtubule cortical tracks, not the instantaneous density of cortical contacts), longer-lasting pulling events (increased lifetimes) reduce the number of distinct pulling events that can occur within the same time window (reduced density). This occurs because all available cortical force generators are engaged in pulling during anaphase [66, 72, 75], explaining the simultaneous observation of increased pulling-event lifetimes and decreased pulling-event density in *zyg-8(RNAi)* embryos. However, these modest changes in pulling events could not account for the strong increase in oscillation amplitudes. For comparison, when we depleted GBP-1 to enhance cortical pulling force [111–113], we observed a significant increase in pulling-event density, as expected, while the lifetimes of both pulling and pushing events remained unaffected (Figure 7F-I). This phenotype differed from that of *zyg-8(RNAi)*, leading us to conclude that *zyg-8(RNAi)* had at most a minor effect on cortical pulling forces. Concurrently, we found that pushing microtubules resided at the cortex significantly longer upon *zyg-8(RNAi)*, with a 37% increase in lifetimes (Figure 7D). This was consistent with their sensitivity to microtubule rigidity (Figure 7A). Besides, the density of pushing events increased by 22% (Figure 7E), partially due to an enhanced microtubule nucleation capacity (Figure 3F). Despite the rise in pushing event density, we proposed a global reduction in effective cortical pushing force, since a bent microtubule has a limited pushing force (Euler force). Consequently, the larger oscillation amplitudes observed following *zyg-8(RNAi)* treatment likely resulted from reduced pushing force, while cortical pulling forces appeared to be largely unaffected.

To further investigate whether cortical pulling forces remain unaffected when targeting *zyg-8*, we tracked cortical dynein using a CRISPR strain expressing GFP::DHC-1 [114] (Method M13), a biophysical assay previously employed [114, 115]. It is important to note that only cortical dynein molecules incorporated into at least the GPR-1/2-LIN-5-dynein trimeric complex actively generate pulling forces on astral microtubules [74, 116]. In control RNAi embryos, we identified two distinct dynein populations at the cell cortex: a short-lived population with a lifetime of 0.66 s, and a long-lived population with a lifetime of 1.66 s (Table S3). The lifetime of the predominant short-lived population aligns with previously reported measurements [115], and falls within the range of cortical pulling event lifetimes (Figure 7B, F), suggesting that this population likely includes active pulling events. Upon *zyg-8(RNAi)*, our analysis revealed similar dynein dynamics (Table S3). The increased dynein density aligns with the elevated microtubule contact density (Figure 4C), while the prolonged dynein lifetimes mirror the increased microtubule cortical contact durations. Crucially, the magnitude of these changes in dynein dynamics was comparable to those observed for microtubule characteristics, suggesting no additional regulatory mechanism. Given that not all cortical dynein is engaged in pulling [114, 115] and that cortical force generators are saturated (i.e., actively pulling) during anaphase in untreated embryos [66, 72, 75], it is unlikely that ZYG-8 depletion substantially increases cortical pulling forces.

**Figure 7:**
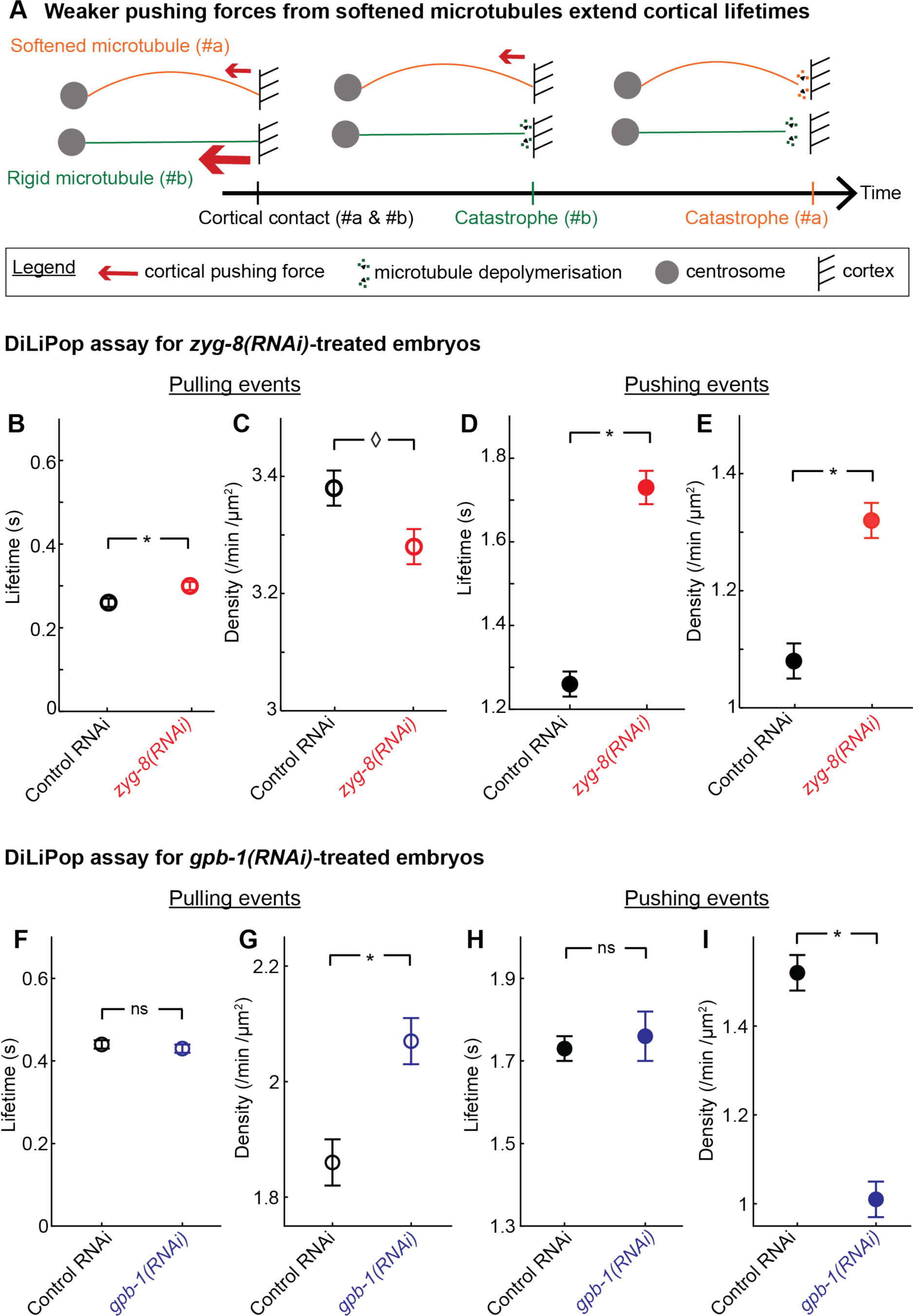
Depletion of ZYG-8 significantly weakens the cortical pushing events. **(A)** Diagram illustrating the correlation between microtubule rigidity and cortical lifetime: (orange) softened microtubules lead to increased pushing-event lifetimes compared to (green) rigid microtubules, due to weaker pushing force (red arrow) delaying catastrophe. Note that rescue of depolymerising astral microtubules is rare in *C. elegans* zygote cytoplasm, supporting continued depolymerisation. (**B, F**) Lifetimes and (**C, G**) densities of the cortical pulling events, and (**D, H**) lifetimes and (**E, I**) densities of the cortical pushing events during anaphase. We tracked and analysed the microtubule contacts at the cell cortex (B-E) using a GFP::TBB-2 fluorescent labelling in (red) *N* = 14 *zyg-8(RNAi)*-treated embryos and (black) *N* = 17 control RNAi embryos (same embryos as for Figure 4C, E & F), or (F-I) using a YFP::TBA-2 fluorescent labelling in (blue) *N* = 8 *gpb-1(RNAi)*-treated embryos and (black) *N* = 9 control RNAi embryos (Method M10). *N* represents the total number of embryos analysed across all replicates. ◊ and * indicate significant differences with 1x10^-2^ &< *p* ≤ 5x10^-2^ and 1x10^-3^ &< *p* ≤ 1x10^-2^, respectively. ns denotes non-significant differences (*p* > 0.05) (Method M15). This figure does not include data for *zyg-8(or484ts)* mutant, as severe spindle misorientation precluded reliable quantification. Additionally, data for *zyg-8* overexpressing embryos are not included, as this figure focuses on elucidating the origin of increased spindle-pole oscillation amplitudes—a phenotype not observed in this condition (Figure 1C).

To examine the role of ZYG-8 in controlling microtubule rigidity—and, in turn, the centring force—we sought an alternative characterisation of spindle positioning forces at the cellular level. To do this, we used a biophysical assay previously developed by our laboratory focusing on the spindle positional micro-movements acquired at high frame rate. We extracted the mechanical fingerprint of these micro-movements along the transverse axis during anaphase. By fitting a second-order model to the experimental spectrum, we derived three mechanical parameters that characterise spindle position mechanics: the diffusion coefficient *D* and two corner frequencies *f_c_* and *f_0_* [25] (Method M14). Our analysis revealed that ZYG-8 depletion caused a large 79% reduction in the centring-to-damping corner frequency *f_c_*, compared to the control (Table 1). We also observed a 24% decrease in the diffusion coefficient *D*, which scales inversely with damping, and a 35% increase in the damping-to-inertia frequency *f_0_* (Table 1). We interpreted the decrease in *f_c_* as primarily reflecting a centring force reduction due to softer microtubules [25]. The moderate decrease in *D* likely resulted from an increased number of microtubules contacting the cortex. The rise in *f_0_* could be partly attributed to the reduced *D*, and appeared inconsistent with any notable increase in inertia, suggesting that the pulling forces were only mildly or not significantly affected. Overall, the mechanical fingerprint of spindle positioning following *zyg-8(RNAi)* treatment was qualitatively consistent with a reduction in the centring force, mainly due to softer microtubules. This reduction likely contributed to the increased amplitude of spindle pole oscillations.

**Table 1:** Targeting *zyg-8* or *ptl-1* reduces the centring frequency. Mechanical parameters measured during anaphase, including the diffusion coefficient (*D*) and two corner frequencies (*fc* and *f0*). They were obtained by fitting the experimental power spectra of spindle positional micromovements along the transverse axis within the specified frequency interval using a second-order model (Method M14). The confidence intervals were estimated using the bootstrap resampling method with a significance level of 0.05. *N* represents the total number of embryos analysed across all replicates for each experimental condition.

| Conditions | Frequency interval | $D$ (m<sup>2</sup>/s) | $f_c$ (Hz) | $f_o$ (Hz) |
| --- | --- | --- | --- | --- |
| Control RNAi ($N = 15$) | 0.03 -> 0.3 Hz | $1.7 \times 10^{-13}$<br>[0.7 - $15.0 \times 10^{-13}$] | 0.86<br>[0.03 - 6.68] | $3.1 \times 10^{-3}$<br>[0.4 - $51.4 \times 10^{-3}$] |
| *zyg-8(RNAi)* ($N = 20$) | | $1.3 \times 10^{-13}$<br>[0.7 - $166 \times 10^{-13}$] | 0.18<br>[0.13 - 5.78] | $4.2 \times 10^{-3}$<br>[0.1 - $7.4 \times 10^{-3}$] |
| *ptl-1(RNAi)* ($N = 13$) | | $1.0 \times 10^{-13}$<br>[0.8 - $4.9 \times 10^{-13}$] | 0.34<br>[0.09 - 5.46] | $5.9 \times 10^{-3}$<br>[0.1 - $83.3 \times 10^{-3}$] |
| Untreated at the restrictive temperature ($N = 11$) | 0 -> 0.5 Hz | $7.3 \times 10^{-14}$<br>[5.8 - $9.0 \times 10^{-14}$] | 0.19<br>[0.15 - 0.22] | $1.3 \times 10^{-2}$<br>[1.1 - $1.5 \times 10^{-2}$] |
| *zyg-8(or484ts)* at the restrictive temperature ($N = 12$) | | 11.3<br>$\times 10^{-14}$<br>[6.2 - $687.9 \times 10^{-14}$] | $0.9 \times 10^{-3}$<br>[0.89 - $0.95 \times 10^{-3}$] | $1.3 \times 10^{-2}$<br>[0.01 - $2.4 \times 10^{-2}$] |

In *zyg-8(or484ts)* mutants at the restrictive temperature, we observed larger amplitudes of spindle-pole oscillations compared to those seen with *zyg-8(RNAi)* treatment (Figures 1C, S3). We hypothesised that this phenotype was due to a severely weakened centring force caused by excessively soft microtubules. This was supported by the increased proportion and magnitude of microtubule curvatures observed in these mutants (Figures 5 and 6). To test this hypothesis, we analysed spindle positional micro-movements in *zyg-8(or484ts)* mutants. We found a very large and significant reduction in the centring-to-damping corner frequency *f_c_*, while the other two mechanical parameters, *D* and *f_0_*, were not significantly altered (Table 1). These results suggested that in *zyg-8(or484ts)* mutants, the pronounced reduction in *f_c_*, indicative of weakened centring forces, accounted for the exaggerated spindle pole oscillations. This reduction was likely due to decreased microtubule rigidity, consistent with the observed abnormalities in microtubule shapes.

Altogether, our findings demonstrated that ZYG-8 regulates the cortical pushing force responsible for spindle centring, primarily by modulating microtubule flexural rigidity. This pushing force acted as a restoring force during anaphase spindle rocking. While cortical pulling forces remain the dominant drivers of spindle posterior displacement, our work highlighted the important, though often underappreciated, contribution of pushing forces to overall spindle choreography.

### 5 Very soft microtubules disrupt final spindle positioning and orientation

Having identified an additional role for ZYG-8 in controlling microtubule flexural rigidity, we next investigated the importance of this function for successful asymmetric mitosis. In *zyg-8(or484ts)* mutants at the restrictive temperature, the enhanced spindle-pole oscillation amplitudes – greater than those observed in *zyg-8(RNAi)* – resulted in a significantly reduced centrosome-to-cortex distance (Figure S3B). We thus examined the consequences of this reduced distance in the course of anaphase. In untreated embryos, spindle oscillations gradually built up and then died down, ultimately leaving the spindle properly aligned along the anteroposterior (AP) axis of the embryo (Figure 8A). In contrast, in *zyg-8(or484ts)* mutants, the closer proximity of the centrosome to the cell cortex pulled the entire spindle further toward the cell periphery. This appeared to prevent the spindle from re-centring during the oscillation die-down (Figure 8B). The reduction in cortical pushing disrupted the force balance, favouring excessive net pulling. As a result, the spindle failed to return to the central position by the end of anaphase, leading to both mispositioning and misorientation (Figure 8B-D). Notably, embryonic lethality of *zyg-8(or484ts)* mutants was elevated (> 99%) at the restrictive temperature. Based on these observations, we proposed that the spindle misalignment seen in *zyg-8(or484ts)* mutants arose from impaired pushing forces, which disturbed the balance between pulling and pushing forces required for proper spindle positioning. Notably, spindle mispositioning and misorientation in *zyg-8(or484ts)* mutants were restricted to anaphase, as we observed no significant differences in spindle position or orientation between mutants and untreated embryos at either mid-metaphase or anaphase onset (Figure 8C, D). These observations suggested that the anaphase-specific increase in cortical pulling forces may contribute to the observed defects.

We thus hypothesised that reducing cortical pulling force could rescue the final spindle orientation in *zyg-8(or484ts)* mutants, by preventing an excessive imbalance between cortical pulling and pushing forces. To test this, we depleted GPR-1/2^LGN^ by RNAi, a component of the cortical force generator that pulls on astral microtubules. In non-mutated embryos treated with *gpr-1/2(RNAi)* at the restrictive temperature, we observed no spindle pole oscillations, and the most-posterior position of the spindle was significantly shifted toward the cell centre, consistent with a decrease in cortical pulling forces (Figure S13A). Similarly, in *zyg-8(or484ts)* mutants treated with *gpr-1/2(RNAi)* at the restrictive temperature, oscillations were absent and the most-posterior position was also shifted closer to the cell centre (Figure S13A). Strikingly, depleting GPR-1/2 in *zyg-8(or484ts)* mutants rescued the spindle misorientation defects (Figure S13B). This finding indicated that low microtubule rigidity did not affect final spindle orientation when cortical pulling forces were reduced. These results highlighted the critical importance of maintaining a proper balance between cortical pulling and pushing forces for accurate spindle positioning and successful cell division.

Importantly, we quantified spindle final position and orientation across all experimental conditions in our study. The selective perturbation of microtubule growth via *zyg-9(RNAi)*—which induces changes in growth rates comparable to those observed in *zyg-8(or484ts)* mutants—did not affect spindle final position or orientation (Figure S4E, F). Moreover, specific impairments in microtubule nucleation (SPD-2 depletion; Figure S5E, F), microtubule stability (CLS-2 depletion; Figure S6D, E), or microtubule depolymerisation (KLP-7 depletion; Figure S7D, E) similarly failed to perturb spindle final position or orientation. Together, these observations further supported our conclusion that reduced microtubule rigidity, rather than altered microtubule dynamics, is the determining factor of spindle misorientation in *zyg-8(or484ts)* mutants.

**Figure 8:**
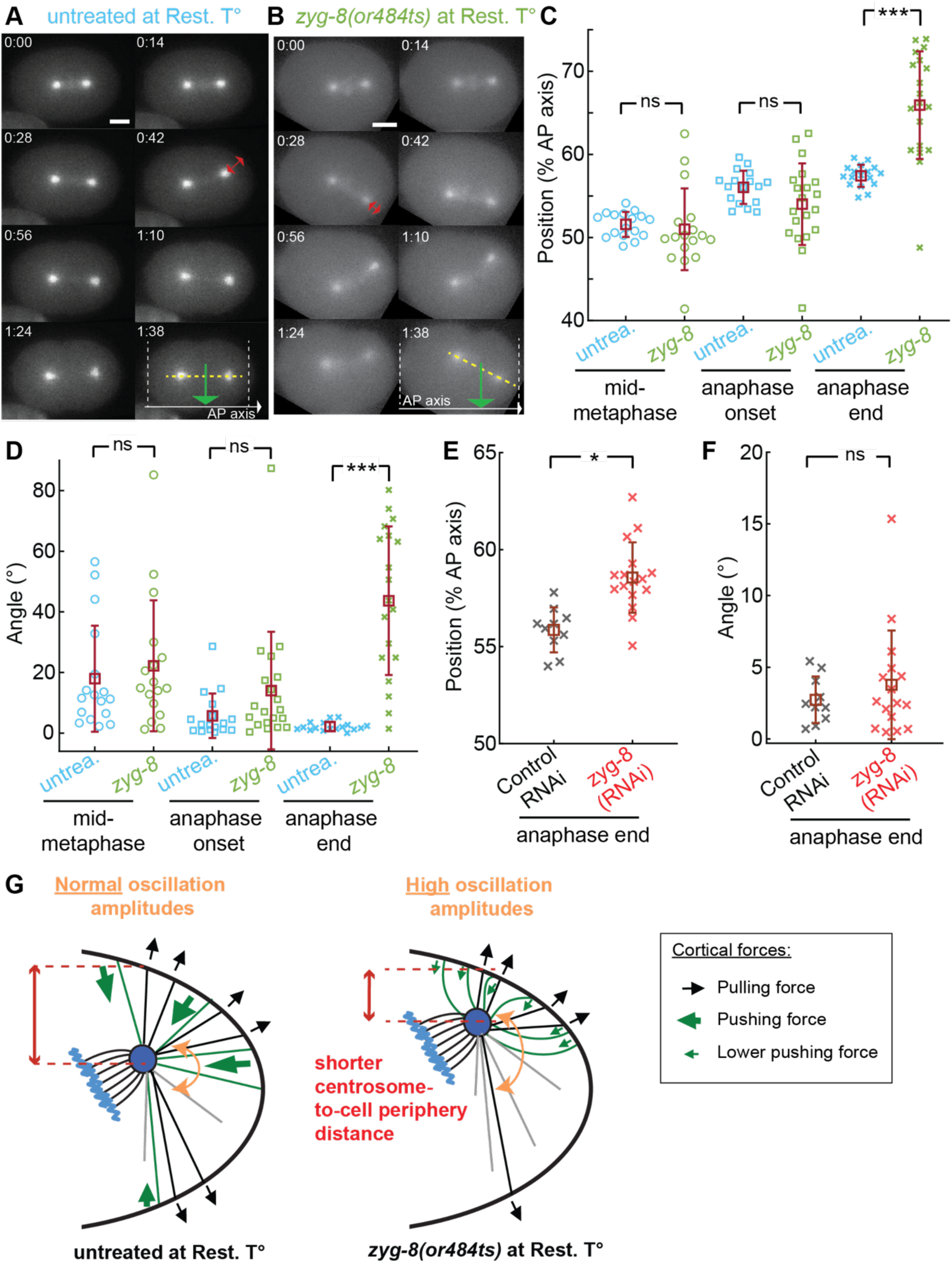
In *zyg-8(or484ts)* mutants, overly soft microtubules result in a misoriented and mispositioned spindle at the anaphase end. (**A-B**) Time-lapse series of (A) an exemplar untreated embryo and (B) an exemplar *zyg-8(or484ts)* mutant along anaphase, both at the restrictive temperature (Rest. T°) and with EBP-2::mKate2 labelling. The green arrows indicate the spindle’s final positions along the anteroposterior (AP) axis and the dotted yellow lines indicate the final orientations. The red double-headed arrows underline the minimal centrosome-to-cortex distances. The scale bars represent 10 μm. (**C-D**) Spindle (C) positions along the AP axis and (D) angles for (light green) *N* = 20 *zyg-8(or484ts)* mutants and (light blue) *N* = 17 untreated embryos, both at the restrictive temperature: (circle) at -60 s from anaphase onset, (square) at anaphase onset, and (cross) at anaphase end. (**E-F**) Spindle final (E) positions along the AP axis and (F) angles for (black) *N* = 16 *zyg-8(RNAi)*-treated embryos and (red) *N* = 10 control RNAi embryos. We tracked and analysed the centrosomes of embryos with (C-D) EBP-2::mKate2 labelling and (E-F) GFP::TBB-2 labelling (Method M7). *N* represents the total number of embryos analysed across all replicates. The brown squares and error bars correspond to the means and SD. * and *** indicate significant differences with 1x10^-3^ &< *p* ≤ 1x10^-2^ and *p* ≤ 1x10^-4^, respectively. ns denotes non-significant differences (*p* > 0.05) (Method M15). (**G**) Schematics illustrating how a large reduction in microtubule flexural rigidity, which reduces cortical pushing force, permits centrosome proximity to the cell cortex. This, in turn, causes defects in spindle positioning and orientation at the end of anaphase. Data for *zyg-8* overexpressing embryos are not included in this figure, as it focuses on the consequences of reduced microtubule rigidity—a phenotype not observed under this condition (Figure S10).

In conclusion, sufficiently rigid microtubules were required to generate the centring force necessary during anaphase, when cortical pulling forces are at their peak. This centring force prevented the centrosome from being displaced too far from the centre, ensuring proper final orientation of the spindle (Figure 8G). Reliable spindle positioning at the end of mitosis is crucial for accurate cleavage furrow placement during cytokinesis, a process that determines the proper partitioning of cell fate determinants [120].

## DISCUSSION

Based on three distinct, complementary genetic perturbations of *zyg-8*, our work highlights a dual role for ZYG-8^DCLK1^ during mitosis, involving the control of both microtubule rigidity and dynamics. We first confirmed previous findings by Srayko *et al.*, showing that ZYG-8 modestly promotes microtubule growth while limiting the nucleation of astral microtubules in *C. elegans* zygote [48] (Figures 2 and 3). However, the modest changes observed in astral microtubule dynamics in the cytoplasm upon *zyg-8* targeting, as well as its potential role in stabilising microtubules, are not sufficient to fully explain the phenotypes related to spindle pole oscillations during anaphase, spindle final position and orientation, or the altered behaviour of astral microtubule contacts at the cell cortex. If ZYG-8’s main function were to stabilise microtubules by reducing their catastrophe rate at the cell cortex, *zyg-8(RNAi)* treatment would be expected to decrease the cortical lifetimes of astral microtubules and reduce their contact counts. This phenotype would resemble what we observed with *cls-2(RNAi)* (Figure S6F, G). In contrast, we found the opposite effect: *zyg-8(RNAi)* increases both astral microtubule contact counts and durations (Figure 4C, E). Upon *klp-7(RNAi)*, which reduces microtubule depolymerisation, we found an increase in both microtubule contact counts and durations (Figure S7F, G). If ZYG-8 primarily acted to prevent microtubule depolymerization, the phenotypes would have differed markedly from those in *zyg-8(RNAi)*—yet they remained similar. Similarly, if ZYG-8 primarily promoted microtubule growth, z*yg-8(RNAi)* should result in fewer astral microtubules reaching the cortex and shorter contact durations, as previously seen with *zyg-9(RNAi)* [80]. Instead, in our current study, we observed the opposite phenotypes (Figure 4C, E-F). Furthermore, if ZYG-8 limited microtubule catastrophe within the cytoplasm, then *zyg-8(RNAi)* would be expected to decrease the number of astral microtubule contacts at the cell cortex, as more microtubules would depolymerise before reaching it. This, again, contradicts our observations (Figure 4E). By contrast, the increased number of astral microtubule contacts upon *zyg-8(RNAi)* aligns with a role for ZYG-8 in limiting microtubule nucleation. However, this effect alone cannot fully account for the observed variations in microtubule cortical-contact number. While our comparisons using depletions of various MAP provide valuable insights into the role of microtubule dynamics in spindle oscillations and cortical microtubule contact phenotypes, we acknowledge some limitations. First, some perturbations introduced additional confounding effects, e.g. *cls-2(RNAi)* affecting midzone integrity, complicating direct comparisons with *zyg-8* perturbations. Second, cumulative effects of multiple perturbations may contribute to the phenotypes observed upon *zyg-8* targeting. Nonetheless, taken collectively, our set of experiments suggests that alterations in microtubule dynamics alone cannot fully recapitulate the spindle-pole oscillation variations, spindle final position defects, and cortical microtubule contact phenotypes observed upon *zyg-8* targeting. This supports our conclusion that ZYG-8 plays an additional role beyond simply regulating microtubule dynamics.

Given that Doublecortin family proteins have been proposed to regulate microtubule rigidity, we investigated whether ZYG-8 performs a similar function in *C. elegans* zygotes. Initial evidences for ZYG-8’s role in rigidifying microtubules came from quantitative measurements of microtubule curvature. We observed increased tortuosity and curvature—both in magnitude and frequency—in *zyg-8(RNAi)* embryos and *zyg-8(or484ts)* mutants (Figures 6). The gradual differences in these curvature and tortuosity phenotypes between the two conditions aligned with ZYG-8’s binding level to microtubules (Figure S2) [81]. Furthermore, the consistency of the curvature perturbations across multiple experimental replicates (fixed and live embryos) provided strong support for ZYG-8’s role in controlling microtubule rigidity (Figures 5, 6, S9). A second line of evidence came from the agreement between our simulation predictions and experimental observations regarding the relationship between microtubule cortical lifetime and rigidity (Figures 4E, G, S12). Our DiLiPop analysis, which enables monitoring cortical microtubule force-generating events, provided a third line of evidence. It revealed that the reduction in the cortical lifetimes is associated specifically to the pushing events: less rigid microtubules bend more when growing against the cortex, thereby reducing pushing force, delaying catastrophe onset, and prolonging cortical contact lifetimes (Figure 7A, D). As a fourth argument, the altered frequency of spindle pole oscillations upon *zyg-8* targeting is consistent with a role in controlling microtubule rigidity [66, 77] (Figure 1D). Increased cortical pulling forces upon *zyg-8(RNAi)* could be thought to alternatively explain the observed increase in microtubule bending. However, such a view contradicts our DiLiPop experiments (Figure 7B, C). Additionally, cortical dynein tracking revealed that changes in dynein behaviour were insufficient to account for a large increase in pulling forces (Table S3). While we did observe increased dynein density—consistent with more microtubules reaching the cortex upon ZYG-8 depletion—this did not translate into increased pulling events, as GPR-1/2 association in the force-generating complex and dynein engagements remain the limiting factors. The increase in cortical dynein lifetimes aligned with the prolonged residency times of cortical microtubules, without suggesting an additional contribution. We conclude that ZYG-8 primarily acts by increasing microtubule rigidity rather than by regulating cortical pulling force generation.

Our data suggest that, while ZYG-8 contributes to microtubule dynamics, its primary role in regulating spindle positioning and orientation is mediated through modulation of microtubule rigidity, particularly under conditions of elevated cortical pulling forces (Figure 8A-D). This mechanical function likely represents the dominant mechanism accounting for the observed phenotypic effects, with alterations in microtubule dynamics playing a secondary role. Microtubule rigidity is more substantially reduced in *zyg-8(or484ts)* mutants compared to *zyg-8(RNAi)* embryos (Figures 5, 6, S9). This correlates with a greater decrease in centring forces (as measured by positional fluctuation assays) (Table 1). These findings align with the gradual phenotypes observed in spindle-pole oscillations, and spindle final position or orientation (Figure 1, 8). Why ZYG-8 plays a greater role in regulating microtubule rigidity rather than microtubule dynamics. This is likely because ZYG-8 and EB compete for the same microtubule binding site at the corner of four tubulin dimers [39, 121, 122]. However, EB binds preferentially to the GTP cap at growing ends [121], while Doublecortin-family proteins bind *in vitro* to the whole lattice [123]. *In vivo*, DCX was excluded from microtubule ends [124]. An exclusion of ZYG-8 from microtubule ends in *C. elegans* zygote could account for differences in the efficiency of its dual role in regulating microtubule dynamics and rigidity. This may be particularly relevant in specific cellular contexts where proteins essential for microtubule growth or nucleation (e.g., γ-tubulin ring complexes) are absent or present at low levels. In neurons, proteins from the Doublecortin family are highly expressed and regulate microtubule rigidity [37, 100]. These proteins may also contribute to microtubule nucleation and/or growth, consistent with their *in vitro* capabilities [38, 91, 92, 125]. For instance, collateral branching in hippocampal neurons was significantly delayed following DCX depletion, possibly reflecting impaired microtubule dynamics [126]. Overall, ZYG-8^DCLK1^ performs multiple microtubule-related functions in a context-dependent manner. We propose that the control of microtubule flexural rigidity represents a novel function during cell division and possibly beyond. While our data strongly implicate microtubule rigidity in spindle misorientation in *zyg-8(or484ts)* mutants, we cannot rule out the possibility of additional contributions from mild perturbations in microtubule dynamics.

Looking ahead, it will be important to explore the mechanisms by which ZYG-8 rigidifies the microtubules. Several mechanisms have been proposed, primarily in the context of neurons. Here, we draw on the body of research on the DCX protein, which shares the same two microtubule-binding domains as DCLK1^ZYG-8^. Besides, these two doublecortin domains are highly conserved in ZYG-8 [54]. (1) Microtubules can be rigidified by hindering the sliding of protofilaments through the binding of MAPs across these protofilaments. DCX binding on the microtubules is compatible with this mechanism since it reinforces lateral and longitudinal dimer couplings [38]. (2) Rigidification can also occur by bundling multiple microtubules together. Previous studies revealed such a mechanism for the Doublecortin family *in vitro* and *in vivo* [57, 90, 91, 100, 125, 127, 128]. In particular, *zyg-8(or484ts)* mutants displayed a perturbed bundle organisation of microtubules in touch receptor neurons [100]. However, in the *C. elegans* zygote, electron microscopy failed to reveal microtubule bundles in the spindle or astral microtubules close to the poles [129]. Therefore, we disfavour such a mechanism. (3) Microtubule rigidity is also regulated through post-translational modifications (PTM) [130–134]. Notably, a study showed that DCX expression increases tubulin polyglutamylation in neurons [135]. However, the *ttll-4(-);ttll-5(-);ttll-11(-)* triple mutants, which prevented tubulin glutamylation, did not show any defect in embryonic viability. It suggests that microtubule glutamylation may not significantly affect the zygote cell division [136]. (4) It was previously shown that damages along microtubule lattice soften microtubules [137, 138]. Thus, self-repair of local damages may also help restore the rigidity or stabilise the curved region [139]. Interestingly, the number of microtubule defects could notably correlate with the microtubule growth rate [140, 141]. The latter is particularly high in nematodes [142]. If a high occurrence of holes exists in the *C. elegans* embryo, ZYG-8 could contribute to healing as the Doublecortin family also acts as tubulin oligomer chaperones [92]. However, to date, there has been no evidence of microtubule lattice defects in *C. elegans* zygote. For example, embryos with EBP-2 fluorescent labelling did not show stationary fluorescent spots along the astral microtubules [48, 67]. The various mechanisms proposed above, in which ZYG-8 could play a key role in regulating microtubule flexural rigidity, are not mutually exclusive; several of them may be combined. ZYG-8 rigidifying the microtubule by coupling the protofilaments appears to us the most plausible mechanism currently.

To explore whether other microtubule rigidity regulators contribute to spindle positioning in *C. elegans* embryos, we examined two candidates: PTL-1, the sole nematode Tau homolog [117, 118], and SPD-1^PRC1^ [36]. PTL-1 depletion resulted in modest increases in oscillation amplitudes (34% anterior, 13% posterior, Figure S14A) and slight frequency decreases (9% anterior, 11% posterior, Figure S14B), alongside a 60% reduction in centring-to-damping corner frequency (Table 1). While these alterations resembled *zyg-8(RNAi)* phenotypes, their reduced magnitude likely reflects PTL-1’s limited zygotic expression [119]. Similarly to *zyg-8(RNAi)* treatment, we observed no defects in spindle final position or orientation following *ptl-1(RNAi)* (Figure S14C, D). Conversely, SPD-1 depletion produced no detectable changes in spindle oscillations (Figure S14E, F) or spindle final position and orientation (Figure S14G, H). This is consistent with SPD-1’s spindle midzone localisation and absence from astral microtubules [59, 143, 144]. These comparative analyses highlight the unique role of ZYG-8 among microtubule rigidity regulators in controlling external forces on the spindle during *C. elegans* zygote division.

Focusing on mitotic spindle positioning in the *C. elegans* zygote, the current study both complements and departs from previous investigations. Firstly, we show that controlling microtubule flexural rigidity participates in the spindle choreography. In particular, this control helps prevent the spindle poles from approaching the cell periphery too closely during anaphase (Figure 8A, B, G). Consistently, we observe increased amplitudes of spindle pole oscillations following ZYG-8 or PTL-1 depletion (Figures 1C & S14A). This control through microtubule rigidity is complementary to the well-established regulation involving microtubule dynamics [25, 67, 72, 80]. Secondly, our study underlines that cortical pushing force contributes to spindle positioning during anaphase, although in a manner distinct from that of cortical pulling forces. Cortical pulling forces are dominant, and their asymmetry, controlled by PAR polarity, ensures spindle posterior displacement [74, 78, 145–148]. To date, cortical pushing forces have been proposed to contribute to spindle centring during metaphase [25, 27, 29]. During anaphase, although restoring forces are known to be necessary for spindle oscillations, the role of cortical pushing force as a restoring force was not demonstrated [66]. Besides, it is often assumed that cortical pushing force plays a minimal role during anaphase, due to the high magnitude of cortical pulling forces and the dampening effect of microtubule buckling [68, 79, 149]. However, we report that weakening the cortical pushing forces, as observed in *zyg-8(or484ts)* mutants, hinders spindle alignment along the AP axis (Figure 8B). This force imbalance compromises the final position and orientation of the spindle (Figure 8C-D). Interestingly, decreasing cortical pulling force by depleting GPR-1/2 in *zyg-8(or484ts)* mutants rescues the spindle misorientation phenotype (Figure S13). Together, these findings highlight the importance of sufficiently strong cortical pushing forces during anaphase, which contribute to proper force balance and ensure accurate spindle orientation at the end of mitosis. Thirdly, we show that in *zyg-8(or484ts)* mutants, astral microtubules are still contacting the opposite cortex (Figure 5A, Movies S5 and S6). This rules out a model in which spindle misorientation arises from a failure of microtubules to reach the cortex, as previously suggested [54]. Overall, the astral microtubule network functions like a spring, restricting the spindle transverse motion away from the centre in *C. elegans* zygotes [25, 27, 77]. From this perspective, microtubule flexural rigidity emerges as a key physical parameter controlling the magnitude of centring forces. This mechanical contribution is essential for generating cortical pushing forces strong enough to counteract the intense cortical pulling forces that act during anaphase.

We foresee that the essential role for pushing extends beyond the nematode’s first division and may be relevant to other asymmetric divisions and maybe further. It is well established that the pulling forces dominate spindle orientation. It is accounted for by the canonical model based on the molecular machinery composed of the cortical Gα-LGN^GPR^-NuMA^LIN-5^ complex and the cytoplasmic dynein motor [10, 150]. However, less attention has been paid to the contribution of the pushing forces to spindle orientation [151]. The dependence of spindle orientation on cell shape, which is often neglected, suggests that the pushing force could play a significant role in this process [152–156]. In other systems, some studies found that the spindle did not align with the dynein-enriched cortical zones [157–160]. Furthermore, spindle orientation and centring can occur through cortical-dynein-independent mechanisms, such as those involving MARK2/Par1 kinase [29, 161]. Consequently, the traditional canonical model alone cannot account for spindle orientation in various contexts. Consistent with these observations, we propose that the cortical pushing force contributes to this phenomenon and that the flexural rigidity of microtubules must be sufficiently high. Notably, the orientation of the mitotic spindle defines the direction in which a cell divides. Therefore, many biological processes, ranging from germline stem cell division to epithelial tissue homeostasis and regeneration, require correct mitotic spindle orientation. Defects in spindle orientation have been associated with various diseases, such as organogenesis and morphogenesis failures, or cancer [162–169].

By uncovering ZYG-8’s dual role in *C. elegans* mitosis, we can now question its contribution to zygote meiosis, especially through controlling microtubule flexural rigidity. A recent study highlighted that ZYG-8 promotes meiotic spindle stability and suggested that it may regulate microtubule dynamics and motor-driven forces [56]. We wonder whether some of the perturbations that the authors observed upon *zyg-8* targeting could be attributed to the deregulation of microtubule rigidity. Notably, they found that most *zyg-8(or484ts)* meiotic spindles were significantly bent at the restrictive temperature. Additionally, some images of the mutant showed microtubules exhibiting high local curvatures. Similar perturbations were also observed following ZYG-8 depletion using the Auxin-inducible degradation system. In light of these observations, it is plausible that ZYG-8 contributes to the assembly or/and stability of meiotic spindles in the zygote by stiffening microtubules.

In conclusion, our findings regarding the pivotal role of ZYG-8^DCLK1^ in spindle positioning and orientation, primarily through rigidifying microtubules during mitosis, offer not only a deeper understanding of the fundamental mechanisms governing cell division but also highlight significant implications for cancer therapy. Given that accurate spindle positioning is essential for maintaining the balance between cell proliferation and differentiation, our results may have broader relevance for understanding how disruptions in these processes contribute to tumour initiation and progression [170–172]. The frequent deregulation of DCLK1 in various solid tumours (e.g., those from the colon, pancreas, kidney and breast) suggests its involvement in a common process such as cell division [173–176]. Additionally, the proteins of the Tau family are also found to be deregulated or mutated in cancers [177–179]. While our study focuses on fundamental mechanisms, these observations raise the possibility that microtubule rigidity could represent an underexplored aspect of cancer cell biology.

## MATERIAL AND METHODS

### M1 C. elegans strains and their culturing

The *C. elegans* strains used in the present study are listed in Table S1. *C. elegans* nematodes were cultured as described in [180] and dissected to obtain embryos. The strains were maintained at 20°C, except the thermosensitive mutant *zyg-8(or484ts)* maintained at 15°C and AZ244 maintained at 25°C to increase transgene expression. The strains were handled on nematode growth medium (NGM) plates and fed with OP50 *E. coli* bacteria. The approval number for the use of geneJcally modified organisms (GMOs) in this study is 6674.

### M2 RNAi feeding

RNA interference (RNAi) experiments were performed by feeding, as described in [181]. Bacteria were obtained by Source BioScience (*zyg-8*: III-6C10; *ptl-1*: III-1A12; *spd-2*: I-4O08; *zyg-9*: II-6M11; *cls-2*: III-4J10; *klp-7*: III-5B24; *gpr-1/2*: III-4J09; *gpb-1*: II-8A05; *spd-1*: I-7D17). The feedings were performed at 20°C, except for *gpr-1/2* realised at 15°C to be at the restrictive temperature of *zyg-8(or484ts)* mutant. For RNAi targeting of *gpr-1/2, zyg-8* and *ptl-1*, the treatment lasted 96 hours and dsRNA expression was induced with 4 mM of IPTG (Isopropyl β-D-1-thiogalactopyranoside). For *spd-1*, *spd-2*, *zyg-9*, *klp-7* and *cls-2*, dsRNA expression was induced with 3 mM IPTG, and for *gpb-1* with 1 mM IPTG. The RNAi treatment duration was 6 hours for *zyg-9* and *spd-2*, 24 hours for *klp-7* and *gpb-1*, and 48 hours for *cls-2* and *spd-1*. The control embryos for the RNAi experiments were fed with bacteria carrying the empty plasmid L4440.

### M3 Preparation of the embryos for live imaging or immunofluorescence

<u>For live imaging</u>, embryos were dissected in M9 buffer (prepared by combining 3 g KH_2_PO_4_, 6 g Na_2_HPO4, 5 g NaCl and 1 mL of 1 M MgSO_4_ diluted in ultrapure water to 1 L) and mounted on a pad (2% w/v agarose, 0.6% w/v NaCl, 4% w/v sucrose) between a slide and a coverslip.

<u>For immunofluorescence staining</u>, embryos were dissected in M9 buffer and mounted on a slide coated with poly-L-lysine (0.1%) (Sigma-Aldrich P1524). The slides were dipped in liquid nitrogen to crack the embryo’s eggshell and immediately transferred to -20°C methanol for fixation for 20 minutes (freeze-cracking method [182]). They were then rinsed for 10 minutes in PBS 1X before blocking for 20 minutes in PBS-Tween (0.2%) with BSA (1%). Importantly, for the thermosensitive *zyg-8(or484ts)* mutants, we transferred them as L4 larvae to 25°C and incubated for 15 hours before use, as previously done in [54].

### M4 Fluorescence microscopic imaging conditions

The various microscopy setups used are presented in the table below. Embryos were imaged at 23°C, except otherwise stated. Images were stored using the Omero software [183].

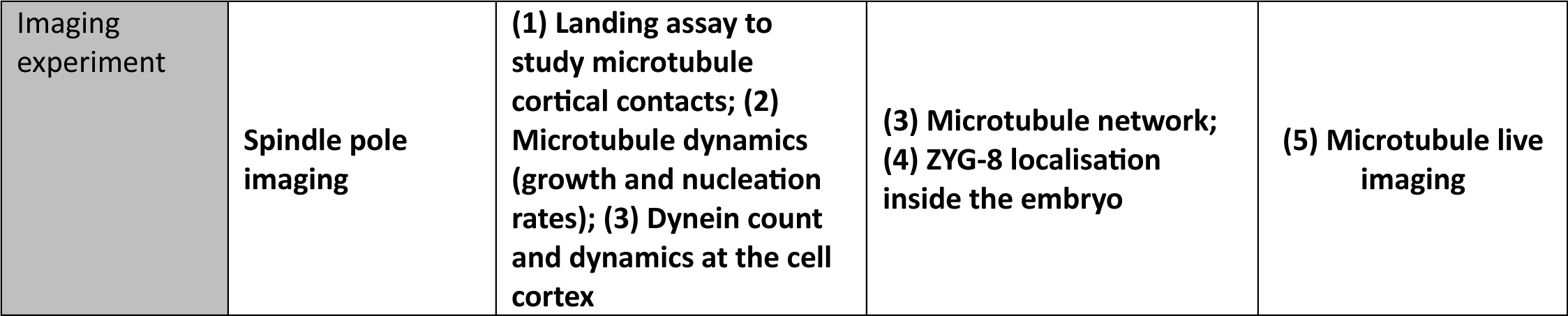
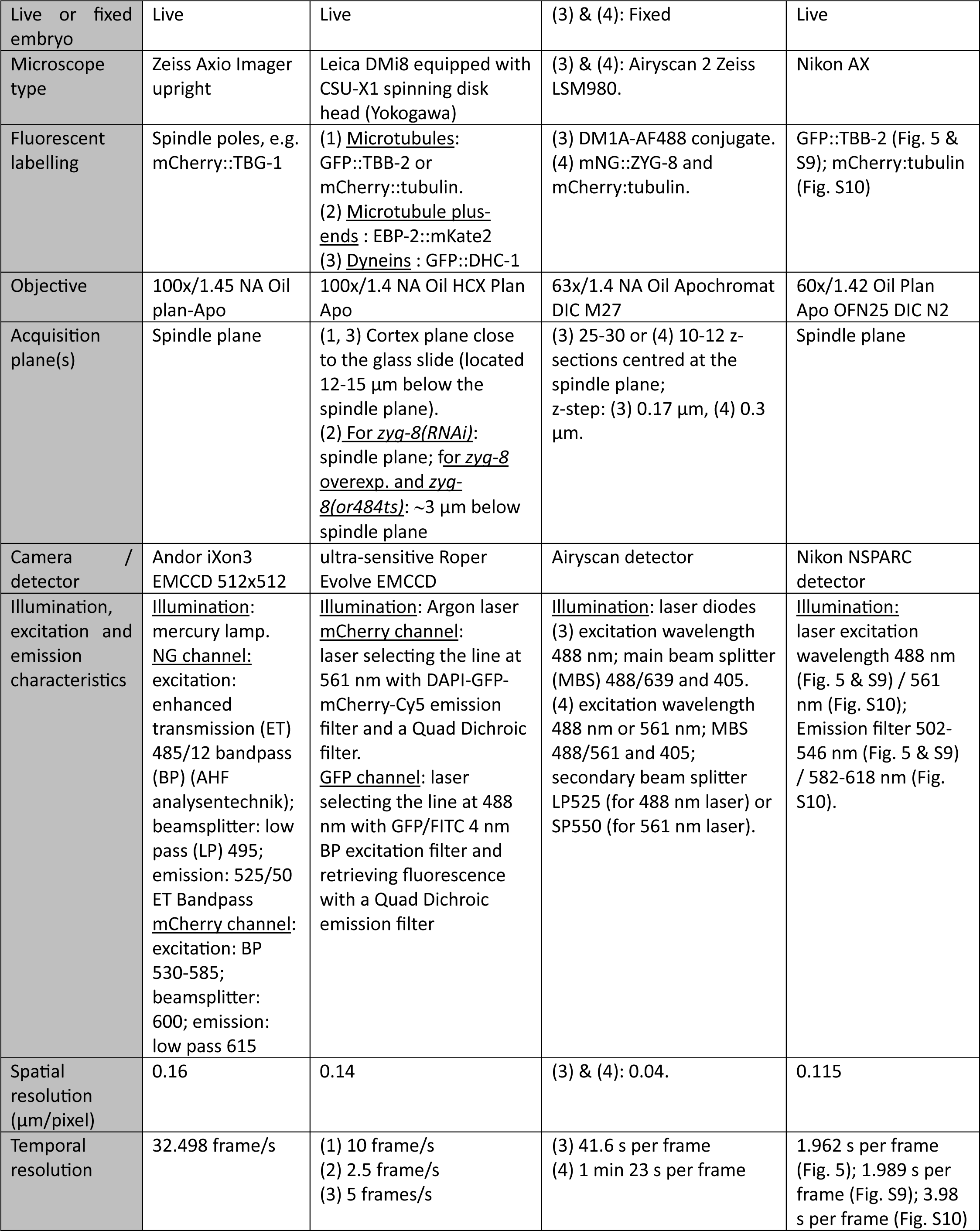
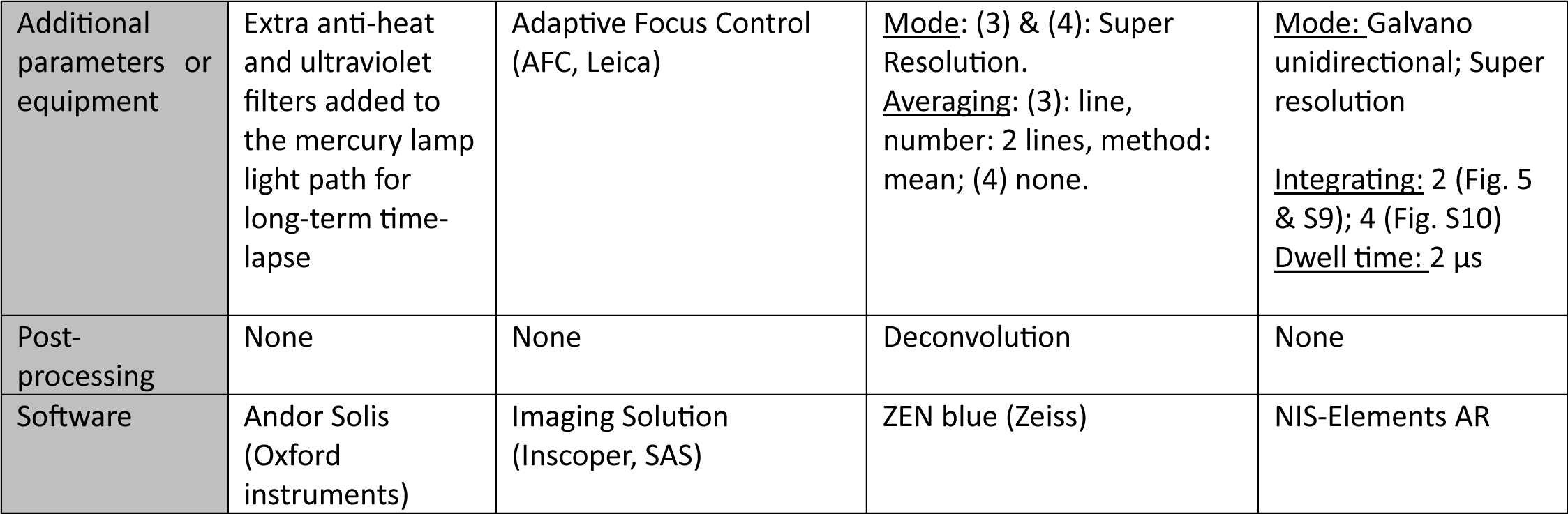

|  |  |  |  |  |
| --- | --- | --- | --- | --- |
| Imaging experiment | Spindle pole imaging | (1) Landing assay to study microtubule cortical contacts; (2) Microtubule dynamics (growth and nucleation rates); (3) Dynein count and dynamics at the cell cortex | (3) Microtubule network; (4) ZYG-8 localisation inside the embryo | (5) Microtubule live imaging |
| Live or fixed embryo | Live | Live | (3) & (4): Fixed | Live |
| Microscope type | Zeiss Axio Imager upright | Leica DMI8 equipped with CSU-X1 spinning disk head (Yokogawa) | (3) & (4): Airyscan 2 Zeiss LSM980. | Nikon AX |
| Fluorescent labelling | Spindle poles, e.g. mCherry::TBG-1 | (1) <u>Microtubules</u>: GFP::TBB-2 or mCherry::tubulin.<br>(2) <u>Microtubule plus-ends</u> : EBP-2::mKate2<br>(3) <u>Dyneins</u> : GFP::DHC-1 | (3) DM1A-AF488 conjugate.<br>(4) mNG::ZYG-8 and mCherry:tubulin. | GFP::TBB-2 (Fig. 5 & S9); mCherry:tubulin (Fig. S10) |
| Objective | 100x/1.45 NA Oil plan-Apo | 100x/1.4 NA Oil HCX Plan Apo | 63x/1.4 NA Oil Apochromat DIC M27 | 60x/1.42 Oil Plan Apo OFN25 DIC N2 |
| Acquisition plane(s) | Spindle plane | (1, 3) Cortex plane close to the glass slide (located 12-15 μm below the spindle plane).<br>(2) <u>For *zyg-8(RNAi)*</u>: spindle plane; <u>for *zyg-8 overexp. and zyg-8(or484ts)*</u>: ~3 μm below spindle plane | (3) 25-30 or (4) 10-12 z-sections centred at the spindle plane;<br>z-step: (3) 0.17 μm, (4) 0.3 μm. | Spindle plane |
| Camera / detector | Andor iXon3 EMCCD 512x512 | ultra-sensitive Roper Evolve EMCCD | Airyscan detector | Nikon NSPARC detector |
| Illumination, excitation and emission characteristics | <u>Illumination</u>: mercury lamp.<br><u>NG channel</u>: excitation: enhanced transmission (ET) 485/12 bandpass (BP) (AHF analysentechnik); beamsplitter: low pass (LP) 495; emission: 525/50 ET Bandpass<br><u>mCherry channel</u>: excitation: BP 530-585; beamsplitter: 600; emission: low pass 615 | <u>Illumination</u>: Argon laser<br><u>mCherry channel</u>: laser selecting the line at 561 nm with DAPI-GFP-mCherry-Cy5 emission filter and a Quad Dichroic filter.<br><u>GFP channel</u>: laser selecting the line at 488 nm with GFP/FITC 4 nm BP excitation filter and retrieving fluorescence with a Quad Dichroic emission filter | <u>Illumination</u>: laser diodes<br>(3) excitation wavelength 488 nm; main beam splitter (MBS) 488/639 and 405.<br>(4) excitation wavelength 488 nm or 561 nm; MBS 488/561 and 405; secondary beam splitter LP525 (for 488 nm laser) or SP550 (for 561 nm laser). | <u>Illumination</u>: laser excitation wavelength 488 nm (Fig. 5 & S9) / 561 nm (Fig. S10); Emission filter 502-546 nm (Fig. 5 & S9) / 582-618 nm (Fig. S10). |
| Spatial resolution (μm/pixel) | 0.16 | 0.14 | (3) & (4): 0.04. | 0.115 |
| Temporal resolution | 32.498 frame/s | (1) 10 frame/s<br>(2) 2.5 frame/s<br>(3) 5 frames/s | (3) 41.6 s per frame<br>(4) 1 min 23 s per frame | 1.962 s per frame (Fig. 5); 1.989 s per frame (Fig. S9); 3.98 s per frame (Fig. S10) |
| Additional parameters or equipment | Extra anti-heat and ultraviolet filters added to the mercury lamp light path for long-term time-lapse | Adaptive Focus Control (AFC, Leica) | Mode: (3) & (4): Super Resolution.<br>Averaging: (3): line, number: 2 lines, method: mean; (4) none. | Mode: Galvano unidirectional; Super resolution<br>Integrating: 2 (Fig. 5 & S9); 4 (Fig. S10)<br>Dwell time: 2 μs |
| Post-processing | None | None | Deconvolution | None |
| Software | Andor Solis (Oxford instruments) | Imaging Solution (Inscoper, SAS) | ZEN blue (Zeiss) | NIS-Elements AR |

### M5 Localisation of ZYG-8 in the C. elegans one-cell embryo

To study ZYG-8 localisation in the zygote, we used *C. elegans* strains expressing fluorescently labelled microtubules (mCherry::tubulin) and fluorescently labelled ZYG-8 (mNG::ZYG-8) (Figures S1A-C and S1G-I). We analysed deconvolved dual-colour images of fixed embryos acquired using the Zeiss Airyscan 2 microscope. First, we computed the intensity profiles of both mCherry::tubulin and mNG::ZYG-8 signals along a 5-pixel wide line (Figures S1D-E, J-K). The lines used for this analysis are shown in Figure S1D, J. Co-localisation of intensity peaks indicated the binding of ZYG-8 to microtubules. Second, we investigated pixel-wise co-localisation by producing a scatterplot and computed Pearson’s correlation coefficient *r* (Figure S1F, L). Statistical significance was assessed through the Costes statistical test using 200 rounds of randomisation and a bin width of 0.001 [184]. These analyses were performed using Fiji and the JACoP plugin [185]. To test ZYG-8 localisation specifically on astral microtubules, we computed the Pearson’s coefficient after masking the spindle area in the images.

### M6 Quantification of ZYG-8 level at the worm and embryo scales

To quantify the ZYG-8 level at the worm scale, we performed a Western blot analysis using protein extracts obtained from different *C. elegans* strains (Figure S2A-B). For each experiment, 30 adult hermaphrodites were picked into 200 µL of M9 buffer and snap-frozen in liquid nitrogen, then thawed, mixed with 6.5 µL loading buffer (DTT 0.1 M), and loaded onto a Mini Protean TGX 4–20% Tris-Glicine gel from Bio-Rad (Hercules, CA). Immunoblots were probed using the primary anti-OLLAS antibody (Novus Biologicals NBP1-06713, 1 mg/mL, host: rat) diluted at 1/2000 and the primary anti-tubulin antibody (Sigma-Aldrich T5168, host: mouse) diluted at 1/2000, and then revealed using an HRP (horseradish peroxidase) conjugated secondary antibody (1:20000; Jackson ImmunoResearch, West Grove, PA; host: rat for antiOLLAS & mouse for anti-tubulin). Membrane saturation and all antibody dilutions were made in PBS–Tween 0.2% and 5% non-fat dry milk. Blots were incubated with WesternBright ECL spray (Advansta K-12049-D50) before detecting by chemiluminescence on an Amersham Imager 680 (GeHealthcare, Chicago, IL). Quantification was performed using Fiji [186].

To complement the first quantification, we analysed deconvolved dual-colour images of embryos carrying two fluorescent tags: mCherry::tubulin and mNG::ZYG-8. For each embryo, we realised, in the same regions of interest (ROI), red and green fluorescence intensity measurements (mean within ROIs) using Fiji: 2 square ROIs (40 x 40 pixels; 1.6 x 1.6 μm) within the spindle (1 per half-spindle), 3 square ROIs (40 x 40 pixels) within the cytoplasm, 5 curvilinear ROIs on astral microtubules using the segmented line tool, and a ROI (40 x 40 pixels) outside the embryo, the latter to measure the background fluorescence. Exemplar ROIs are highlighted in Figure S2C. For spindle and cytoplasm measurements, fluorescence quantification was performed on maximum projected images (3 z-sections centred on the spindle mid-plane). After subtracting the background fluorescence (mean intensity in corresponding ROI) from each fluorescence of interest (spindle, cytoplasm and astral microtubule) and for each channel separately, we computed the ratio of ZYG-8 fluorescence over tubulin fluorescence for the three measurements (Figure S2G, H).

### M7 Centrosome-tracking assay and oscillation characterisation

The tracking of labelled centrosomes (e.g., mCherry::TBG-1) and the analysis of their trajectories were conducted using custom tracking software developed using Matlab (The MathWorks) [66]. Tracking of γ-tubulin labelled embryos fixed in -20°C methanol demonstrated an accuracy of 10 nm [25]. To determine embryo orientations and centres, we performed cross-correlation of the embryo background cytoplasmic fluorescence with artificial binary images that represented an embryo. We calculated the embryo size (lengths of the long and short axes) using the active contour algorithm [187]. From the spindle pole positions, we measured the maximal amplitudes of anaphase oscillations (peak-to-peak) for both poles, normalised to the length of embryo’s short axis, as well as the frequencies of these maximal oscillations (e.g., Figures 1 and S3). We also assessed mitotic spindle length, normalised to the length of embryo’s long axis, and the maximal spindle elongation rate following nuclear envelope breakdown (e.g., Figure S7A). Finally, we quantified the spindle position (the midpoint between the two centrosomes), and the spindle orientation (the angle between the spindle axis and the embryo’s long axis) at the end of mitosis, specifically at 150 s after the anaphase onset (e.g., Figure 8C-D).

### M8 Characterisation of astral microtubule growing and nucleation rates

We labelled microtubule plus-ends using EBP-2::mKate2, which form comet-like structures as the microtubules grow (Figures 2A and 3A). We performed an analysis of the comets adapted from the article by Srayko *et al.* using Fiji (except otherwise cited) [48]. We measured microtubule dynamics during metaphase using single-focal-plane imaging.

<u>To measure the growth rates</u>, we first denoised the movies using Kalman filtering, with the gain set to 0.5 and the initial noise estimate set to 0.05 [188] (Figure 2A). Next, we traced a line along the comet trajectory in the pre-processed movie, restricting to comets that remained visible in the focal (XY) plane for at least 10 frames (4 s) to exclude comets growing in the three dimensions (XYZ). We generated a kymograph from the traced line to assess comet displacement velocity. We analysed ∼50 comets per condition (distributed across ∼10 embryos, with ∼5 comets analysed per embryo), ensuring balanced sampling from both anterior and posterior centrosomes (∼25 comets per centrosome) when possible.

<u>To measure the nucleation rate</u>, we first denoised the movies using Candle filtering (parameters: *beta* = 0.1; *patchRadius* = 1; *searchRadius* = 3; *back* = 1) (Figure 3A). We did not apply additional pre-processing for *zyg-8(RNAi)* treatment, *zyg-8* overexpression and their control. However, for *zyg-8(or484ts)* mutants and untreated embryos, we enhanced contrast using an unsharp mask (*radius* = 1; *Mask weight* = 0.7). Since centrosomes moved during metaphase, we applied Icy’s rigid registration plugin (v1.4) on the denoised images [189]. This registration aligns each image with a reference image to correct for centrosome displacements, as described in [190]. Doing so, we accurately counted the comets crossing a half-circle region of interest (ROI) positioned 9 µm from the centrosome after [48]. We generated kymographs over 150 frames (60 seconds). To address the inhomogeneous background in the raw kymographs, we applied background subtraction using Gaussian blurring. We then set an intensity threshold to retain the brightest 2% of pixels in *zyg-8(RNAi)* and control kymographs, and 2.5% for *zyg-8* overexpression ones, *zyg-8(or484ts)*, and their controls. Finally, we used the “Analyse particles” command and counted objects considered as individual EBP-2 comets (particle size: 2 pixels to infinity, 0.14 μm/pixel). Measurements were done for both anterior and posterior centrosomes, except when only a single centrosome was clearly visible.

### M9 Centrosome-size measurements

We analysed images of embryos labelled with either mCherry::TBG-1 and mCherry::HIS-58, or GFP::TBG-1 and GFP::HIS-11, by tracking their centrosomes (cf. M7). Based on the tracked positions, we extracted vignettes of 6.4 x 6.4 µm (40 x 40 pixels; Figure 3E, F). We then performed a 2D Gaussian least-square fit to characterise both the posterior and anterior centrosomes every second [191]. From these fits, we obtained the *x*- and *y*-widths and calculated the diameter of the centrosomes, assuming they are round, using the formula: 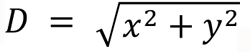.

### M10 Landing assay to study astral microtubule cortical contacts

The entire microtubule lattices were fluorescently labelled using GFP::TBB-2, YFP::TBA-2, or mCherry::tubulin to view them in all their states. We used the same approach as the one previously implemented in [80] and available on Zenodo (doi: 10.5281/zenodo.4552485); we combined a Kalman filtering [188] (gain set to 0.5 and initial noise to 0.05) and *u-track* [192] whose parameters are listed in the table below. Movies S1–S4 show representative examples of microtubule contact tracking under the four conditions we studied. After tracking the microtubule contacts, we analysed them using a two-step approach to examine distinct aspects of microtubule-cortical interactions.

1. <u>Basic quantification</u>. We quantified the number of microtubule cortical contacts per frame, averaged them over a 10-second window, and normalised them by the embryo area. This provided an instantaneous count of microtubule contacts throughout mitosis for each embryo, which we then averaged across embryos within each experimental condition (e.g., Figure 4C). We also computed the duration histograms of astral microtubule contacts per embryo and averaged these histograms across embryos for each condition (e.g., Figure 4E). This analysis assessed whether ZYG-8 contributes to microtubule stabilisation by preventing e.g. cytoplasmic depolymerisation or cortical catastrophe events.
2. <u>DiLiPop (Distinct Lifetime Population) analysis</u>. This statistical analysis of the microtubule cortical contact durations reveals the short-lived and long-lived populations of astral microtubules at the cortex associated with pulling and pushing events, respectively [80]. This analysis evaluated ZYG-8’s role in cortical force generation. By globally fitting the embryo-set duration distributions of a given condition, we obtained the characteristic lifetimes and densities of the cortical pulling and pushing events (e.g., Figure 7B-E). Error bars of the lifetimes and densities were obtained by bootstrapping as explained in [80]. These densities of events are distinct from instantaneous contact density and thus independent of the cortical contact durations (except in the case of anaphase pulling events due to all force generators being involved in pulling). We noticed that the lifetimes of pulling and pushing events in the present study differed from those in our previous work [80]. This discrepancy likely stemmed from differences in microtubule labelling (GFP::TBB-2 vs. YFP::TBA-2) and variations in microscope sensitivity. Therefore, we consistently compared results from experiments conducted within the same time frame (within two months).

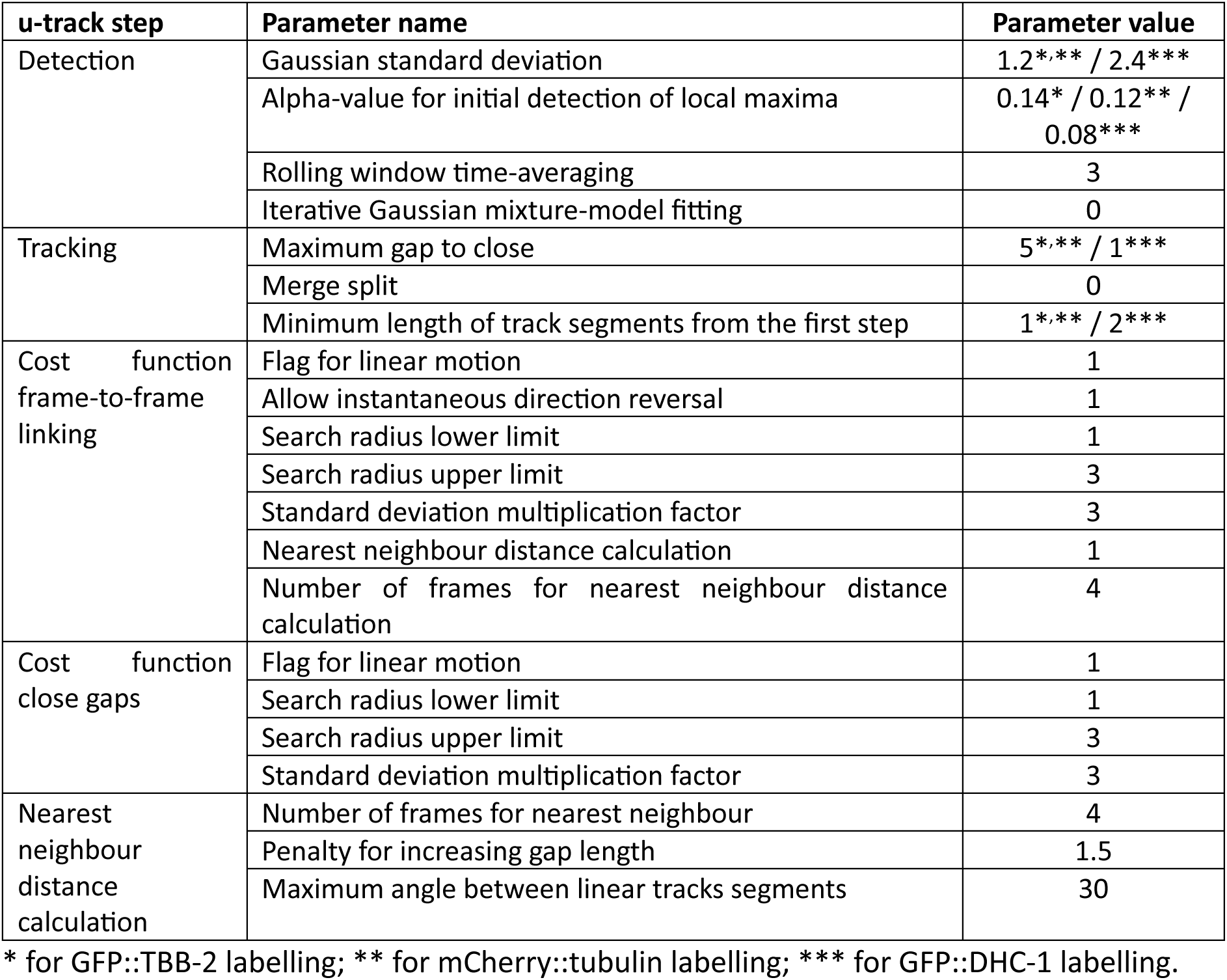

### M11 Mapping astral microtubules and characterising their local curvatures and their tortuosity

We stained microtubules using an anti-α-tubulin antibody, clone DM1A raised in mouse, Alexa Fluor 488 conjugate (Sigma-Aldrich 16-232), diluted at 1/500 in PBS-Tween (0.2%) with BSA (1%). It targets several ɑ-tubulin isoforms in various species. Antibody incubation was performed overnight at 4°C. Slides were then rinsed for 10 minutes in PBS-Tween (0.2%), twice, in a light-free environment. We could then visualise microtubules all along their length. We identified the z-section(s) as the more appropriate for astral microtubule segmentation. For most embryos, we found a single z-section per embryo for which both centrosome asters were visible. For some embryos having the two centrosomes on distinct z-sections, we considered two planes of interest.

<u>Five-step image processing pipeline</u> to map the local curvatures of the astral microtubules (Figure S8D). **(1)** We used the planes above and below the plane of interest to perform an extended depth of field projecting the 3 z-sections into a single image (Figure S8E) [193]. We thus better visualise microtubules all along their length. **(2)** We applied the *Orientation Filter Transform* (OFT, number of orientations = 20; radius = 5 pixels) since it is effective in enhancing filamentous pattern against noises [20, 194] (Figure S8F). **(3)** We then used the interactive machine-learning tool, *Ilastik,* to segment the astral microtubules (Figure S8G). The dedicated paragraph below provides more detailed information about our segmentation. **(4)** We generated the skeleton of the filament traces from the binary segmentation, taking advantage of *SOAX* (Figure S8H). Further information about the use of SOAX is available in a dedicated paragraph below. **(5)** We finally measured the local curvatures of astral microtubules pixel-by-pixel from the skeleton coordinates, enabling mapping of the local curvatures (Figure S8I). For a set of three neighbouring points sampling the curve, the local curvature corresponds to the inverse of the radius of the circle going through the mid-point and best fitting the two others. Local curvatures were computed along microtubules according to the three-point method [195] using a code retrieved on Matlab Central [196]. In a nutshell, microtubules are viewed as parametrised curves; at each point, the *x* and *y* coordinates are represented as second-order polynomials of the curvilinear abscissa passing through the point and its two neighbours. The curvature is then computed with the Frenet-Serret formula.

*<u>Ilastik</u>*: We initially trained *Ilastik* using the conditions *zyg-8(RNAi)* and its control RNAi during metaphase. This involved approximately 10 manual annotations of microtubules across three embryos for each condition to cover various intensities, contrasts and microtubule shapes. We then used this trained model, referred to as #1M (M for metaphase), to segment, in a semi-supervised way, the astral microtubules in metaphase images. Next, we trained the model to segment anaphase images of *zyg-8(RNAi)*-treated embryos and their controls. Indeed, denser microtubule networks in anaphase required additional annotations. This led to the upgraded model #1A that we used to segment anaphase images. We then analysed *zyg-8(or484ts)* mutants and untreated embryos in metaphase. The model was further improved by manually annotating three mutants, resulting in model #2M, which we used to segment images of both mutants and untreated embryos in metaphase. We applied a similar process for segmenting anaphase images of mutants and untreated embryos, enhancing model #2M with a few annotations of anaphase mutants to get the model #2A. The table below summarised the *Ilastik* models used to produce the data displayed in the different figures.

| ***Ilastik* model** | **Used for data displayed in figures** |
| --- | --- |
| #1M | 6A; 6E & 6G (only *zyg-8(RNAi)* and its control RNAi) |
| #1A | 6B; 6F & 6H (only *zyg-8(RNAi)* and its control RNAi) |
| #2M | 6C; 6E & 6G (only *zyg-8* mutant and untreated embryo at Rest. T°) |
| #2A | 6D; 6F & 6H (only *zyg-8* mutant and untreated embryo at Rest. T°) |

*<u>SOAX</u>*: We extracted the centrelines from the thick filaments utilising SOAX mostly with standard parameters. We made the few specific adjustments highlighted in the table below. These five adjusted parameters were crucial for accurately delineating the curved regions of the microtubules. The settings used ensured that the centrelines faithfully represented the shapes of the microtubules, as confirmed through visual validation. Additionally, manual corrections could be applied at this stage, especially when two microtubules were mistakenly merged into one.

| ***SOAX* parameter name** | **Parameter value** |
| --- | --- |
| Snake Point Spacing | 4 |
| Beta | 0.01 |
| Stretch Factor (k_str) | 0.8 |
| Radial Near (pixels) | 8 |
| Radial Far (pixels) | 16 |

<u>Microtubule shape quantification</u>: We analysed only the astral microtubules that were at least 4 μm long after segmentation, as shorter ones often represented fragments. This approach was appropriate for comparing conditions without seeking absolute quantification. We conducted three complementary approaches.

1. For each z-projected stack, we calculated the normalised histogram of local curvatures with a bin size of 0.05 μm^-1^ (Figures S8A & S8J) and averaged these histograms per condition (Figure 6A-D). The curvatures, expressed in per micrometre, range from 0.025 to 1 μm^-1^, corresponding to curvature radii from 40 μm to 1 μm, respectively. 40 µm could be interpreted as mostly straight astral microtubules.
2. We computed the 95^th^ percentile curvature for each microtubule to capture faithfully the highest curvatures (Figure S8B) and then calculated the median of these values across all microtubules in each z-projected stack. Each stack exhibited 15 to 74 microtubules. We finally compared the set of medians between two conditions (Figure 6E, F).
3. We measured the tortuosity for each microtubule, defined as the curvilinear distance divided by the end-to-end distance (Figure S8C). Straight microtubules have tortuosity values near one, while bent microtubules show higher values. Then, we calculated the medians of tortuosity values across all microtubules in each z-projected stack, and compared the set of medians between two conditions (Figure 6G, H).

### M12 Simulation using Cytosim

We used *Cytosim* to simulate cytoskeletal filaments [110]. We modified *Cytosim* to allow multiple fibre types to emanate from a single aster and to exert a given external force on solids, fibres or spheres (doi: 10.5281/zenodo.14246919). We conducted a 2D simulation of the first mitosis of the *C. elegans* zygote, focussing on spindle positioning and the astral microtubule network (Figure S12A). The simulation represented the spindle as a rigid rod implemented with static spindle microtubules, excluding any dynamic elements such as motor proteins. The parameters used in the simulation are detailed in Table S2 and the simulation files are provided as supplemental files S1, S2 and S3 with an exemplar video for each microtubule rigidity (Movies S12-S14). Importantly, to mimic cortical asymmetric pulling forces, we applied external forces at the spindle poles: -180 pN at the anterior pole and +300 pN at the posterior pole [78]. We used solids to simulate the centrosomes and attached two types of fibres to each, to differentiate between spindle and astral microtubules. The astral microtubules were defined as fibres without dynamics at their minus ends and with plus-ends displaying classic dynamic instability. We set their total number to 75, according to our estimated count of astral microtubules longer than about 2 µm in a 2D plane. Their rigidities were set to 2, 10, or 25 pN·µm², reflecting the range of *in vitro* measurements, varying between 3.7 and 26 pN.μm^2^ [21], due to the current lack of *in vivo* rigidity values. While the actual rigidity in *C. elegans* zygotes may differ due to the presence of MAPs, tubulin specificity, and cellular environment factors, we aimed at linking relative rigidity variations and microtubule cortical residency to support our conclusions. All microtubule dynamic parameters were kept constant across simulations. Additionally, these fibres could engage with cortical force generators simulated as hands bound to the cortex.

We measured both the highest curvatures (95^th^ percentile) and the tortuosity of the astral microtubules (Figure S12B-C) after 45 seconds of simulation, following the approach outlined in method M11. To estimate the durations of fibre cortical contacts, we analysed fibre lengths, end positions, and cortical anchor positions. When microtubules push against the cortex, they grow slowly due to the force-velocity relationship, remaining near the cell cortex. We first recovered fibre lengths over time to identify instances of very low growth rates (under 0.18 μm/s), indicating moments when the fibres were stalled (neither growing nor shrinking). Next, we filtered out stalled fibre ends further than 1 μm from the cortex. We allowed 15 s for warming-up and considered cortical contacts during the next 45 s to compute the duration histogram. We then fitted it with a mono-exponential model [67, 197]. This analysis yielded a characteristic lifetime for each simulation (Figure S12D). We realised 40 simulations per condition.

### M13 Studying cortical dyneins

To study cortical dynein behaviour, we used a strain expressing GFP::DHC-1 [114]. Movies acquired at the cell cortex were first denoised using the Candle filter (ß = 0.05, patch radius = 1, search radius = 3, background = 1) [198], followed by dynein spot tracking in uTrack using the parameters listed in the table from M10. Movie S15 shows a representative example of dynein tracking in a control RNAi embryo.

To characterise the dynamic behaviour of cortical dynein, we fitted duration histograms with either a mono- or a double-exponential model to determine whether the cortical dynein population is homogeneous or comprises two distinct populations. For robust analysis, we included only tracks lasting ≥ 0.6 s (3 frames) at the cell cortex to exclude noisy, short-lived signals.

### M14 Fourier analysis of the positional micromovements

We investigated the micromovements of spindle position along the transverse axis over time, based on our previously published method [25]. From the time series of the spindle position along the transverse axis during anaphase obtained by tracking the poles (cf. M7), we computed the power spectral density (*PSD*) of the signal multiplied by the Hann window to safeguard against artefact due to the position difference between the beginning and end of the considered time interval [199]. It contrasted with [25], where we selected only embryos with low drift in their position over time, rather than windowing. The Hann windows appeared to be a sensible choice for fitting the *PSD* later since its discrete Fourier transform has only three non-zero coefficients [199, 200]. Therefore, once convolved to the real *PSD* of the spindle length, it minimally “blurs” it.

We then fitted this *PSD* with a second-order Lorentzian model, as in [25]. Along a mechanistic line, this model corresponds to a Kelvin-Voigt-with-inertia discrete model corresponding to a Hookean spring (centring, *K*), a dashpot (damping, *G*), and an inertial element (*I*) in parallel. However, these quantities, when retrieved by fitting, depend on the amplitude of the stochastic noise that powers the fluctuations. We, therefore, utilised an alternative representation featuring (1) a diffusion coefficient *D*, inversely proportional to the damping *G* and dependent on the stochastic noise powering the fluctuations [25], (2) the centring-to-damping corner frequency *f_c_* (∝*K*/*G*) reflecting centring efficiency, and (3) the damping-to-inertia corner frequency *f_0_* (∝*G*/*I*); the inertia *I* arising from load-dependent detachment of force generators as for anaphase rocking [66]. Such a model was transformed into the Fourier space and convolved on the fly with the transform of the Hann window to perform the fit [199]. We then performed the fit using augmented Lagrangians and refined it using the pattern search algorithm [201, 202] as implemented in [199]. To quantify the variability from biological origin of the spectrum over *N* embryos, we estimated the confidence interval at significance 0.05 using bootstrap with bias-corrected and accelerated percentile method [203, 204]. In particular, it involved performing 30 iterations of fitting a set of *N* embryos drawn with replacement from the available ones.

### M15 Experimental replication and Statistical analysis

Each experimental condition was examined across multiple biological replicates (at least two independent experimental sessions) and each embryo was treated as an independent biological sample, consistent with established practices in *C. elegans* research. The total sample size for each condition, denoted *N*, represents pooled data from these replicate sessions. The high reproducibility of our worm cultivation and treatment protocols, microscopy acquisition parameters and phenotypic measurements across replicates justifies pooling data from different sessions.

Statistical significance was assessed on embryo-level measurements. The difference between two samples was assessed by the two-tailed Student’s *t*-test with Welch–Satterthwaite correction for unequal variance. When Gaussian distribution cannot be assumed, we used the Mann–Whitney test (also known as the Wilcoxon rank-sum test). We used it for instance to compare microtubule highest curvatures (e.g., Figure 6E) or microtubule tortuosity (e.g., Figure 6G). For the sake of simplicity, we reported confidence levels using diamond (◊, 0.01 < *p* ≤ 0.05), stars (*, 0.001 < *p* ≤ 0.01; **, 1x10^-4^ < *p* ≤ 0.001; ***, *p* ≤ 1x10^-4^) or ns (non-significant, *p* > 0.05). We abbreviated the standard deviation by SD. Last, we used the Kolmogorov–Smirnov test to reveal whether the two distributions differed significantly (distributions of contact durations or of local curvatures).

## Supporting information

**Figure S1: mNG::ZYG-8 colocalises with mCherry::tubulin on both astral and spindle microtubules.**

(**A-F**) Exemplar fixed untreated embryo in metaphase with fluorescent labelling of (A) microtubules (mCherry::tubulin) and (B) mNG::ZYG-8, imaged using deconvolved confocal microscopy (Methods M3 & M4). (C) Merged view of panels A (magenta) and B (cyan), revealing co-localisation in white. (D) Zoom of the region outlined in panel C, with a yellow line indicating the location of the intensity profile shown in panel E. (E) Intensity profile of mCherry::tubulin and mNG::ZYG-8 fluorescence signals along the line shown in panel D. (F) Scatter plot of fluorescence intensities from the two channels shown in panels A & B. The Pearson correlation coefficient indicates significant co-localisation, calculated (black) either over the entire frame or (orange) after masking the spindle area (Method M5). (**G-L**) Exemplar fixed embryo overexpressing *zyg-8* in metaphase with fluorescent labelling of (G) microtubules (mCherry::tubulin) and (H) mNG::ZYG-8, imaged using deconvolved confocal microscopy. (I) Merged view of panels G (magenta) and H (cyan), revealing co-localisation in white. (J) Zoom of the region outlined in panel I, with a yellow line indicating the location of the intensity profile shown in panel K. (K) Intensity profile of mCherry::tubulin and mNG::ZYG-8 fluorescence signals along the line shown in panel J. (L) Scatter plot of the fluorescence intensities from the two channels shown in panels G & H. The Pearson correlation coefficient indicates significant co-localisation, calculated (black) either over the entire frame or (orange) after masking the spindle area (Method M5). Scale bars represent 10 μm, unless otherwise stated.

**Figure S2: ZYG-8 localisation to astral and spindle microtubules correlates with its expression level.**

**(A-B)** Western blots of *C. elegans* lysates showing OLLAS-tagged ZYG-8 and tubulin levels: (A) overexpression strain MCP554 (*pPie-1::mNG::3*OLLAS::zyg-8*) compared to strain MCP347; (B) RNAi-mediated depletion of *zyg-8* in the endogenous 3×OLLAS::*zyg-8* strain MCP437 compared to control RNAi (empty vector L4440). Wild-type N2 worms were used as a negative control (Method M6). (**C-F**) Fixed embryos in metaphase labelled with mCherry::tubulin (microtubules) and mNG::ZYG-8, imaged by deconvolved confocal microscopy (Methods M3 & M4), used to quantify the fluorescence ratio of mNG::ZYG-8 to mCherry::tubulin: (C) exemplar untreated embryo, (D) exemplar embryo overexpressing *zyg-8*, (E) exemplar control RNAi-treated embryo, and (F) exemplar *zyg-8(RNAi)*-treated embryo. Scale bars represent 10 μm. (**G-H**) Quantification of mNG::ZYG-8 over mCherry::tubulin fluorescence ratio: (G) in (purple) *N* = 9 *zyg-8* overexpressing embryos and (black) *N* = 9 untreated embryos; (H) in (red) *N* = 4 *zyg-8(RNAi)*-treated embryos and (black) *N* = 4 control RNAi embryos. Fluorescence measurements were performed in fixed embryos at metaphase or anaphase (Methods M3, M4 and M6) in the following regions: on spindle microtubules (2 measurements per embryo, shown as red squares in panel C), on astral microtubules (5 measurements per embryo, shown as purple lines in panel C and indicated by a purple arrow head), in the cytoplasm (3 measurements per embryo, shown as green squares in panel C), and outside the embryo (1 measurement per embryo, shown as orange square in panel C). Numbers indicate (G) the mean levels of *zyg-8* overexpression and (H) the mean levels of ZYG-8 depletion across different regions of the zygote. Brown squares and associated error bars correspond to the means and SD. *, **, and *** indicate significant differences with 1x10-3 < *p* ≤ 1x10-2, 1x10-4 < *p* ≤ 1x10-3, and *p* ≤ 1x10-4, respectively. ns denotes non-significant differences (*p* > 0.05) (Method M15).

**Figure S3: Spindle pole oscillations are exaggerated in *zyg-8(or484ts)* mutants.**

(**A-B**) Representative images used for centrosome analysis: (A) untreated embryo, and (B) *zyg-8(or484ts)* mutant, both at the restrictive temperature (Rest. T°) and expressing EBP-2::mKate2: (left) Still at anaphase onset; (middle & right) Stills captured near the time of maximal oscillation. Scale bars represent 10 μm. (**C-D**) Exemplar posterior-centrosome positions along the transverse axis for (C) the same untreated embryo as in A, and (D) the same mutant embryo as in B. Maximal amplitude is highlighted in orange, while its frequency is annotated in grey. We applied a moving-average filter over a window size of 5 s to smooth the centrosome positions.

**Figure S4: Mild depletion of the polymerase ZYG-9XMAP215 does not affect spindle-pole oscillation, spindle final position, or orientation.**

(**A**) Embryo-averaged length of the mitotic spindle along mitosis in percentage of the antero-posterior axis, measured as the distance between the two centrosomes of embryos with a GFP::TBB-2 fluorescent labelling, in (orange) *N* = 16 *zyg-9(RNAi)*-treated embryos and (black) *N* = 8 control RNAi embryos. (**B**) Growth rates of astral microtubules measured at (cross) anterior and (circle) posterior centrosomes: (orange) *Nc* = 25 comets from microtubules emanating from (cross) the anterior centrosome and (circle) *Nc* = 25 comets from posterior centrosome in 10 *zyg-9(RNAi)*-treated embryos and (black) *Nc* = 25 comets from (cross) anterior centrosome and (circle) *Nc* = 25 comets from posterior centrosome in 10 control RNAi embryos. Microtubule plus-ends were fluorescently labelled with EBP-2::mKate2 (Method M8, Figure 2A). *Nc* represents the total number of comets analysed across all replicates, separately for anterior and posterior centrosomes, with about 5 comets analysed per embryo. (**C-D**) (C) Maximal oscillation amplitudes and (D) their frequencies for the (cross) anterior and (circle) posterior centrosomes during anaphase. (**E-F**) Spindle final (E) positions along the AP axis and (F) angles. We tracked the centrosomes of the same embryos as for panel A and analysed their oscillations and positions (Method M7). *N* represents the total number of embryos analysed across all replicates. The brown squares and error bars correspond to the means and SD. *, **, and *** indicate significant differences with 1x10-3 < *p* ≤ 1x10-2, 1x10-4 < *p* ≤ 1x10-3, and *p* ≤ 1x10-4, respectively. ns denotes non-significant differences (*p* > 0.05) (Method M15).

**Figure S5: A decrease in microtubule nucleation, through partial depletion of SPD-2CEP192, does not affect spindle-pole oscillations, spindle final position, or orientation.**

(**A**) Regions centred on the centrosomes (40 x 40 pixels; 6.4 x 6.4 μm) from exemplar microscopy images showing centrosomes fluorescently labelled with GFP::TBG-1: representative images of control RNAi and *spd-2(RNAi)*-treated embryos. Scale bars represent 2 μm. **(B**) Diameters during metaphase of (cross) anterior and (circle) posterior centrosomes (Method M9): (dark blue) *N* = 11 *spd-2(RNAi)*-treated embryos and (black) *N* = 10 control RNAi embryos. (**C**) Maximal oscillation amplitudes and (**D**) their frequencies for the (cross) anterior and (circle) posterior centrosomes during anaphase. Spindle final (**E**) positions along the AP axis and (**F**) angles. Embryos are the same as for panel B. We tracked the centrosomes and analysed their positions (Method M7). *N* represents the total number of embryos analysed across all replicates. The brown squares and error bars correspond to the means and SD. ◊, *, and *** indicate significant differences with 1x10-2 < *p* ≤ 5x10-2, 1x10-3 < *p* ≤ 1x10-2, and *p* ≤ 1x10-4, respectively. ns denotes non-significant differences (*p* > 0.05) (Method M15).

**Figure S6: Spindle pole oscillations and cortical microtubule dynamics in *cls-2CLASP(RNAi)*-treated embryos do not indicate a major role for ZYG-8 in preventing microtubule catastrophe.**

(**A**) Embryo-averaged length of the mitotic spindle along mitosis in percentage of the antero-posterior axis, measured as the distance between the two centrosomes of embryos with a YFP::TBA-2 fluorescent labelling, in (green) *N* = 9 *cls-2(RNAi)*-treated embryos and (black) *N* = 9 control RNAi embryos. We tracked the centrosomes and analysed their positions (Method M7). The error bars correspond to the SD. (**B-E**) (B) Maximal oscillation amplitudes and (C) their frequencies for the (cross) anterior and (circle) posterior centrosomes during anaphase. Spindle final (D) positions along the AP axis and (E) angles. Embryos are the same as for panel A. The brown squares and error bars correspond to the means and SD. ◊ indicates significant differences with 1x10-2 < *p* ≤ 5x10-2. ns denotes non-significant differences (*p* > 0.05) (Method M15). (**F-G**) (F) Embryo-averaged density of astral microtubule contacts at the cortex plane along mitosis, and (G) embryo-averaged distributions of cortical lifetimes of astral microtubules during anaphase (Kolmogorov–Smirnov test: *p* = 0.02, Methods M10 and M15). We tracked and analysed the cortical contacts of microtubules labelled by YFP::TBA-2 in (green) *N* = 11 *cls-2(RNAi)*-treated embryos and (black) *N* = 16 control RNAi ones. *N* represents the total number of embryos analysed across all replicates.

**Figure S7: The phenotypes of spindle-pole oscillations and cortical microtubule dynamics upon *klp-7MCAK(RNAi)* treatment disfavour a role for ZYG-8 in preventing microtubule depolymerisation.**

(**A**) Embryo-averaged length of the mitotic spindle along mitosis in percentage of the antero-posterior axis, measured as the distance between the two centrosomes of embryos with a YFP::TBA-2 fluorescent labelling, in (pink) *N* = 9 *klp-7(RNAi)*-treated embryos and (black) *N* = 12 control RNAi embryos. We tracked the centrosomes and analysed their positions (Method M7). The error bars correspond to the SD. Inset: Maximal spindle elongation rate measured after nuclear envelope breakdown. (**B-E**) (B) Maximal oscillation amplitudes and (C) their frequencies for the (cross) anterior and (circle) posterior centrosomes during anaphase. Spindle final (D) positions along the AP axis and (E) angles. Embryos are the same as for panel A. The brown squares and error bars correspond to the means and SD. ** and *** indicate significant differences with 1x10-4 < *p* ≤ 1x10-3 and *p* ≤ 1x10-4, respectively. ns denotes non-significant differences (*p* > 0.05) (Method M15). (**F-G**) (F) Embryo-averaged density of astral microtubule contacts at the cortex plane along mitosis, and (G) embryo-averaged distributions of cortical lifetimes of astral microtubules during metaphase (Kolmogorov–Smirnov test: *p* = 0.42, Methods M10 and M15). We tracked and analysed the cortical contacts of microtubules labelled by YFP::TBA-2 in (green) *N* = 7 *klp-7(RNAi)*-treated embryos and (black) *N* = 12 control RNAi ones. *N* represents the total number of embryos analysed across all replicates.

**Figure S8: Quantifying microtubule curvature and tortuosity in fixed embryos.**

(**A-C**) Three complementary approaches used to quantify microtubule shapes are illustrated using schematic diagrams: (A) comparison of the local curvature distributions (at the pixel level) across all microtubules between two images; (B) for each microtubule, the 95th percentile of its local curvature distribution was used as a robust proxy of maximal curvature; and (C) tortuosity was computed for each microtubule as the ratio of curvilinear distance to end-to-end distance (Method M11). (**D**) Flow diagram of the image-analysis pipeline assembled to quantify the local curvatures of the astral microtubules (Method M11). (**E-J**) Exemplar analysis of an untreated fixed embryo with α-tubulin immunostaining: (E) the extended depth of field over three successive z-planes (F) was processed using the Oriented Filter Transform (OFT). (G) Segmentation was performed using the semi-supervised method *Ilastik* to generate the binary image. (H) This image was then skeletonised, focusing on astral filaments longer than 4 μm. (I, J) Local curvatures were calculated along each filament and (I) displayed into a colour-encoded curvature map. (J) We also generated the local curvature distribution from all microtubule pixels. Scale bars represent 10 μm.

**Figure S9: In *zyg-8(RNAi)*-treated embryos, astral microtubules are more bent than in control RNAi embryos during anaphase spindle-pole oscillations.**

Time-lapse confocal super-resolved images along anaphase of (**A**) an exemplar *zyg-8(RNAi)*-treated embryo and (**B**) an exemplar control RNAi embryo. Microtubules were labelled by GFP::TBB-2. Time is indicated from first image in mm:ss:ms. Scale bars represent 10 μm. These time-lapse images are sourced from the movies S8 and S9. Red arrows highlight bent microtubules. Similar images for *zyg-8(or484ts)* mutant and *zyg-8* overexpression are shown in Figures 5 and S10, with associated movies provided as Movies S5-S7 and S10-S11.

**Figure S10: In embryos overexpressing *zyg-8*, astral microtubules are similarly bent as in untreated embryos during anaphase spindle-pole oscillations.**

Time-lapse confocal super-resolved images along anaphase of (**A**) an exemplar embryo overexpressing *zyg-8* and (**B**) an exemplar untreated embryo. Microtubules were labelled by mCherry::tubulin. Time is indicated from first image in mm:ss:ms. Scale bars represent 10 μm. These time-lapse images are sourced from the movies S10 and S11. Red arrows highlight bent microtubules. Similar images for *zyg-8(or484ts)* mutant and *zyg-8(RNAi)* are shown in Figures 5 and S9, with associated movies provided as Movies S5-S9

**Figure S11: Image processing pipeline mapping the curvatures of astral microtubules.**

The deconvolved confocal images of fixed embryos with α-tubulin immunostaining reveal microtubule shapes during anaphase: (left) the microscopy image, and (right) its curvature map for the four conditions investigated in Figure 6. (**A**) control RNAi embryo, (**B**) *zyg-8(RNAi)*-treated embryo, (**C**) untreated embryo at the restrictive temperature, and (**D**) *zyg-8(or484ts)* mutant at the restrictive temperature (Methods M3 and M4). Scale bars represent 10 μm. Red arrows indicate fragmented microtubules in the *zyg-8(or484ts)* mutant. For each condition are indicated the number of microtubules analysed (*N*MT) and the total count of pixels (*N*pixels).

**Figure S12: *In silico,* decreasing microtubule flexural rigidity increases cortical lifetimes.**

(**A**) Snapshots from *Cytosim* agent-centred simulations showing fibres mimicking microtubules in white, with cortical contacts indicated by red dots (Table S2, Method M12). Simulations were performed using three different microtubule flexural rigidity values: *K* = 2, 10 and 25 pN.μm2. The frames were taken 45 seconds after the start of each simulation, corresponding to movies S12, S13 and S14. (**B-C**) Parameters of astral microtubules extracted from simulations varying microtubule rigidity: (B) medians of the 95th percentile curvatures and (C2) medians of tortuosity values were calculated per simulation. These measurements were taken from the set of microtubules present in the frame recorded 45 seconds after simulation start. (**D**) Cortical lifetimes of astral microtubules were calculated per simulation, using data collected from 15 seconds onward to allow for system warm-up. Black squares represent the mean, and error bars indicate the standard deviation across 40 simulations per condition.

**Figure S13: Reducing cortical pulling forces rescues mitotic spindle misorientation at anaphase end.**

(**A**) Most-posterior position of the mitotic spindle during anaphase and (**B**) mitotic spindle angle at 150 s after anaphase onset for the following conditions, all at the restrictive temperature (Rest. T°): (purple) *N* = 8 *gpr-1/2(RNAi)*-treated non-mutant embryos, (black) *N*= 8 control RNAi non-mutant embryos, (pink) *N* = 21 *zyg-8(or484ts)* mutants treated with *gpr-1/2(RNAi)* and (grey) *N* = 12 *zyg-8(or484ts)* mutants treated with control RNAi. We tracked and analysed the centrosomes of GFP::TBB-2 labelled embryos (Method M7). *N* represents the total number of embryos analysed across all replicates. The brown squares and error bars correspond to the means and SD. * and *** indicate significant differences with 1x10-3 < *p* ≤ 1x10-2 and *p* ≤ 1x10-4, respectively. ns denotes non-significant differences (*p* > 0.05) (Method M15).

**Figure S14: PTL-1 or SPD-1 does not play any major role in spindle-pole oscillations, spindle final position or orientation.** (**A**) Maximal amplitudes of spindle pole oscillations and (**B**) frequencies of the oscillations for (green) *N* = 13 *ptl-1(RNAi)*-treated embryos and (black) *N* = 18 control RNAi embryos. We tracked and analysed the centrosomes labelled by mCherry::TBG-1 (Method M7). Spindle final (**C**) positions along the AP axis and (**D**) angles. Embryos are the same as for panels A-B. (**E**) Maximal amplitudes of spindle pole oscillations and (**F**) frequencies of the oscillations for (green) *N* = 11 *spd-1(RNAi)*-treated embryos and (black) *N* = 10 control RNAi embryos. We tracked and analysed the centrosomes labelled by GFP::TBG-1 (Method M7). Spindle final (**G**) positions along the AP axis and (**H**) angles. Embryos are the same as for panels E-F. *N* represents the total number of embryos analysed across all replicates. The brown squares and error bars correspond to the means and SD. ◊ and * indicate significant differences with 1x10-2 < *p* ≤ 5x10-2 and 1x10-3 < *p* ≤ 1x10-2, respectively. ns denotes non-significant differences (*p* > 0.05) (Method M15).

**Movie S1**: Exemplar movie of a control RNAi embryo with fluorescently labelled microtubules (GFP::TBB-2), imaged at the cortex plane using a spinning disk confocal microscope (Methods M3 and M4). Tracked astral microtubule contacts are highlighted as green dots (Method M10). The image shown in Figure 4A was taken from this movie. The movie is 3-times accelerated.

**Movie S2**: Exemplar movie of a *zyg-8(RNAi)*-treated embryo with fluorescently labelled microtubules (GFP::TBB-2), imaged at the cortex plane using a spinning disk confocal microscope (Methods M3 and M4). Tracked astral microtubule contacts are highlighted as green dots (Method M10). The image shown in Figure 4B was taken from this movie. The movie is 3-times accelerated.

**Movie S3:** Exemplar movie of an untreated embryo with fluorescently labelled microtubules (mCherry::tubulin), imaged at the cortex plane using a spinning disk confocal microscope (Methods M3 and M4). Tracked astral microtubule contacts are highlighted as green dots (Method M10). The movie is 3-times accelerated.

**Movie S4:** Exemplar movie of a *zyg-8* overexpressing embryo with fluorescently labelled microtubules (mCherry::tubulin), imaged at the cortex plane using a spinning disk confocal microscope (Methods M3 and M4). Tracked astral microtubule contacts are highlighted as green dots (Method M10). The movie is 3-times accelerated.

**Movie S5**: First exemplar live *zyg-8(or484ts)* mutant at the restrictive temperature with fluorescently labelled microtubules (GFP::TBB-2) imaged using a super-resolved confocal microscope at the spindle plane (Methods M3 and M4). The images shown in Figure 5A were taken from this movie. The movie is 10-times accelerated.

**Movie S6**: Second exemplar live *zyg-8(or484ts)* mutant at the restrictive temperature with fluorescently labelled microtubules (GFP::TBB-2) imaged using a super-resolved confocal microscope at the spindle plane (Methods M3 and M4). The movie is 10-times accelerated.

**Movie S7**: Exemplar live untreated embryo at the restrictive temperature with fluorescently labelled microtubules (GFP::TBB-2) imaged using a super-resolved confocal microscope at the spindle plane (Methods M3 and M4). The images shown in Figure 5B were taken from this movie. The movie is 10-times accelerated.

**Movie S8**: Exemplar live *zyg-8(RNAi)*-treated embryo with fluorescently labelled microtubules (GFP::TBB-2) imaged using a super-resolved confocal microscope at the spindle plane (Methods M3 and M4). The images shown in Figure S9A were taken from this movie. The movie is 10-times accelerated.

**Movie S9**: Exemplar live control RNAi embryo with fluorescently labelled microtubules (GFP::TBB-2) imaged using a super-resolved confocal microscope at the spindle plane (Methods M3 and M4). The images shown in Figure S9B were taken from this movie. The movie is 10-times accelerated.

**Movie S10**: Exemplar live *zyg-8* overexpressing embryo with fluorescently labelled microtubules (mCherry::tubulin) imaged using a super-resolved confocal microscope at the spindle plane (Methods M3 and M4). The images shown in Figure S10A were taken from this movie. The movie is 10-times accelerated.

**Movie S11**: Exemplar live untreated embryo with fluorescently labelled microtubules (mCherry::tubulin) imaged using a super-resolved confocal microscope at the spindle plane (Methods M3 and M4). The images shown in Figure S10B were taken from this movie. The movie is 10-times accelerated.

**Movie S12**: Exemplar simulation with the rigidity of astral microtubule set to 2 pN·µm² (Method M12). All the other simulation parameters are identical to those in Movies S9 and S10. The red and blue disks are the anterior and posterior centrosomes, respectively. Microtubules are shown as white filaments, and cortical contacts as red dots. Note that the first 15 seconds are the warm-up phase of the simulation.

**Movie S13**: Exemplar simulation with the rigidity of astral microtubule set to 10 pN·µm² (Method M12). All the other simulation parameters are identical to those in Movies S8 and S10. The red and blue disks are the anterior and posterior centrosomes, respectively. Microtubules are shown as white filaments, and cortical contacts as red dots. Note that the first 15 seconds are the warm-up phase of the simulation.

**Movie S14**: Exemplar simulation with the rigidity of astral microtubule set to 25 pN·µm² (Method M12). All the other simulation parameters are identical to those in Movies S8 and S9. The red and blue disks are the anterior and posterior centrosomes, respectively. Microtubules are shown as white filaments, and cortical contacts as red dots. Note that the first 15 seconds are the warm-up phase of the simulation.

**Movie S15**: Exemplar movie of a control RNAi embryo with fluorescently labelled dynein (GFP::DHC-1), imaged at the cortex plane using a spinning disk confocal microscope (Methods M3 and M4)

**Table S1:** List of the strains used in this study.

(* 4-times backcrossed; ** 2-times backcrossed.)

**Table S2:** Objects and their parameters used in the three *Cytosim* simulations (Method M12).

**Table S3: Changes in cortical dynein density and lifetime upon ZYG-8 depletion phenocopy variations in cortical astral microtubule dynamics.**

Characteristics of the two cortical dynein populations in strains expressing GFP::DHC-1. Data were obtained by fitting the duration histograms of cortical dynein spots using a two-exponential model (Method M13). *N* represents the total number of embryos analysed across all replicates for each experimental condition.

**File S1**: *Cytosim* file with the rigidity of astral microtubule set to 2 pN·µm² (Method M12).

**File S2**: *Cytosim* file with the rigidity of astral microtubule set to 10 pN·µm² (Method M12).

**File S3**: *Cytosim* file with the rigidity of astral microtubule set to 25 pN·µm² (Method M12).

## Acknowledgements

Strains TH27 and TH30 were kindly offered by Prof Anthony A. Hyman. We thank Dr Grégoire Michaux for supplying the feeding clone library and for technical support. Strains MCP347, MCP361 & MCP554 were generated by SEGiCel (SFR Santé Lyon Est CNRS UAR 3453, Lyon, France) with the support of CNRS and IBiSA. We thank S. Dutertre and X. Pinson of the Microscopy Rennes Imaging Center (MRic, BIOSIT, Biogenouest) for their assistance. MRic is part of the national infrastructure France-BioImaging supported by the French National Research Agency (ANR-10-INBS-04). We thank the Pécréaux lab for their support, and the Gibeaux, Giet, Huet, Michaux, and Tramier labs for helpful discussions. We also thank L. Chesneau for assistance in designing the CRISPR strains, and A. Pacquelet, B. Lacroix and C. Heligon for their careful reading of the manuscript.

## Author contributions

Conceptualisation: Hélène Bouvrais, Jacques Pécréaux.

Experimental data: Louis Cueff, Ewen Huet, Loïc Sschmitt, Sylvain Pastezeur, Méline Coquil, Talia Savary, Anouk Sénard, Hélène Bouvrais.

Data curation: Hélène Bouvrais.

Formal analysis: Louis Cueff, Ewen Huet, Méline Coquil, Jacques Pécréaux, Hélène Bouvrais.

Funding acquisition: Jacques Pécréaux, Hélène Bouvrais.

Methodology: Louis Cueff, Jacques Pécréaux, Hélène Bouvrais.

Project administration: Hélène Bouvrais.

Software: Jacques Pécréaux, Hélène Bouvrais.

Supervision: Jacques Pécréaux, Hélène Bouvrais.

Validation: Louis Cueff, Ewen Huet, Jacques Pécréaux, Hélène Bouvrais.

Visualisation: Louis Cueff, Hélène Bouvrais.

Writing – original draft: Hélène Bouvrais.

Writing – review and editing: Louis Cueff, Jacques Pécréaux, Hélène Bouvrais.

## Data availability statement

All relevant data are included in the manuscript and its supporting information files.

