## Supplemental file 1 for "Microtubule stiffening by the doublecortin-domain protein ZYG-8 contributes to mitotic spindle orientation during zygote division in *Caenorhabditis elegans*"

```
% Cueff L. et al., 2025

% Microtubule stiffening by the doublecortin-domain protein ZYG-8
contributes to mitotic spindle orientation during zygote division in
Caenorhabditis elegans.

% Supplemental file S1 for cytosim simulation

% Fiber rigidity set to 2 pN.um2

%——————————————————————————————————————————————————
set simul celegans_curvature
{
    time_step = 0.01
    viscosity = 5
}

%—————————————————————————————————————————————————
% draw ellipses of the embryo shape

set space cell
{
    geometry = ( ellipse 24.5 16.5 16.5 )
}

new space cell

%————————————————
%———————————————————————————————————————————————————
% simulate astral MT which are dynamic, semi-flexible and engaged
with cortical FGs

% simulate anchoring of astral microtubules at the cortex

set hand strong_hand
{
    unbinding_rate = 0.1
    unbinding_force = 5
    hold_growing_end = 1
    hold_shrinking_end = 1
    display = { size=8 ; color=red}
}

set single cortical_glue
{
    hand = strong_hand
    stiffness = 25
    activity = fixed
}


%———————————————

set fiber astralMTs
```

```
{
    rigidity = 2
    segmentation = 0.5
    confine = inside, 100
    glue = 3, cortical_glue

    activity = classic
    growing_force   = 1.67
    min_length      = 0.005
    growing_speed  = 0.71
    shrinking_speed = -0.84
    catastrophe_rate = 0.05
        catastrophe_rate_stalled = 0.5
    rescue_rate = 0.15
    rebirth_rate = inf
    persistent = 1
    total_polymer = inf
    display = { line_width=1; color=white; }
}


%-----------------------------------------------------------------
%------------
% to build the mitotic spindle (2 centrosomes + spindle MTs)

% define spindle MTs as static fiber

set fiber spindleMTs
{
    rigidity = 100
    segmentation = 0.5
    confine = inside, 100
    binding_key = 2
    display = {line_width = 0.5; color=pink}
}

%-------------------------
% definition of CS anterior

set solid core1
{
    confine = inside, 100
    external_force = -180 5 1
    display = ( style=31; color=red )
}

set aster centrosome1
{
        stiffness = 100, 10
    nb_fibers_type = 3
}


new aster centrosome1
```

```
{
    solid = core1
    position = (-5.6 0 0)
    radius = 1
        orientation = 1 0 0
        fibers1 = 20, spindleMTs, (asterProp_type = angular;
asterProp_aster_angle = -0.52, 0.52; length = 6; plus_end = grow;)
    fibers2 = 38, astralMTs, (asterProp_type = angular;
asterProp_aster_angle = 1.05, 3.14 ; length = 8, 6; plus_end =
grow;)
    fibers3 = 38, astralMTs, (asterProp_type = angular;
asterProp_aster_angle = -3.14, -1.05 ; length = 8, 6; plus_end =
grow;)
}

%———————————————————————
% define posterior centrosome
set solid core2
{
    confine = inside, 100
    external_force = 300 0 0
    display = ( style=31; color=blue )
}

set aster centrosome2
{
        stiffness = 100, 10
    nb_fibers_type = 4
}


new aster centrosome2
{
    solid = core2
    position = (4.7 0 0)
    radius = 1
        orientation = 1 0 0
        fibers1 = 10, spindleMTs, (asterProp_type = angular;
asterProp_aster_angle = 2.62, 3.14; length = 6; plus_end = grow;)
        fibers2 = 10, spindleMTs, (asterProp_type = angular;
asterProp_aster_angle = -3.14, -2.62; length = 6; plus_end = grow;)
    fibers3 = 38, astralMTs, (asterProp_type = angular;
asterProp_aster_angle = -2.09, 0 ; length = 8, 6; plus_end = grow;)
    fibers4 = 38, astralMTs, (asterProp_type = angular;
asterProp_aster_angle = 0, 2.09 ; length = 8, 6; plus_end = grow;)
}


%—————————————————————————————————————————————————
run simul *
{
    write_object = fiber
    nb_steps = 6000
    nb_frames = 600
```

```
}
```
