## Supplemental material for "Microtubule stiffening by the doublecortin-domain protein ZYG-8 contributes to mitotic spindle orientation during zygote division in *Caenorhabditis elegans*"

### Supplemental figures and tables

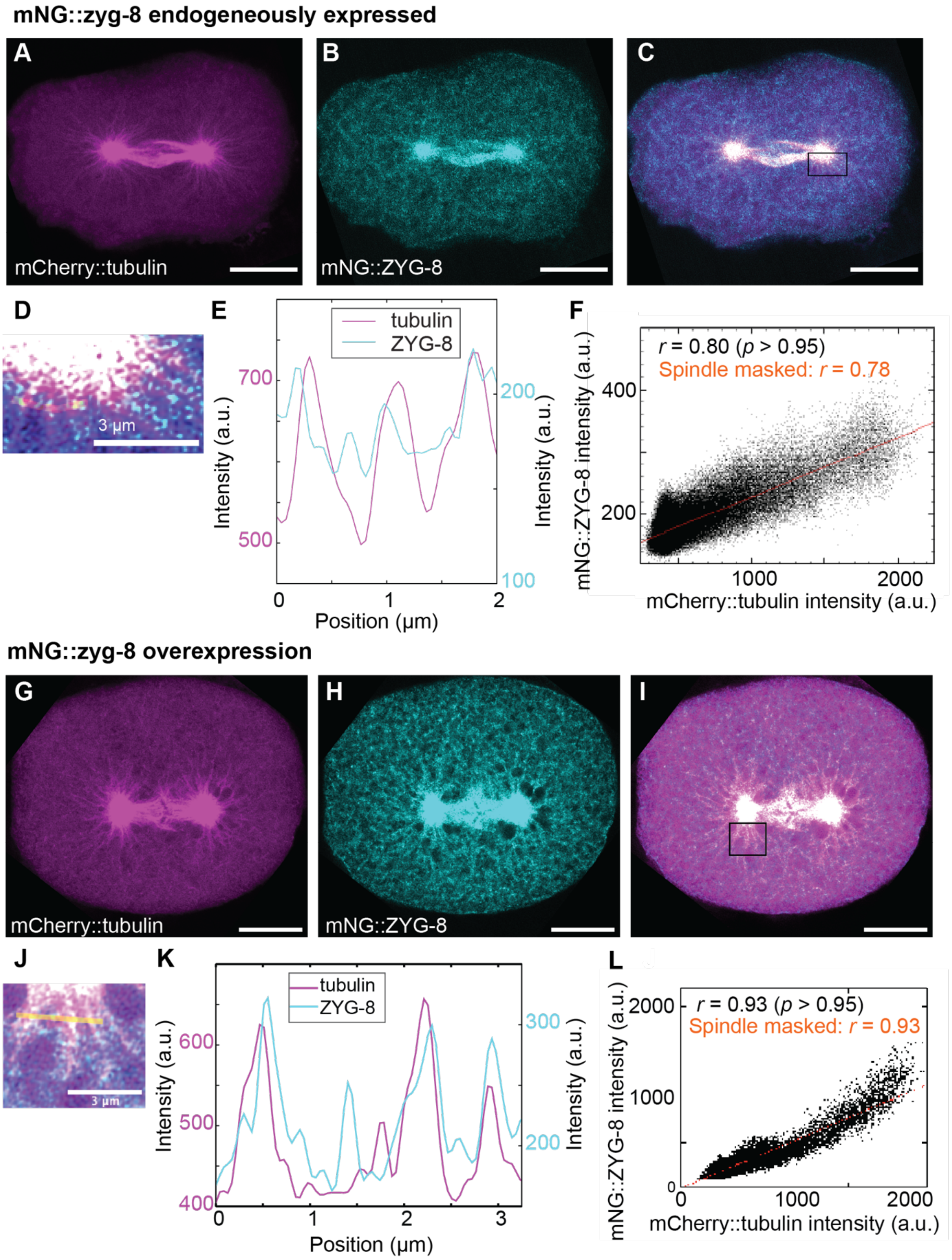


**Figure S1: mNG::ZYG-8 colocalises with mCherry::tubulin on both astral and spindle microtubules.**

Microtubule stiffening by ZYG-8 contributes to spindle orientation.

(**A-F**) Exemplar fixed untreated embryo in metaphase with fluorescent labelling of (A) microtubules (mCherry::tubulin) and (B) mNG::ZYG-8, imaged using deconvolved confocal microscopy (Methods M3 & M4). (C) Merged view of panels A (magenta) and B (cyan), revealing co-localisation in white. (D) Zoom of the region outlined in panel C, with a yellow line indicating the location of the intensity profile shown in panel E. (E) Intensity profile of mCherry::tubulin and mNG::ZYG-8 fluorescence signals along the line shown in panel D. (F) Scatter plot of fluorescence intensities from the two channels shown in panels A & B. The Pearson correlation coefficient indicates significant co-localisation, calculated (black) either over the entire frame or (orange) after masking the spindle area (Method M5). (**G-L**) Exemplar fixed embryo overexpressing *zyg-8* in metaphase with fluorescent labelling of (G) microtubules (mCherry::tubulin) and (H) mNG::ZYG-8, imaged using deconvolved confocal microscopy. (I) Merged view of panels G (magenta) and H (cyan), revealing co-localisation in white. (J) Zoom of the region outlined in panel I, with a yellow line indicating the location of the intensity profile shown in panel K. (K) Intensity profile of mCherry::tubulin and mNG::ZYG-8 fluorescence signals along the line shown in panel J. (L) Scatter plot of the fluorescence intensities from the two channels shown in panels G & H. The Pearson correlation coefficient indicates significant co-localisation, calculated (black) either over the entire frame or (orange) after masking the spindle area (Method M5). Scale bars represent 10 $\mu$m, unless otherwise stated.

Microtubule stiffening by ZYG-8 contributes to spindle orientation.

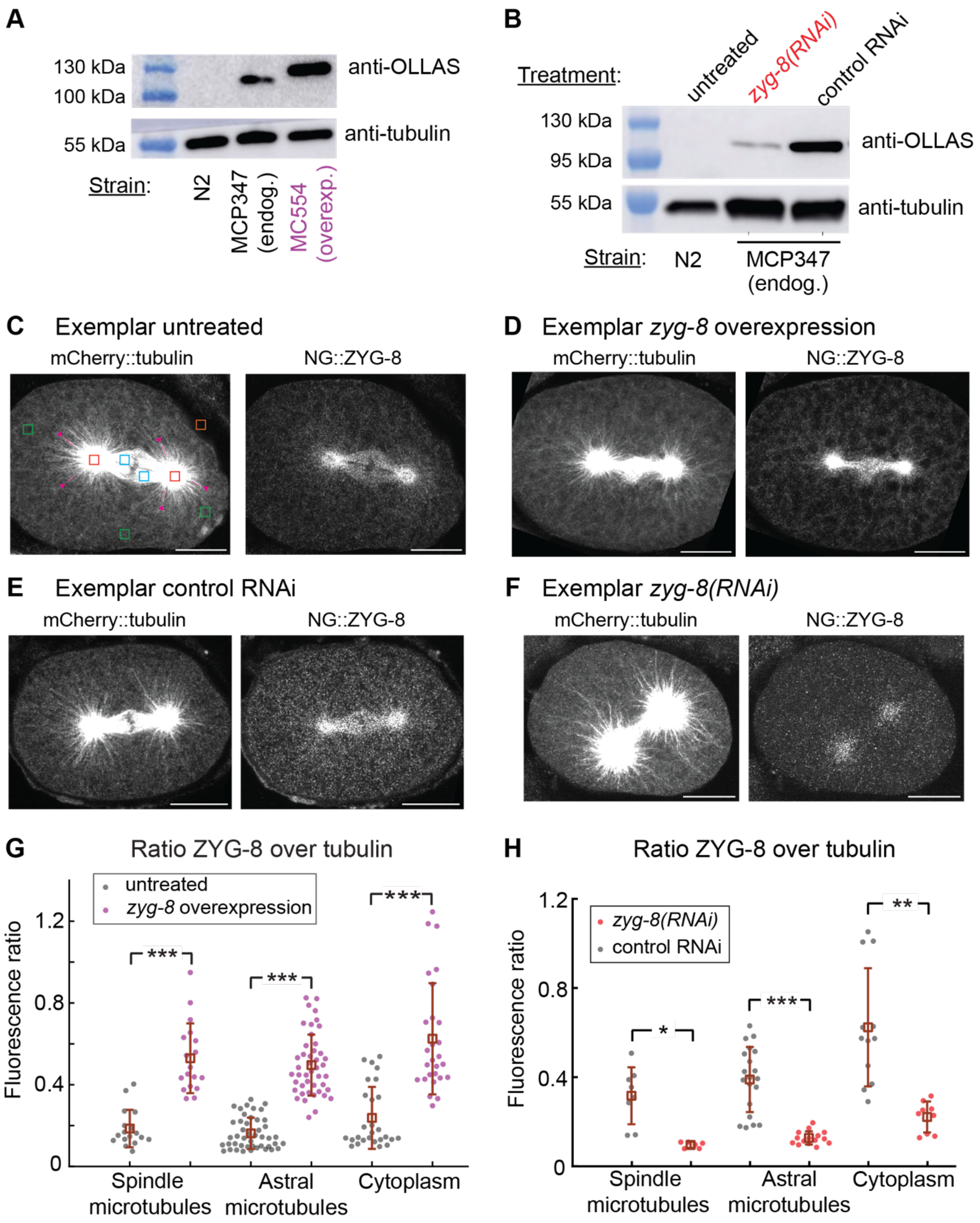


**Figure S2: ZYG-8 localisation to astral and spindle microtubules correlates with its expression level.** **(A-B)** Western blots of *C. elegans* lysates showing OLLAS-tagged ZYG-8 and tubulin levels: (A) overexpression strain MCP554 (*pPie-1::mNG::3*OLLAS::zyg-8*) compared to strain MCP347; (B) RNAi-mediated depletion of *zyg-8* in the endogenous 3×OLLAS::*zyg-8* strain MCP437 compared to control RNAi (empty vector L4440). Wild-type N2 worms were used as a negative control (Method M6). (**C-F**) Fixed embryos in metaphase labelled with mCherry::tubulin (microtubules) and mNG::ZYG-8, imaged by deconvolved confocal microscopy (Methods M3 & M4), used to quantify the fluorescence ratio of mNG::ZYG-8 to mCherry::tubulin: (C) exemplar untreated embryo, (D) exemplar embryo overexpressing *zyg-8*, (E) exemplar control RNAi-treated embryo, and (F) exemplar *zyg-8(RNAi)*-

treated embryo. Scale bars represent 10 μm. (**G-H**) Quantification of mNG::ZYG-8 over mCherry::tubulin fluorescence ratio: (G) in (purple) *N* = 9 *zyg-8* overexpressing embryos and (black) *N* = 9 untreated embryos; (H) in (red) *N* = 4 *zyg-8(RNAi)*-treated embryos and (black) *N* = 4 control RNAi embryos. Fluorescence measurements were performed in fixed embryos at metaphase or anaphase (Methods M3, M4 and M6) in the following regions: on spindle microtubules (2 measurements per embryo, shown as red squares in panel C), on astral microtubules (5 measurements per embryo, shown as purple lines in panel C and indicated by a purple arrow head), in the cytoplasm (3 measurements per embryo, shown as green squares in panel C), and outside the embryo (1 measurement per embryo, shown as orange square in panel C). Numbers indicate (G) the mean levels of *zyg-8* overexpression and (H) the mean levels of ZYG-8 depletion across different regions of the zygote. Brown squares and associated error bars correspond to the means and SD. *, **, and *** indicate significant differences with $1x10^{-3} < p \leq 1x10^{-2}$, $1x10^{-4} < p \leq 1x10^{-3}$, and $p \leq 1x10^{-4}$, respectively. ns denotes non-significant differences ($p > 0.05$) (Method M15).

Microtubule stiffening by ZYG-8 contributes to spindle orientation.

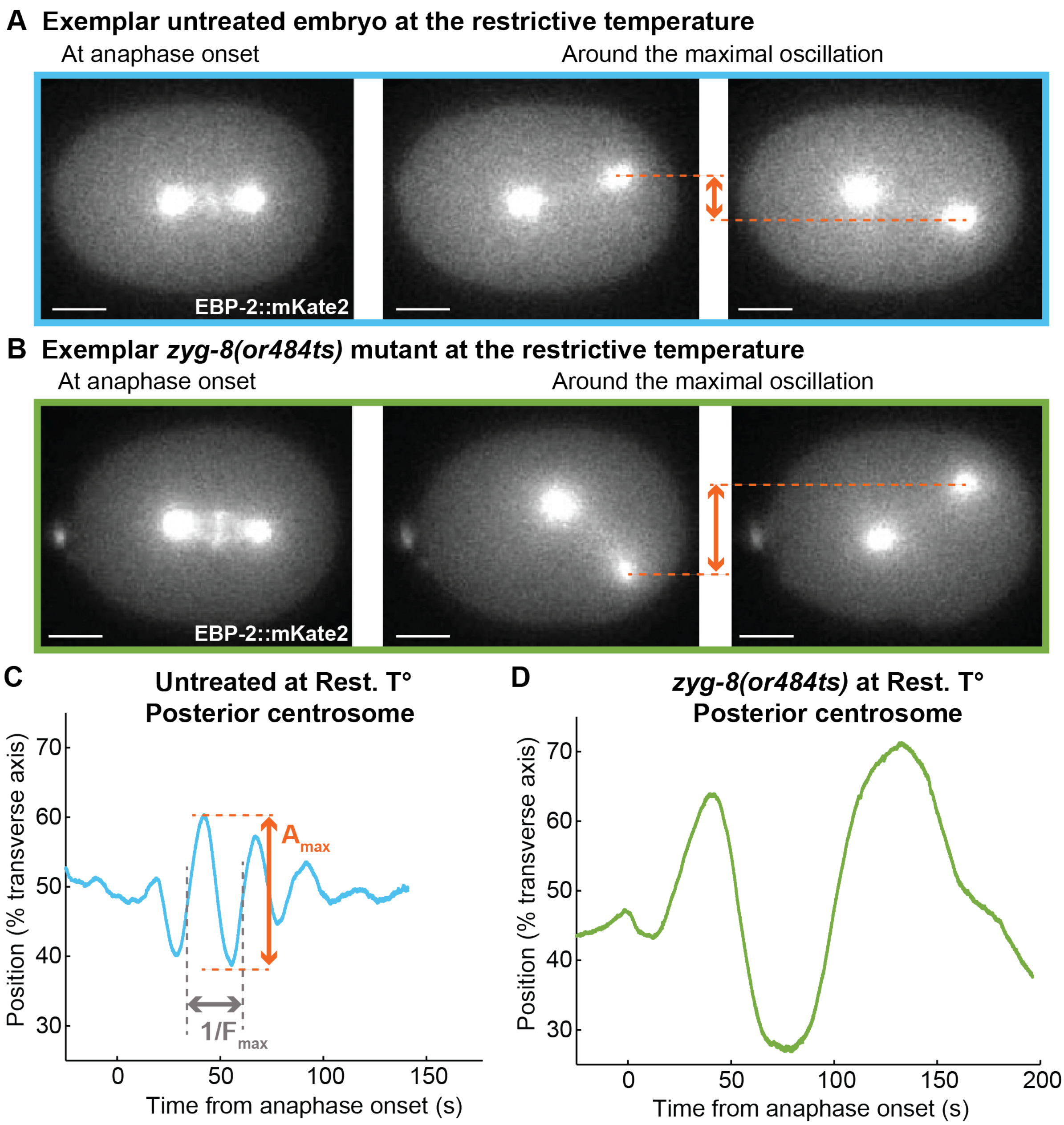


**Figure S3: Spindle pole oscillations are exaggerated in *zyg-8(or484ts)* mutants.**
(**A-B**) Representative images used for centrosome analysis: (A) untreated embryo, and (B) *zyg-8(or484ts)* mutant, both at the restrictive temperature (Rest. T°) and expressing EBP-2::mKate2: (left) Still at anaphase onset; (middle & right) Stills captured near the time of maximal oscillation. Scale bars represent 10 µm. (**C-D**) Exemplar posterior-centrosome positions along the transverse axis for (C) the same untreated embryo as in A, and (D) the same mutant embryo as in B. Maximal amplitude is highlighted in orange, while its frequency is annotated in grey. We applied a moving-average filter over a window size of 5 s to smooth the centrosome positions.

Microtubule stiffening by ZYG-8 contributes to spindle orientation.

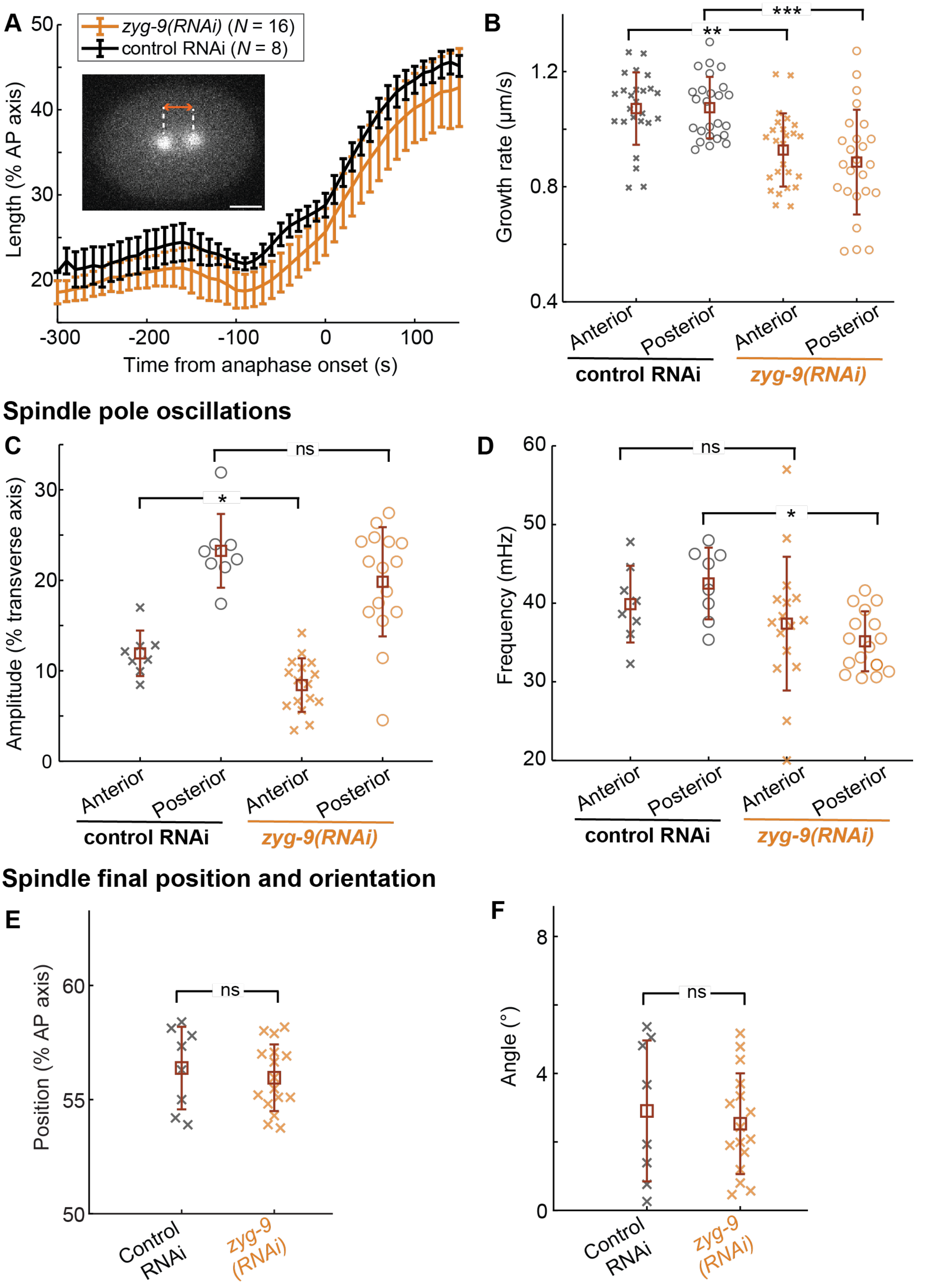


**Figure S4: Mild depletion of the polymerase ZYG-9$^{XMAP215}$ does not affect spindle-pole oscillation, spindle final position, or orientation.**

(**A**) Embryo-averaged length of the mitotic spindle along mitosis in percentage of the antero-posterior axis, measured as the distance between the two centrosomes of embryos with a GFP::TBB-2 fluorescent labelling, in (orange) $N$ = 16 *zyg-9(RNAi)*-treated embryos and (black) $N$ = 8 control RNAi embryos. (**B**) Growth rates of astral microtubules measured at (cross) anterior and (circle) posterior centrosomes: (orange) $N_c$ = 25 comets from microtubules emanating from (cross) the anterior centrosome and (circle) $N_c$ = 25 comets from posterior centrosome in 10 *zyg-9(RNAi)*-treated embryos and (black) $N_c$ = 25 comets from (cross) anterior centrosome and (circle) $N_c$ = 25 comets from posterior centrosome in 10 control RNAi embryos. Microtubule plus-ends were fluorescently labelled with EBP-2::mKate2 (Method M8, Figure 2A). $N_c$ represents the total number of comets analysed across all replicates, separately for anterior and posterior centrosomes, with about 5 comets analysed per embryo. (**C-D**) (C) Maximal oscillation amplitudes and (D) their frequencies for the (cross) anterior and (circle) posterior centrosomes during anaphase. (**E-F**) Spindle final (E) positions along the AP axis and (F) angles. We tracked the centrosomes of the same embryos as for panel A and analysed their oscillations and positions (Method M7). $N$ represents the total number of embryos analysed across all replicates. The brown squares and error bars correspond to the means and SD. *, **, and *** indicate significant differences with $1x10^{-3} < p \leq 1x10^{-2}$, $1x10^{-4} < p \leq 1x10^{-3}$, and $p \leq 1x10^{-4}$, respectively. ns denotes non-significant differences ($p > 0.05$) (Method M15).

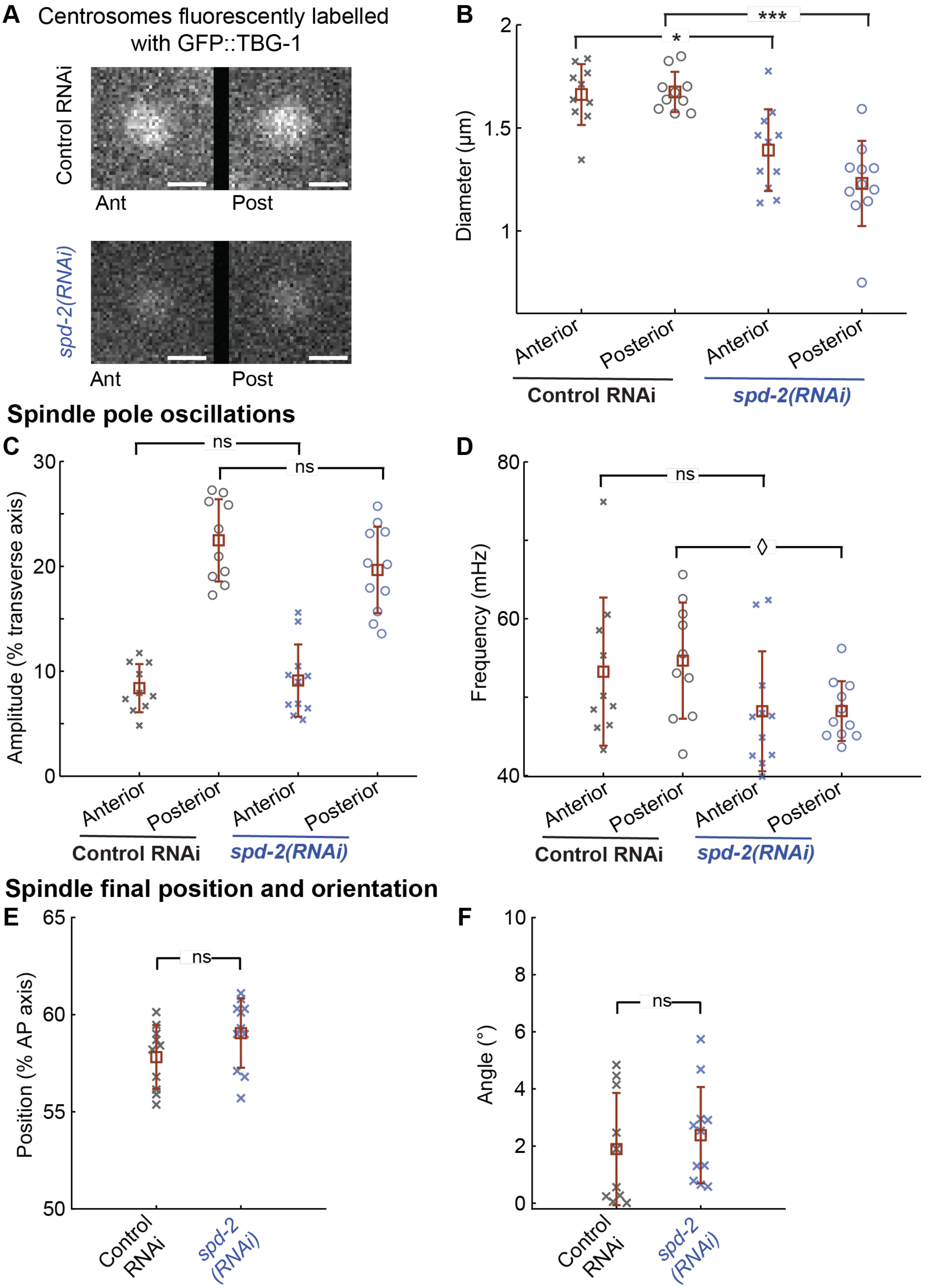


**Figure S5: A decrease in microtubule nucleation, through partial depletion of SPD-2[CEP192], does not affect spindle-pole oscillations, spindle final position, or orientation.**

Microtubule stiffening by ZYG-8 contributes to spindle orientation.

(**A**) Regions centred on the centrosomes (40 x 40 pixels; 6.4 x 6.4 μm) from exemplar microscopy images showing centrosomes fluorescently labelled with GFP::TBG-1: representative images of control RNAi and *spd-2(RNAi)*-treated embryos. Scale bars represent 2 μm. **(B**) Diameters during metaphase of (cross) anterior and (circle) posterior centrosomes (Method M9): (dark blue) *N* = 11 *spd-2(RNAi)*-treated embryos and (black) *N* = 10 control RNAi embryos. (**C**) Maximal oscillation amplitudes and (**D**) their frequencies for the (cross) anterior and (circle) posterior centrosomes during anaphase. Spindle final (**E**) positions along the AP axis and (**F**) angles. Embryos are the same as for panel B. We tracked the centrosomes and analysed their positions (Method M7). *N* represents the total number of embryos analysed across all replicates. The brown squares and error bars correspond to the means and SD. ◊, *, and *** indicate significant differences with $1x10^{-2} < p \le 5x10^{-2}$, $1x10^{-3} < p \le 1x10^{-2}$, and $p \le 1x10^{-4}$, respectively. ns denotes non-significant differences ($p > 0.05$) (Method M15).

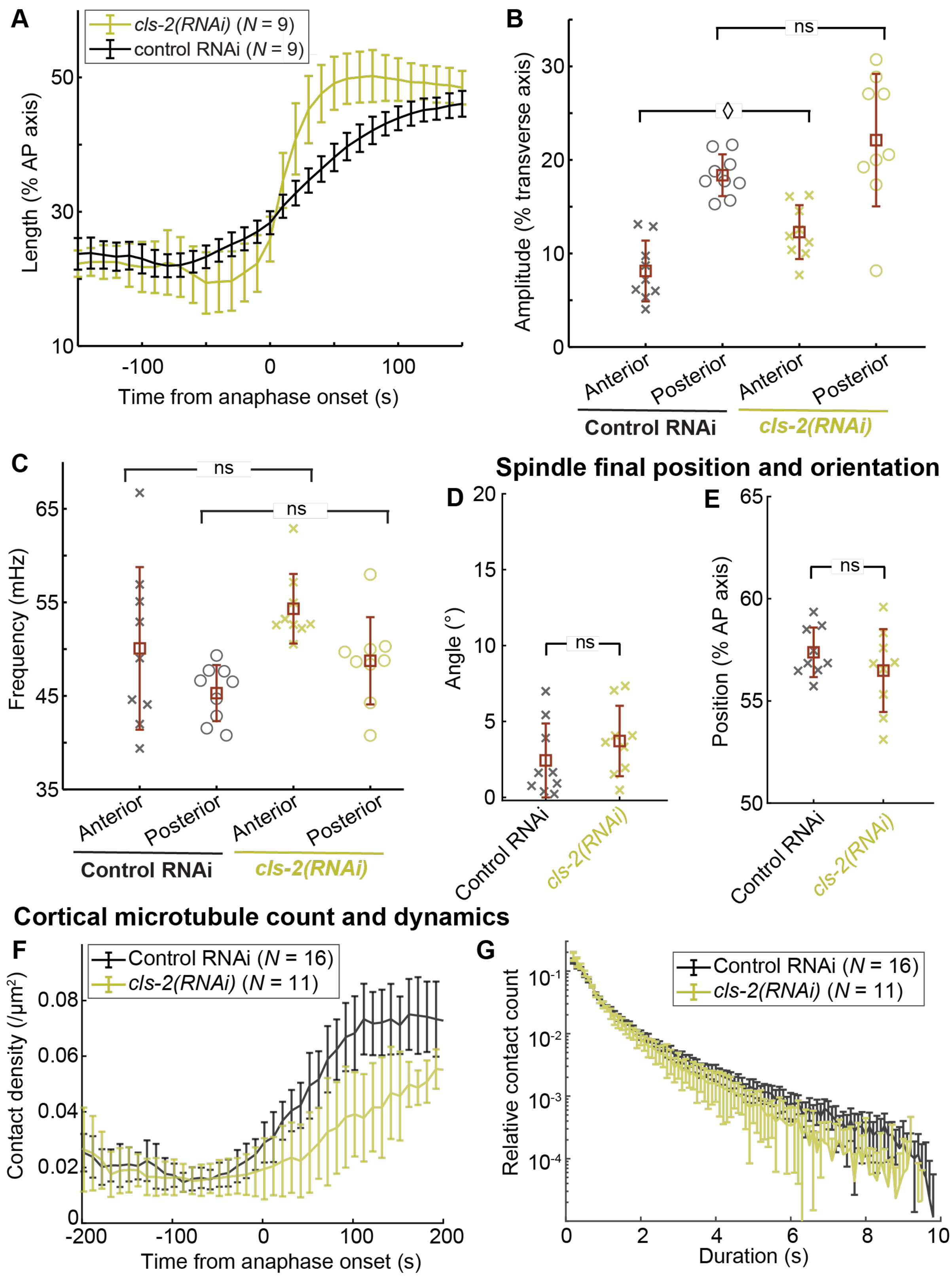


**Figure S6: Spindle pole oscillations and cortical microtubule dynamics in *cls-2*$^{CLASP}$*(RNAi)*-treated embryos do not indicate a major role for ZYG-8 in preventing microtubule catastrophe.**

(**A**) Embryo-averaged length of the mitotic spindle along mitosis in percentage of the antero-posterior axis, measured as the distance between the two centrosomes of embryos with a YFP::TBA-2 fluorescent labelling, in (green) *N* = 9 *cls-2(RNAi)*-treated embryos and (black) *N* = 9 control RNAi embryos. We tracked the centrosomes and analysed their positions (Method M7). The error bars correspond to the SD. (**B-E**) (B) Maximal oscillation amplitudes and (C) their frequencies for the (cross) anterior and (circle) posterior centrosomes during anaphase. Spindle final (D) positions along the AP axis and (E) angles. Embryos are the same as for panel A. The brown squares and error bars correspond to the means and SD. ◊ indicates significant differences with $1x10^{-2} < p \leq 5x10^{-2}$. ns denotes non-significant differences ($p > 0.05$) (Method M15). (**F-G**) (F) Embryo-averaged density of astral microtubule contacts at the cortex plane along mitosis, and (G) embryo-averaged distributions of cortical lifetimes of astral microtubules during anaphase (Kolmogorov–Smirnov test: $p = 0.02$, Methods M10 and M15). We tracked and analysed the cortical contacts of microtubules labelled by YFP::TBA-2 in (green) $N = 11$ *cls-2(RNAi)*-treated embryos and (black) $N = 16$ control RNAi ones. *N* represents the total number of embryos analysed across all replicates.

Microtubule stiffening by ZYG-8 contributes to spindle orientation.

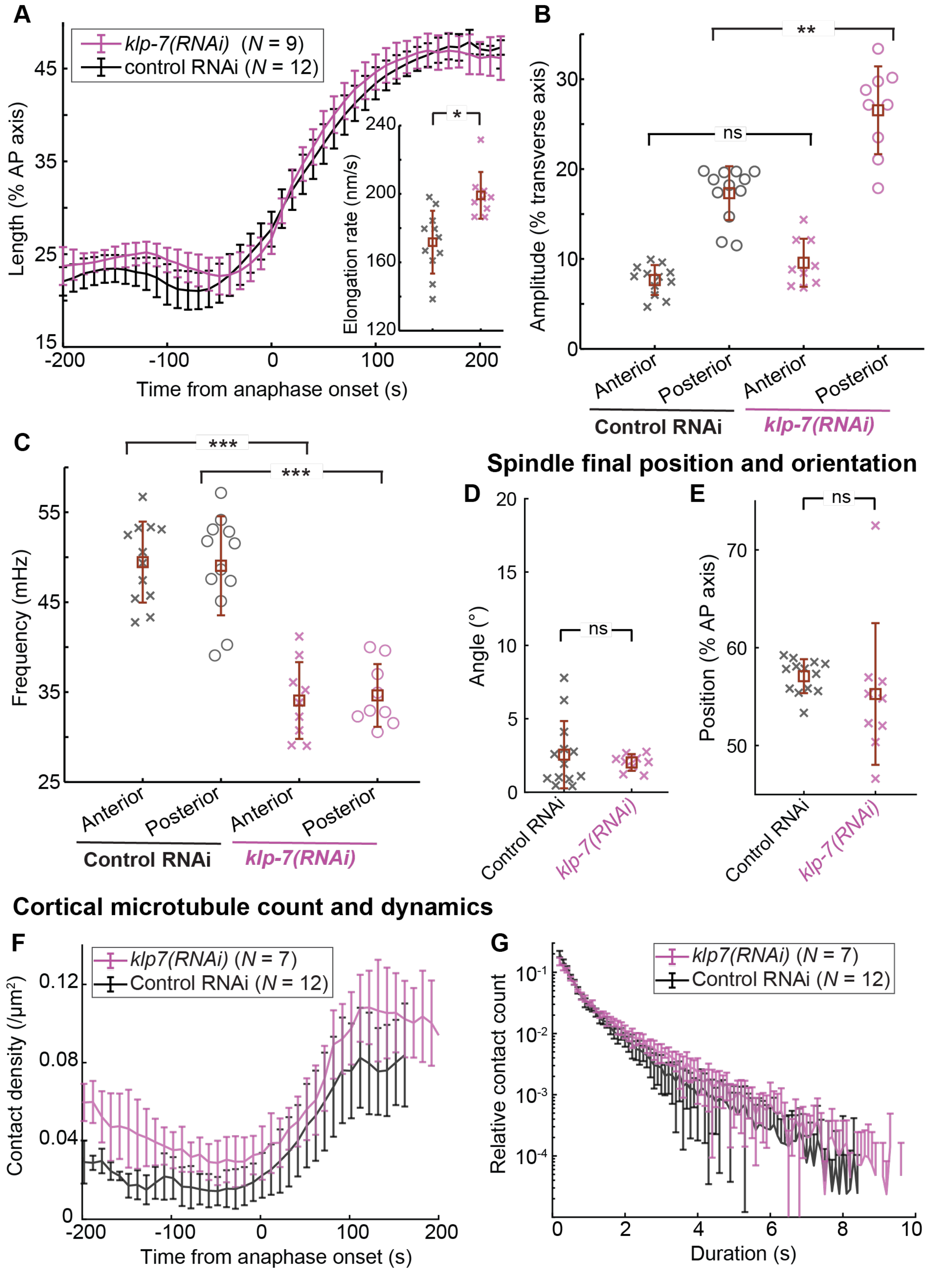


**Figure S7: The phenotypes of spindle-pole oscillations and cortical microtubule dynamics upon *klp-7*$^{MCAK}$*(RNAi)* treatment disfavour a role for ZYG-8 in preventing microtubule depolymerisation.**
(**A**) Embryo-averaged length of the mitotic spindle along mitosis in percentage of the antero-posterior axis, measured as the distance between the two centrosomes of embryos with a YFP::TBA-2

fluorescent labelling, in (pink) $N = 9$ *klp-7(RNAi)*-treated embryos and (black) $N = 12$ control RNAi embryos. We tracked the centrosomes and analysed their positions (Method M7). The error bars correspond to the SD. Inset: Maximal spindle elongation rate measured after nuclear envelope breakdown. (**B-E**) (B) Maximal oscillation amplitudes and (C) their frequencies for the (cross) anterior and (circle) posterior centrosomes during anaphase. Spindle final (D) positions along the AP axis and (E) angles. Embryos are the same as for panel A. The brown squares and error bars correspond to the means and SD. ** and *** indicate significant differences with $1x10^{-4} < p \leq 1x10^{-3}$ and $p \leq 1x10^{-4}$, respectively. ns denotes non-significant differences ($p > 0.05$) (Method M15). (**F-G**) (F) Embryo-averaged density of astral microtubule contacts at the cortex plane along mitosis, and (G) embryo-averaged distributions of cortical lifetimes of astral microtubules during metaphase (Kolmogorov–Smirnov test: $p = 0.42$, Methods M10 and M15). We tracked and analysed the cortical contacts of microtubules labelled by YFP::TBA-2 in (green) $N = 7$ *klp-7(RNAi)*-treated embryos and (black) $N = 12$ control RNAi ones. *N* represents the total number of embryos analysed across all replicates.

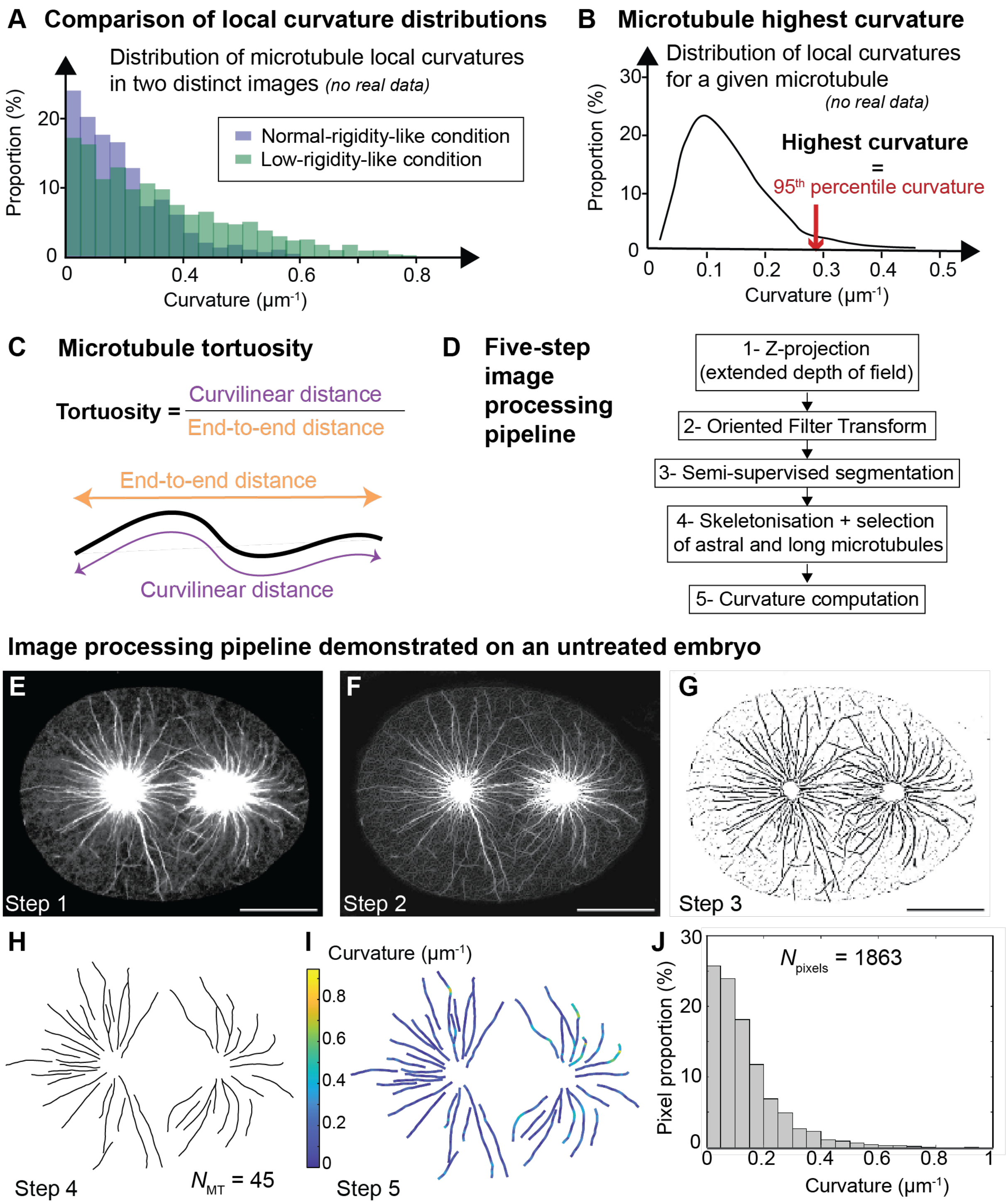


**Figure S8: Quantifying microtubule curvature and tortuosity in fixed embryos.**
(**A-C**) Three complementary approaches used to quantify microtubule shapes are illustrated using schematic diagrams: (A) comparison of the local curvature distributions (at the pixel level) across all microtubules between two images; (B) for each microtubule, the 95$^{th}$ percentile of its local curvature distribution was used as a robust proxy of maximal curvature; and (C) tortuosity was computed for each microtubule as the ratio of curvilinear distance to end-to-end distance (Method M11). (**D**) Flow diagram of the image-analysis pipeline assembled to quantify the local curvatures of the astral microtubules (Method M11). (**E-J**) Exemplar analysis of an untreated fixed embryo with $\alpha$-tubulin immunostaining: (E) the extended depth of field over three successive z-planes (F) was processed using the Oriented Filter Transform (OFT). (G) Segmentation was performed using the semi-supervised

method *Ilastik* to generate the binary image. (H) This image was then skeletonised, focusing on astral filaments longer than 4 μm. (I, J) Local curvatures were calculated along each filament and (I) displayed into a colour-encoded curvature map. (J) We also generated the local curvature distribution from all microtubule pixels. Scale bars represent 10 μm.

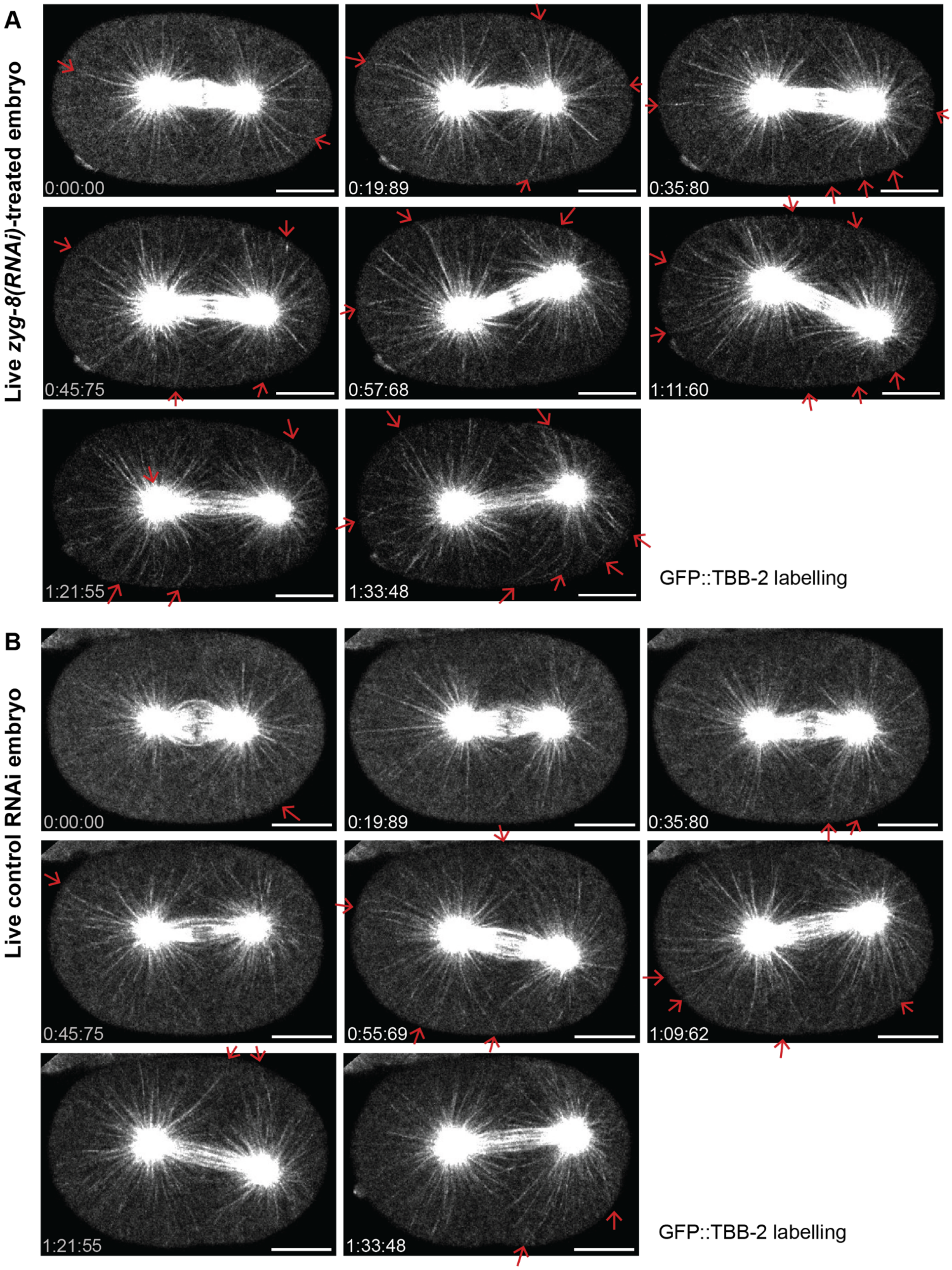


**Figure S9: In *zyg-8(RNAi)*-treated embryos, astral microtubules are more bent than in control RNAi embryos during anaphase spindle-pole oscillations.**

Time-lapse confocal super-resolved images along anaphase of (**A**) an exemplar *zyg-8(RNAi)*-treated embryo and (**B**) an exemplar control RNAi embryo. Microtubules were labelled by GFP::TBB-2. Time is indicated from first image in mm:ss:ms. Scale bars represent 10 μm. These time-lapse images are sourced from the movies S8 and S9. Red arrows highlight bent microtubules. Similar images for *zyg-8(or484ts)* mutant and *zyg-8* overexpression are shown in Figures 5 and S10, with associated movies provided as Movies S5-S7 and S10-S11.

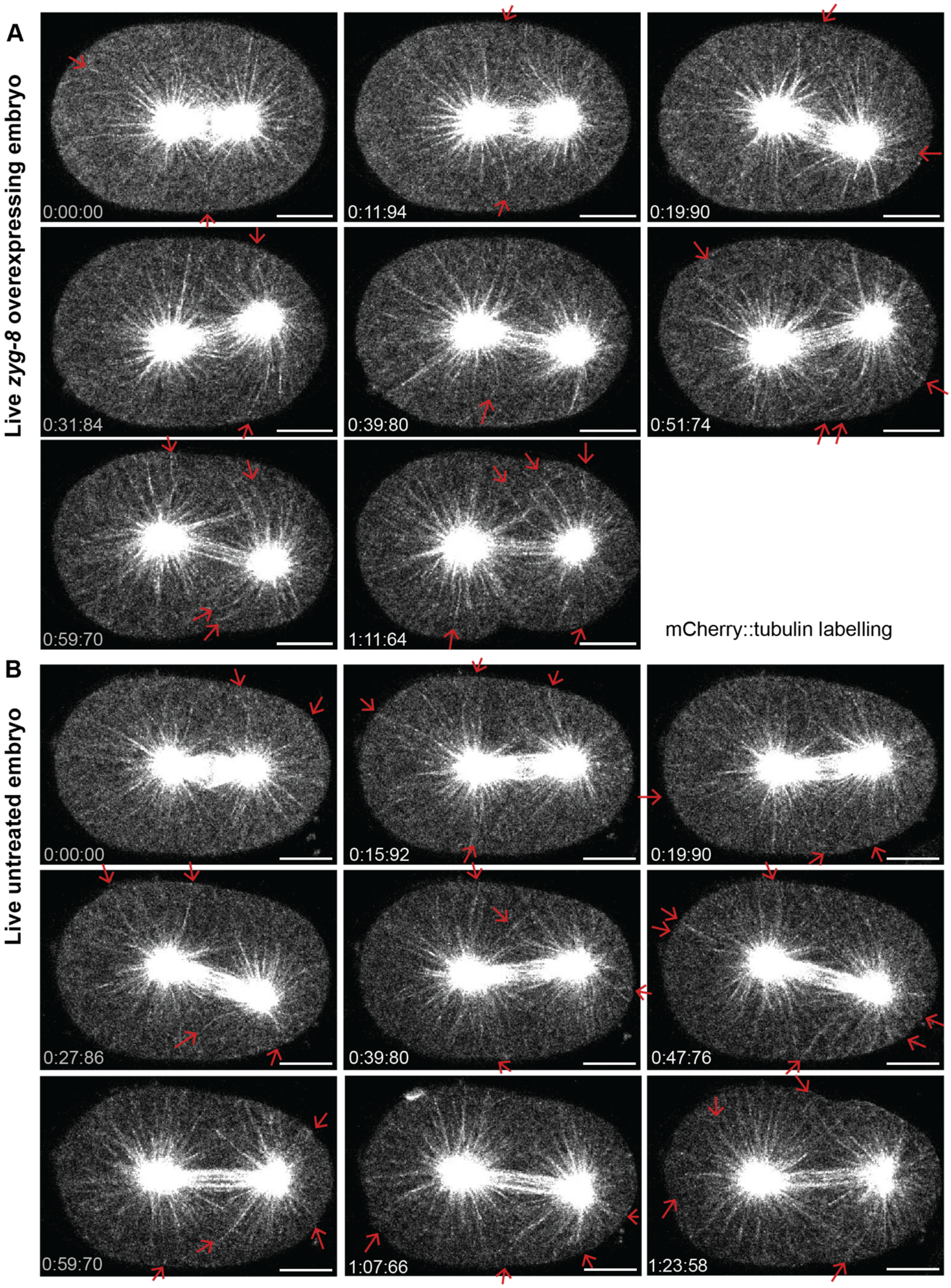


**Figure S10: In embryos overexpressing *zyg-8*, astral microtubules are similarly bent as in untreated embryos during anaphase spindle-pole oscillations.**
Time-lapse confocal super-resolved images along anaphase of (**A**) an exemplar embryo overexpressing *zyg-8* and (**B**) an exemplar untreated embryo. Microtubules were labelled by mCherry::tubulin. Time is indicated from first image in mm:ss:ms. Scale bars represent 10 µm.

Microtubule stiffening by ZYG-8 contributes to spindle orientation.

These time-lapse images are sourced from the movies S10 and S11. Red arrows highlight bent microtubules. Similar images for *zyg-8(or484ts)* mutant and *zyg-8(RNAi)* are shown in Figures 5 and S9, with associated movies provided as Movies S5-S9.

Microtubule stiffening by ZYG-8 contributes to spindle orientation.

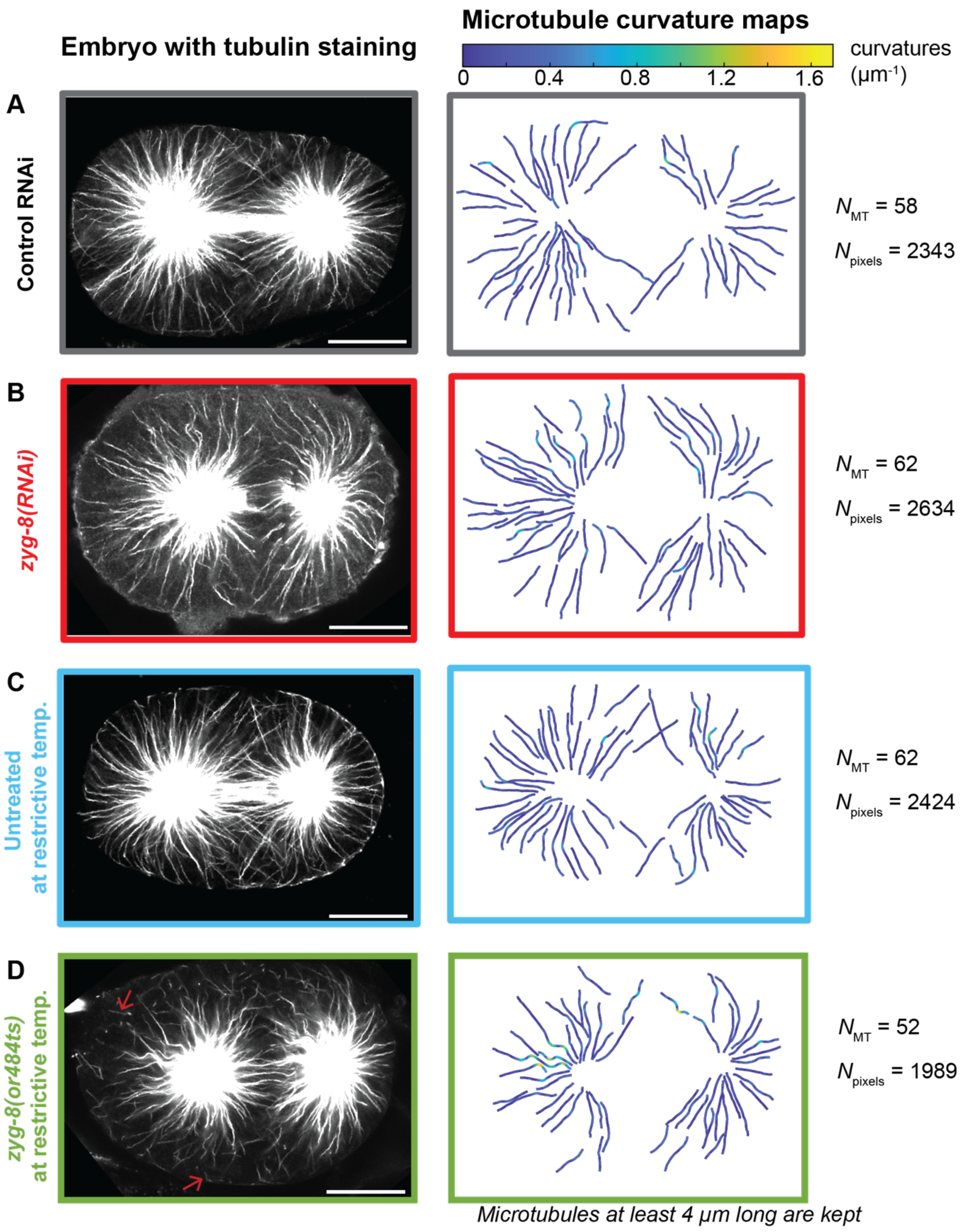


**Figure S11: Image processing pipeline mapping the curvatures of astral microtubules.**
The deconvolved confocal images of fixed embryos with $\alpha$-tubulin immunostaining reveal microtubule shapes during anaphase: (left) the microscopy image, and (right) its curvature map for the four conditions investigated in Figure 6. (**A**) control RNAi embryo, (**B**) *zyg-8(RNAi)*-treated embryo, (**C**) untreated embryo at the restrictive temperature, and (**D**) *zyg-8(or484ts)* mutant at the restrictive temperature (Methods M3 and M4). Scale bars represent 10 µm. Red arrows indicate fragmented

microtubules in the *zyg-8(or484ts)* mutant. For each condition are indicated the number of microtubules analysed ($N_{MT}$) and the total count of pixels ($N_{pixels}$).

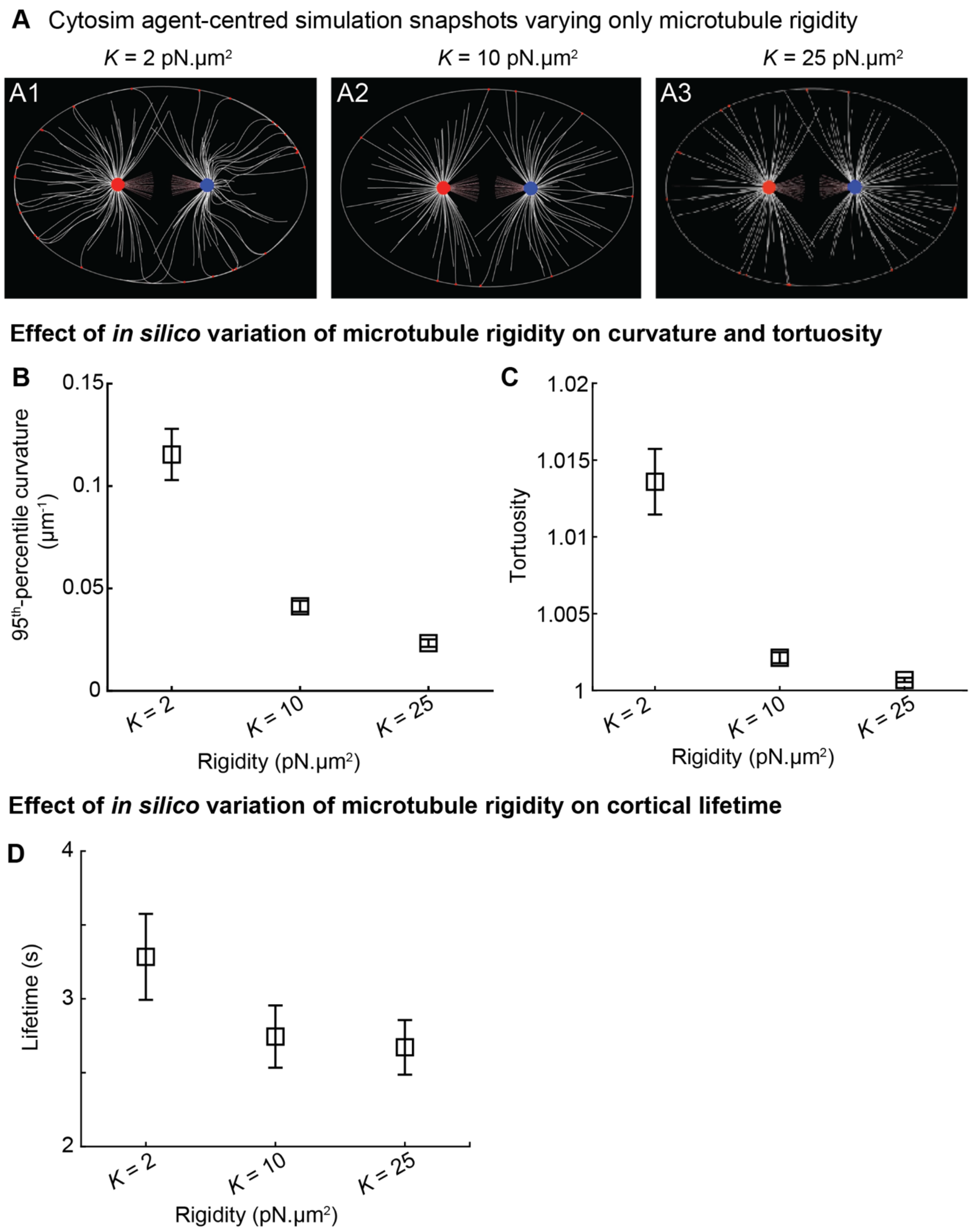


**Figure S12: *In silico,* decreasing microtubule flexural rigidity increases cortical lifetimes.**
(**A**) Snapshots from *Cytosim* agent-centred simulations showing fibres mimicking microtubules in white, with cortical contacts indicated by red dots (Table S2, Method M12). Simulations were performed using three different microtubule flexural rigidity values: $K$ = 2, 10 and 25 pN.µm$^2$. The frames were taken 45 seconds after the start of each simulation, corresponding to movies S12, S13 and S14. (**B-C**) Parameters of astral microtubules extracted from simulations varying microtubule rigidity: (B) medians of the 95$^{th}$ percentile curvatures and (C2) medians of tortuosity values were calculated per simulation. These measurements were taken from the set of microtubules present in the frame recorded 45 seconds after simulation start. (**D**) Cortical lifetimes of astral microtubules were

calculated per simulation, using data collected from 15 seconds onward to allow for system warm-up. Black squares represent the mean, and error bars indicate the standard deviation across 40 simulations per condition.

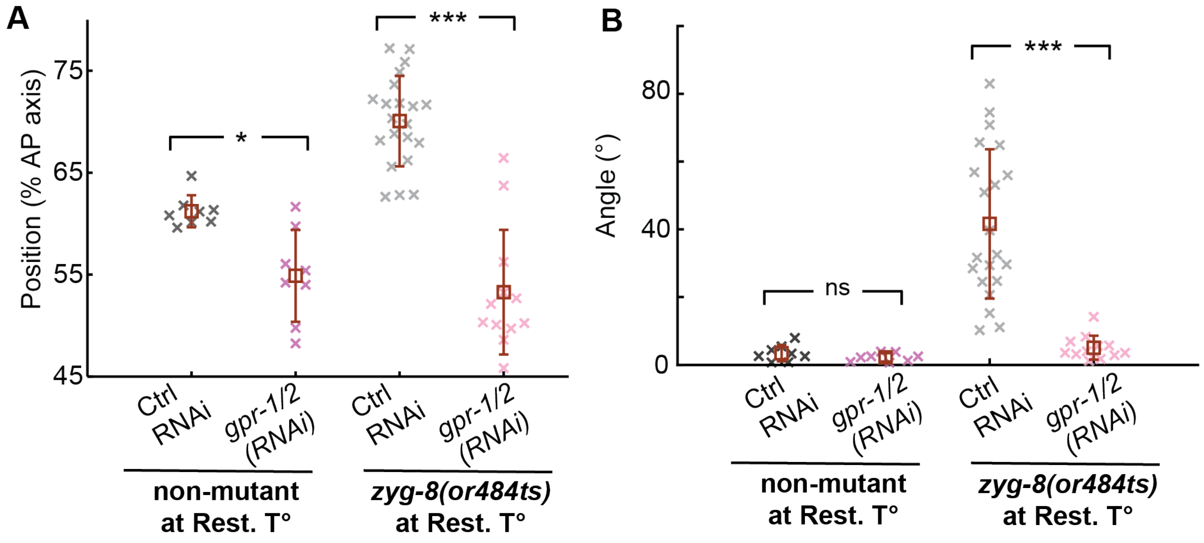


**Figure S13: Reducing cortical pulling forces rescues mitotic spindle misorientation at anaphase end.** (**A**) Most-posterior position of the mitotic spindle during anaphase and (**B**) mitotic spindle angle at 150 s after anaphase onset for the following conditions, all at the restrictive temperature (Rest. T°): (purple) $N$ = 8 *gpr-1/2(RNAi)*-treated non-mutant embryos, (black) $N$ = 8 control RNAi non-mutant embryos, (pink) $N$ = 21 *zyg-8(or484ts)* mutants treated with *gpr-1/2(RNAi)* and (grey) $N$ = 12 *zyg-8(or484ts)* mutants treated with control RNAi. We tracked and analysed the centrosomes of GFP::TBB-2 labelled embryos (Method M7). $N$ represents the total number of embryos analysed across all replicates. The brown squares and error bars correspond to the means and SD. * and *** indicate significant differences with $1 \times 10^{-3} < p \leq 1 \times 10^{-2}$ and $p \leq 1 \times 10^{-4}$, respectively. ns denotes non-significant differences ($p > 0.05$) (Method M15).

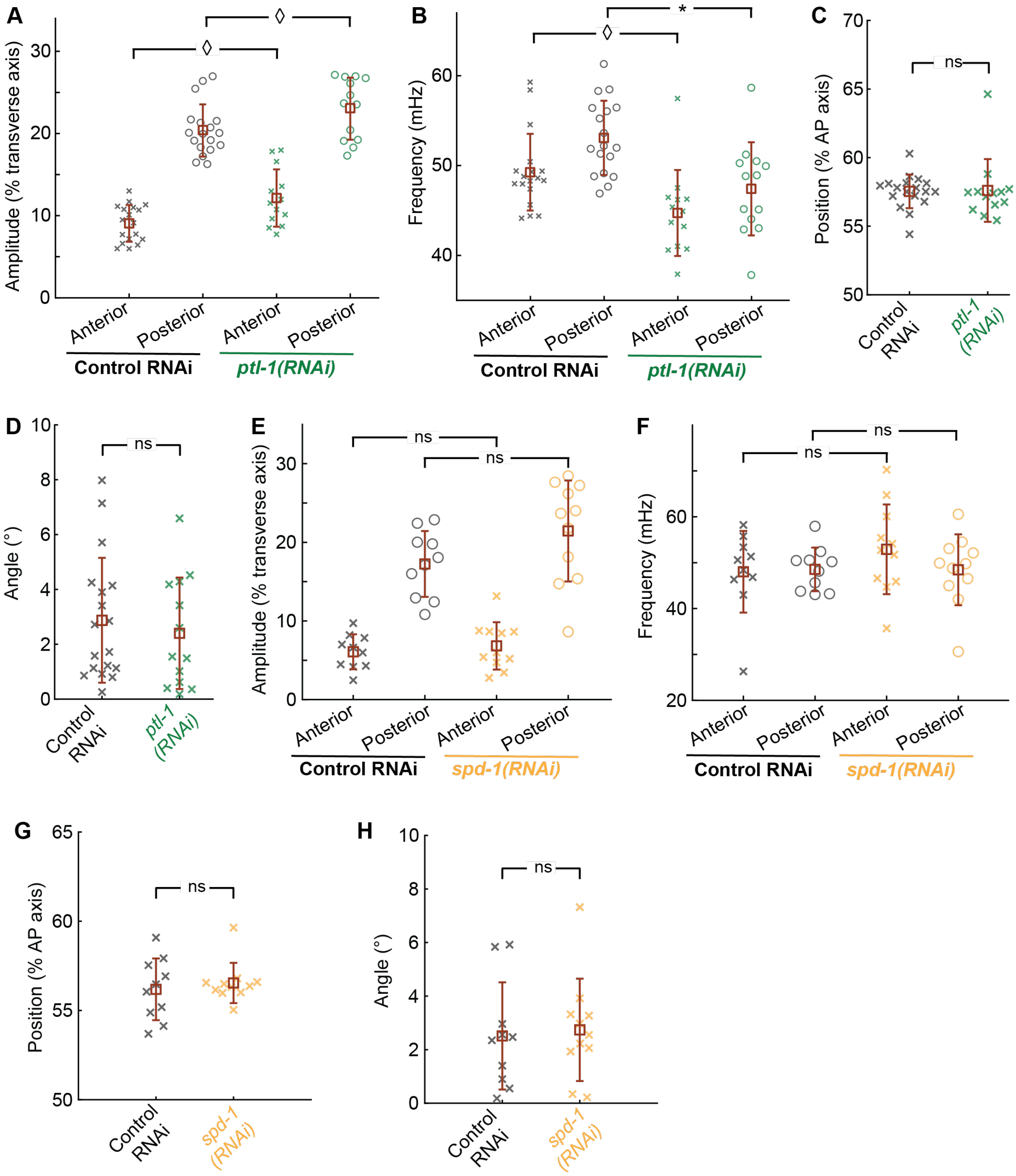


**Figure S14: PTL-1 or SPD-1 does not play any major role in spindle-pole oscillations, spindle final position or orientation.**

(**A**) Maximal amplitudes of spindle pole oscillations and (**B**) frequencies of the oscillations for (green) $N = 13$ *ptl-1(RNAi)*-treated embryos and (black) $N = 18$ control RNAi embryos. We tracked and analysed the centrosomes labelled by mCherry::TBG-1 (Method M7). Spindle final (**C**) positions along the AP axis and (**D**) angles. Embryos are the same as for panels A-B. (**E**) Maximal amplitudes of spindle pole oscillations and (**F**) frequencies of the oscillations for (green) $N = 11$ *spd-1(RNAi)*-treated embryos and (black) $N = 10$ control RNAi embryos. We tracked and analysed the centrosomes labelled by GFP::TBG-1 (Method M7). Spindle final (**G**) positions along the AP axis and (**H**) angles. Embryos are the same as for panels E-F. $N$ represents the total number of embryos analysed across all replicates. The brown squares and error bars correspond to the means and SD. ◊ and * indicate significant differences with

$1 \times 10^{-2} < p \leq 5 \times 10^{-2}$ and $1 \times 10^{-3} < p \leq 1 \times 10^{-2}$, respectively. ns denotes non-significant differences ($p > 0.05$) (Method M15).

| | **Genotype description** | **If crossing, from which strains** | **Origin & Reference** |
|---|---|---|---|
| MCP347 4x* | zyg-8(bab347[3xOLLAS::zyg-8]) III | | This study |
| MCP361 4x* | zyg-8(bab361[mNG::zyg-8]) III | | This study |
| MCP554 2x** | ttTi5605( bab516; bab554[pPie-1::mNG::3*OLLAS::zyg-8::3'UTR tbb-2] II) | | This study |
| TH430 | ddIs14[pie-1p::EBP-2::GFP; unc-119(+)]; weIs21[Ppie-1::mCherry::tub::pie-1 3'UTR] | TH66 crossed with JA1559 | TH66: (Srayko et al., 2005)<br>JA1559: Ahringer lab |
| JEP153 | weIs21[Ppie-1::mCherry::tub::pie-1 3'UTR] | Out-crossing of TH430 | TH430: Hyman lab. |
| JEP170 | zyg-8(bab361[mNG::zyg-8]) III; weIs21[Ppie-1::mCherry::tub::pie-1 3'UTR] | JEP153 crossed with MCP361 4x | This study |
| JEP187 | ttTi5605(bab516; bab554[pPie-1::mNG::3*OLLAS::zyg-8::3'UTR tbb-2])II; weIs21[Ppie-1::mCherry::tub::pie-1 3?UTR] | JEP153 crossed with MCP554 2x | This study |
| JEP97 | unc-119(ed3) ? III; ltIs37 [pAA64; pie-1/mCHERRY::his-58; unc-119 (+)]IV;ddIs44[WRM0614cB02 GLCherry::tbg-1;unc-119(+)] | OD56 crossed with TH169 | OD56: (McNally et al., 2006)<br>TH169: (Woodruff et al., 2015) |
| JEP188 | t tTi5605(bab516; bab554[pPie-1::mNG::3*OLLAS::zyg-8::3'UTR tbb-2])II; unc-119(ed3) ? III; ltIs37 [pAA64; pie-1/mCHERRY::his-58; unc-119 (+)]IV;ddIs44[WRM0614cB02 GLCherry::tbg-1;unc-119(+)] | JEP97 crossed with MCP554 | This study |
| EU3068 | ebp-2(or1954[ebp-2::mKate2]) II; ruIs57[pie-1::GFP::tbb-2 +unc-119(+)] V | | (Sugioka et al., 2018) |
| JEP180 | zyg-8(or484) III; ebp-2(or1954[ebp-2::mKate2]) II | EU3068 crossed with EU924 | EU924: (Encalada et al., 2000) |
| JEP10 | such-1(h1960) III; unc-46(e177) mdf-1(gk2) V. ddIs180[WRM062cF05 spd-2:: 2xTY1 GFP FRT 3xFlag;unc-119(+)] | KR4012 crossed with TH231 | KR4012: (Tarailo et al., 2007)<br>TH231: (Decker et al., 2011) |
| AZ244 | unc-119(ed3) III; ruIs57[pie-1::GFP::tbb-2 +unc-119(+)] V | | (Praitis et al., 2001) |
| JEP190 | zyg-8(or484) III; unc-119(ed3) III; ruIs57[pie-1::GFP::tbb-2 +unc-119(+)] V | AZ244 crossed with EU924 | This study |
| JEP82 | ebp-2(or1954[ebp-2::mKate2]) II | Out-crossing of EU3068 | This study |

| | | | |
|---|---|---|---|
| JEP193 | ttTi5605(bab516; bab554[pPie-1::mNG::3*OLLAS::zyg-8::3'UTR tbb-2] II); ebp-2(or1954[ebp-2::mKate2]) II | JEP82 crossed with MCP554 | This study |
| TH30 | unc-119(ed3) ruIs32[pAZ132: pie-1/GFP/histoneH2B] III; ddIs6 [GFP::tbg-1; unc-119(+)] V | | (Quintin et al., 2003) |
| TH65 | unc-119(ed3); ddIs15 [pPIE-1::YFP::tba-2(genomic);unc-119(+)] | | (Schlaitz et al., 2007) |
| TH27 | unc-119(ed3) III; ddIs6 [Ppie-1::GFP::tbg-1; unc-119(+)] V | | (Oegema et al., 2001) |
| SV1803 | dhc-1(he264[eGFP::dhc-1]) I | | (Schmidt et al., 2017) |
| JEP199 | zyg-8(or484) III; ebp-2(or1954[ebp-2::mKate2]) II; dhc-1(he264[eGFP::dhc-1]) I. | SV1803 crossed with JEP180 | This study |

**Table S1:** List of the strains used in this study.
(* 4-times backcrossed; ** 2-times backcrossed.)

| **Object type** | **Characteristic parameters** | **Reference** |
|---|---|---|
| Space: ellipse | Radii: 24.5 μm and 16.5 μm<br>Viscosity: 5 Pa.s | (Daniels et al., 2006) |
| 2 solids (that mimic anterior and posterior centrosomes) | External force: Anterior: 180 pN; posterior: 300 pN | (Grill et al., 2003) |
| 2 asters with two fibre types | Initial position: Anterior: -5.6 μm; Posterior 4.7 μm from ellipse centre. Positions not fixed during the simulation. | *In vivo* measurements of the lab |
| Fibre, type #1 (spindle) | Activity: static<br>Number per aster: 20<br>Initial length 6 μm<br>Rigidity: 100 pN.μm$^2$<br>Position: 60°, radial distribution | |
| Fibre, type #2 (astral) | Activity: dynamic<br>Number per aster: 75<br>Initial length 8 $\pm$ 6 μm<br>Rigidity: 5, 10, 25 pN.μm$^2$<br>Position: 240°, random radial distribution<br>Growing force: 1.67 pN<br>Minimal length: 0.005 μm<br>Growing speed: 0.71 μm/s<br>Shrinking sped: -0.84 μm/s<br>Catastrophe rate: 0.05 (no force); 0.5 (stall force)<br>Rescue rate: 0.15 | Dynamics: (Dogterom and Yurke, 1997; Srayko et al., 2005)<br>Rigidity: (Felgner et al., 1996; Gittes et al., 1993; Kikumoto et al., 2006; Mickey and Howard, 1995; Venier et al., 1994) |
| Single with hands | Activity: bind<br>Anchored to a fixed position<br>Unbinding rate: 0.1<br>Unbinding force: 5 pN | |

**Table S2:** Objects and their parameters used in the three *Cytosim* simulations (Method M12).

Microtubule stiffening by ZYG-8 contributes to spindle orientation.

| | **Control RNAi (*N* = 11)** | | ***zyg-8(RNAi)* (*N* = 14)** | |
|---|---|---|---|---|
| | Population #1 | Population #2 | Population #1 | Population #2 |
| **Lifetime (s)** | 0.66 ± 0.04 | 1.66 ± 0.25 | 0.74 ± 0.03 | 2.07 ± 0.24 |
| **Proportion (%)** | 86.3 ± 2.5 | 13.7 | 84.3 ± 6.7 | 15.7 |
| **Density (/min/µm²)** | 0.79 ± 0.02 | 0.13 ± 0.01 | 0.95 ± 0.04 | 0.18 ± 0.01 |

**Table S3: Changes in cortical dynein density and lifetime upon ZYG-8 depletion phenocopy variations in cortical astral microtubule dynamics.**
Characteristics of the two cortical dynein populations in strains expressing GFP::DHC-1. Data were obtained by fitting the duration histograms of cortical dynein spots using a two-exponential model (Method M13). *N* represents the total number of embryos analysed across all replicates for each experimental condition.

Microtubule stiffening by ZYG-8 contributes to spindle orientation.
